# Neural processing of goal and non-goal-directed movements on the smartphone

**DOI:** 10.1101/2022.08.19.504603

**Authors:** Ruchella Kock, Enea Ceolini, Lysanne Groenewegen, Arko Ghosh

**Author notes:** Corresponding address, Arko Ghosh, Leiden University, Wassenaarseweg 52, Leiden, 2333 AK.

## Abstract

The discrete behavioral events captured on the smartphone touchscreen may help unravel real-world neural processing. We find that neural signals (EEG) surrounding a touchscreen event show peculiarly contralateral motor preparation, visual processing, and the consolidation of information. We leveraged these events in conjunction with kinematic recordings of the thumb and an artificial neural network to separate highly similar movements according to whether they resulted in a smartphone touch (goal-directed) or not (non-goal-directed). Despite their kinematic similarity underscored by the model, the signatures of neural control of movement and the post-movement processing were substantially dampened for the non-goal-directed movements, and these movements uniquely evoked error-related signals. We speculate that these unnecessary movements are common in the real world and although inconsequential the brain still provides limited motor preparation and tracks the action outcome. Real-world behavior is composed of neural processes that are difficult to capture in conventional laboratory-based tasks.

## Introduction

How the human brain generates real-world behavior is sparsely understood. This is partly because artificial behaviors – disconnected from daily life and highly instructed – dominate the study of neuro-behavioral correlates, and how to use what has been learned using such paradigms to understand the real-world behavioral outputs is not clear. Behaviors such as the reaction time task and the less instructed voluntary finger movements have been instrumental in isolating specific neural processes for the neural control of movement and sensory processing. Still, in the real world, multiple neural processes may be simultaneously engaged and the statistical properties of the tasks are fundamentally distinct from what is experienced in the real world (1, 2). There is a fast emerging understanding of the complex naturalistic statistics of sounds, images, and movements, and this recent paradigm shift has already helped unravel specific neural processes tuned to naturalistic information and movements (3, 4). Smartphones are ubiquitous in modern human behavior and they are a source of complex visual information largely driven by touchscreen interactions. Studying the brain activity underlying this behavior may help discover how a range of neural processes is orchestrated to generate behavior that is truly meaningful to daily life.

We can simply time-lock the EEG signals to the touchscreen interaction events captured at a millisecond resolution (5). This allows us to leverage the conventional event-related potential framework to recognize the underlying neural processes based on the well-studied signal features. For instance, bilateral negativity over the sensorimotor electrodes and desynchronization of beta oscillations can help infer the processes underlying motor processes (6–8). Furthermore, negativity over the visual electrodes can indicate visual processing and the frontal-to-central wave of positivity can help reveal memory-related information consolidation (6, 9–12).

According to a widely held notion, when engaged in behavior, all of the movements generated are simply linked to the behavioral goal. However, a scattered set of observations challenge this notion by unraveling movements that are not simply related to the set behavioral goal. Firstly, according to subjective self-reports inconsequential movements dubbed *fidgeting* are ubiquitous in the real world (13). Secondly, in the laboratory, there is emerging evidence for the idea that the cortex is less engaged during the task-irrelevant (non-goal-directed) vs. the relevant (goal-directed) actions. According to invasive neural recordings from the frontal eye field of the monkey cortex, the beta-band remains synchronized during the non-goal-directed saccades (14). In humans, when instructed to aimlessly touch the screen, the beta desynchronization and the motor-related potentials are diminished than in the goal-directed touches aimed at a certain location (15). While these studies demonstrate the capacity to generate non-goal-directed actions, if and how they are generated when engaged in real-world behavior remains unclear.

Capturing the non-goal directed neural signals is conceptually more complex than capturing the neural signals surrounding the smartphone touches. And even if they are captured, they may not be simply comparable to the neural signals time-locked to the smartphone interactions. For instance, the difference in the neural signals between smartphone touches and non-goal-directed movements could be attributed to the differences in the peripheral signal features chosen to time-lock the neural signals. Moreover, while a motion sensor attached to the thumb can detect movements with high fidelity, it cannot directly yield decisive temporal landmarks that could be used to study kinematically similar goal and non-goal-directed movements from a series of signal fluctuations – where one movement is followed by another. To circumvent these issues, we trained an artificial neural network to mark smartphone touchscreen interaction timings based on the movement sensor signals. We expected the model to correctly identify smartphone touches (true positives) and anticipated that the kinematically highly similar movements - if they exist - would yield false positives (i.e., movement without a touchscreen touch). As for both the goal and the non-goal-directed movements the temporal landmarks can be based on the same set of learned features this approach offers an opportunity to contrast the time-locked neural signals.

We reveal how neural processes are peculiarly orchestrated surrounding smartphone behavior, by combining data-driven behavioral modeling, smartphone touchscreen interaction logs, and parametric statistics of event-related (spectral) analysis (across all electrodes and broad time range) surrounding the discrete events. Our analysis reveals a stark distinction between the goal and non-goal-directed actions spanning a range of neural processes.

## Results

### The neural signals surrounding smartphone touchscreen events

The event-related brain signals surrounding the smartphone touchscreen events spanned ~2.3 s. The initial signals starting at −700 ms consisted of a slow rise of signals (positivity) over the contralateral to the movement (i.e., left side) sensorimotor electrodes (**Figure 1**, Supplementary Movie 1). This rise was followed by gradual negativity starting at about −300 ms in the same region. This negativity lasted till 200 ms, with the negativity shifting to the occipital electrodes where it was sustained for another 250 ms. This was followed by a frontal to central positivity spanning between 500 to 700 ms. These signals were followed by more topologically scattered activations that finally terminated by ~1.6 s. The initial sensorimotor activations were notably contralateral and in contrast to the initially bilateral sensorimotor activity described in laboratory-designed behaviors. Indeed, we could reproduce bilateral negativity when we instructed participants to touch a smartphone-like surface (Supplementary Movie 2).

**Figure 1.**
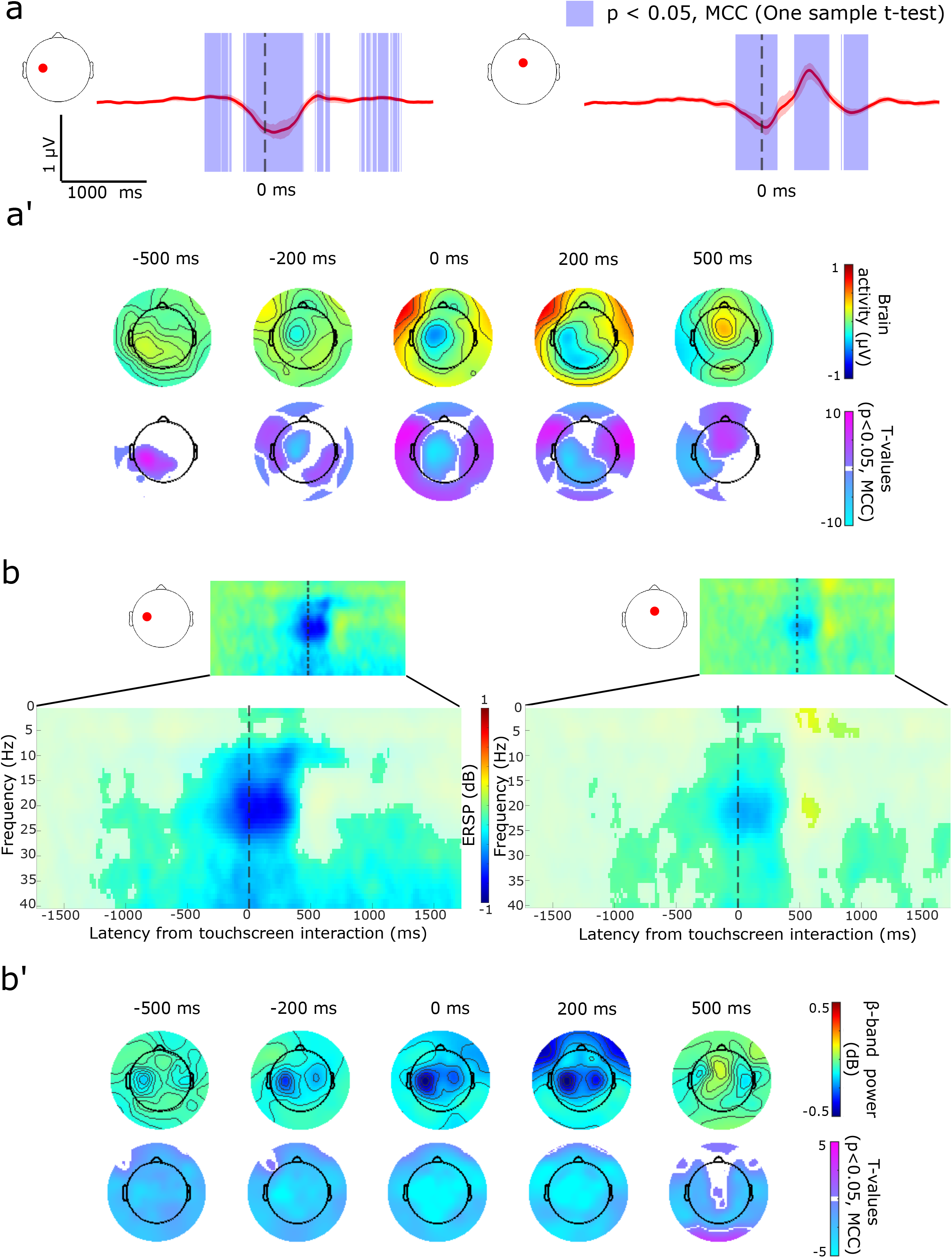
Event-related potential and spectral analysis of EEG signals surrounding smartphone touchscreen interactions. Participants used their right thumb to interact with their smartphone on commonly used apps determined based on usage history. (**a**) Touchscreen interactions show negative deviations at electrodes over the left sensorimotor cortex (left plot, red dot), shown with trimmed means (20%) and 95% confidence intervals. Similar negative deflections occur at a mid-frontal electrode. The data were band-pass filtered between 0.5 to 3 Hz for visualization. Significant statistical clusters were determined by using one-sample *t*-tests and multiple comparison corrections across all electrodes and time points (MCC, p < 0.05, shaded purple in the plot). **(a’)** Scalp topologies of trimmed mean signals and T-values show marginal significant positive activity preceding the touchscreen interaction followed by prominent negativity over the sensorimotor cortex. Positive activity recorded over the central to frontal areas occurs after the interaction (MCC, p < 0.05). (**b**) Prominent event-related spectral desynchronization was observed over the left sensorimotor cortex (left plot, red dot). A similar pattern was observed over mid-frontal electrodes. **(b’)** Scalp topology shows widespread beta-band desynchronization (for visualization, the data is collapsed across the beta-band by estimating the 20% trimmed means at each time point and the significant masked T-values were collapsed by using the maximum absolute amplitude). Significant statistical clusters were determined by using one-sample *t*-tests and multiple comparison correction (MCC, p < 0.05). Approximate times are used for scalp topologies as time information was adjusted due to continuous wavelet transform. For the full statistical outcomes of touchscreen event-related (spectral) potentials see Supplementary Movie 1, Supplementary Movie 3 (one-sample *t*-test, focused on the beta-band, touchscreen interactions), and Supplementary Movie 4 (one-sample *t*-test, focused on the alpha-band). See Supplementary Movie 2 for event-related potentials surrounding touches on a smartphone-like surface.

Time-frequency analysis revealed strong beta-band desynchronization (reduced power) surrounding the touchscreen interactions. The desynchronization over the sensorimotor electrodes appeared ~1.2 s before the interaction and strengthened up to the interaction. The pre-interaction desynchronization appeared asymmetric with lower power over the ipsilateral electrodes. Strong event-related desynchronization persisted after the interaction and the oscillations rebounded by 550 ms over the central electrodes. The desynchronization was not limited to the beta band during the interaction but extended to the alpha and gamma bands (**Figure 1,** Supplementary Movie 3 for the beta band, Supplementary Movie 4 for the alpha band).

### The identification of goal and non-goal-directed smartphone movements

While the EEG signals surrounding the smartphone interactions could be simply studied by time-locking to the touchscreen events, we deployed an artificial neural network to find landmarks on the movement signals corresponding to the touchscreen interaction. The model performed with an F2 score (a measure of model recall and precision, with a higher weight on the recall) of 0.35 (median of all trained subjects, n = 68) and 0.34 (median based on subjects considered here, n = 32) (Supplementary Table 1, Supplementary Figure 1). The identified landmarks that did not coincide with a smartphone interaction – non-goal-directed movements – revealed a degree of kinematic similarity with the goal-directed movements (**Figure 2a’** shows an example participant, see Supplementary Figure 2 for all participants). As we were interested in studying the neural correlates contrasting these two movement types notwithstanding any kinematic differences, we further considered only those subjects with highly similar kinematic fluctuations between the two movement types (R > 0.8, See Supplementary Methods for the distribution of Pearson Rs).

**Figure 2.**
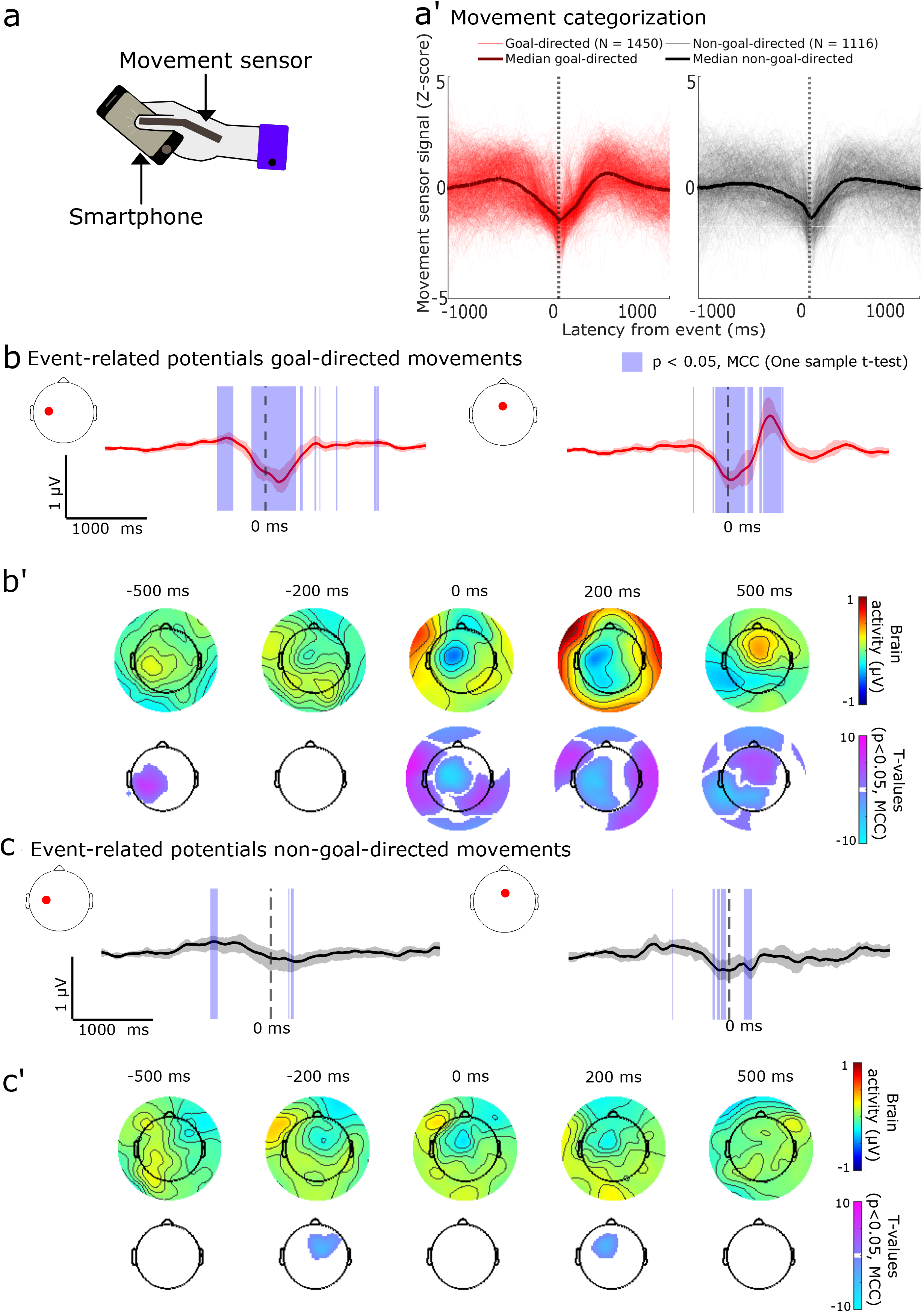
Movement dynamics and event-related potentials of the goal and non-goal-directed movements. **(a)** Movement signal traces for goal (red, left) and non-goal-directed (black, right) movements time-locked to the predicted events for one participant illustrating the high similarity between the movements (median traces overlaid). Movement signals were Z-score normalized for visualization. **(b-b’)** Event-related potential surrounding goal-directed movements show similar activations as touchscreen interactions (one-sample *t*-tests, MCC, p < 0.05). Same legend as for main figure 1a-a’. **(c-c’)** Event-related potential surrounding non-goal-directed movements show small constrained significant deviations over the left sensorimotor cortex and the midline frontal electrode before and after movement onset. Topological plots show statistically significant clusters on the frontal-to-central electrodes unique to the goal-directed movements. For paired *t*-tests comparing goal and non-goal-directed movements see Supplementary Figure 3. For the full statistical outcomes of event-related potentials see Supplementary Movie 5 (one-sample *t*-test goal-directed movement), Supplementary Movie 6 (one-sample *t*-test non-goal-directed movement), and Supplementary Movie 7 (paired *t*-test goal vs. non-goal-directed movements).

The non-goal-directed movements occurred as frequently as the goal-directed movements in the selected population (Supplementary Figure 1b). The inter-event intervals were typically separated by ~1 s for the goal-directed movements whereas the non-goal-directed movements were separated by a broader distribution with a primary peak under ~1 s and a secondary peak at ~10 s (Supplementary Figure 1c). The non-goal-directed movements were more likely to occur right after the goal-directed movement rather than before (*t*_(32,31)_ = −2.6910, p = 0.0114, paired *t*-test, Supplementary Figure 1d).

### EEG potentials surrounding goal-directed vs. non-goal-directed movements

The temporal landmarks deduced by the artificial neural network were used to time-lock the EEG signals for both goal and non-goal-directed movements enabling a fair comparison between the neural activations surrounding the two movement types. Unsurprisingly, the patterns for the goal-directed movements were highly similar to the patterns seen for the touchscreen interactions as described above (**Figure 2**). Interestingly, there were some notable differences in the event-related potentials between the two movement types.

With regards to the initial sensorimotor signal features, both movement types displayed a slow rising positivity over the contralateral electrodes before the temporal landmarks. However, the subsequent negativity was substantially dampened for the non-goal-directed movements, as they failed to yield statistically significant clusters over the contralateral electrodes (**Figure 2**, for full results, see Supplementary Movie 5 – goal-directed movements, Supplementary Movie 6 – for non-goal-directed movements, for paired *t*-test, see Supplementary Figure 3, Supplementary Movie 7, for model output distributions underlying these analysis see Supplementary Figure 4). Interestingly, we discovered a statistically significant cluster corresponding to a negative signal over the ipsilateral sensorimotor cortex before the temporal landmark of the non-goal-directed movements (See between −200 ms to −50 ms, in Supplementary Movie 6, see **Figure 2** for snapshot). The differences between the movements were even more striking after the temporal landmarks. While the goal-directed movements displayed a rich array of activations spanning various regions, there was only a constrained cluster detected over the frontal electrodes. The negativity over the fronto-central electrodes was unique to the non-goal-directed movements (and persisted between ~200 – 280 ms).

### Desynchronization surrounding the goal-directed movements is dampened for the non-goal-directed movements

We next analyzed the oscillatory hallmarks well implicated in the cortical control of movements. Time locking to the model predicted temporal landmarks confirmed a striking desynchronization (suppressed power) of the alpha, beta, and gamma oscillations for the goal-directed movements, which was similarly observed surrounding the smartphone interactions (**Figure 3**, Supplementary Figure 5, alpha - Supplementary Movie 8, beta - Supplementary Movie 9, gamma - Supplementary Movie 10). Only a marginal desynchronization of the beta oscillations was observed in the case of non-goal-directed movements predominantly over the contralateral sensorimotor areas (**Figure 3,** Supplementary Movie 11). This desynchronization was temporally constrained between ~350 ms preceding the movement to ~450 ms after the event. The gamma and alpha oscillations showed similar desynchronizations, but with the gamma signal being more prolonged (for topology see Supplementary Figure 5, gamma - Supplementary Movie 12, alpha - Supplementary Movie 13). A paired *t*-test established that the beta desynchronization was marginal in contrast to the goal-directed movements, and revealed statistically significant clusters mostly over the contralateral hemisphere spanning ~100 ms before the event to ~400 ms after the event (Supplementary Figure 3, Supplementary Movie 14).

**Figure 3.**
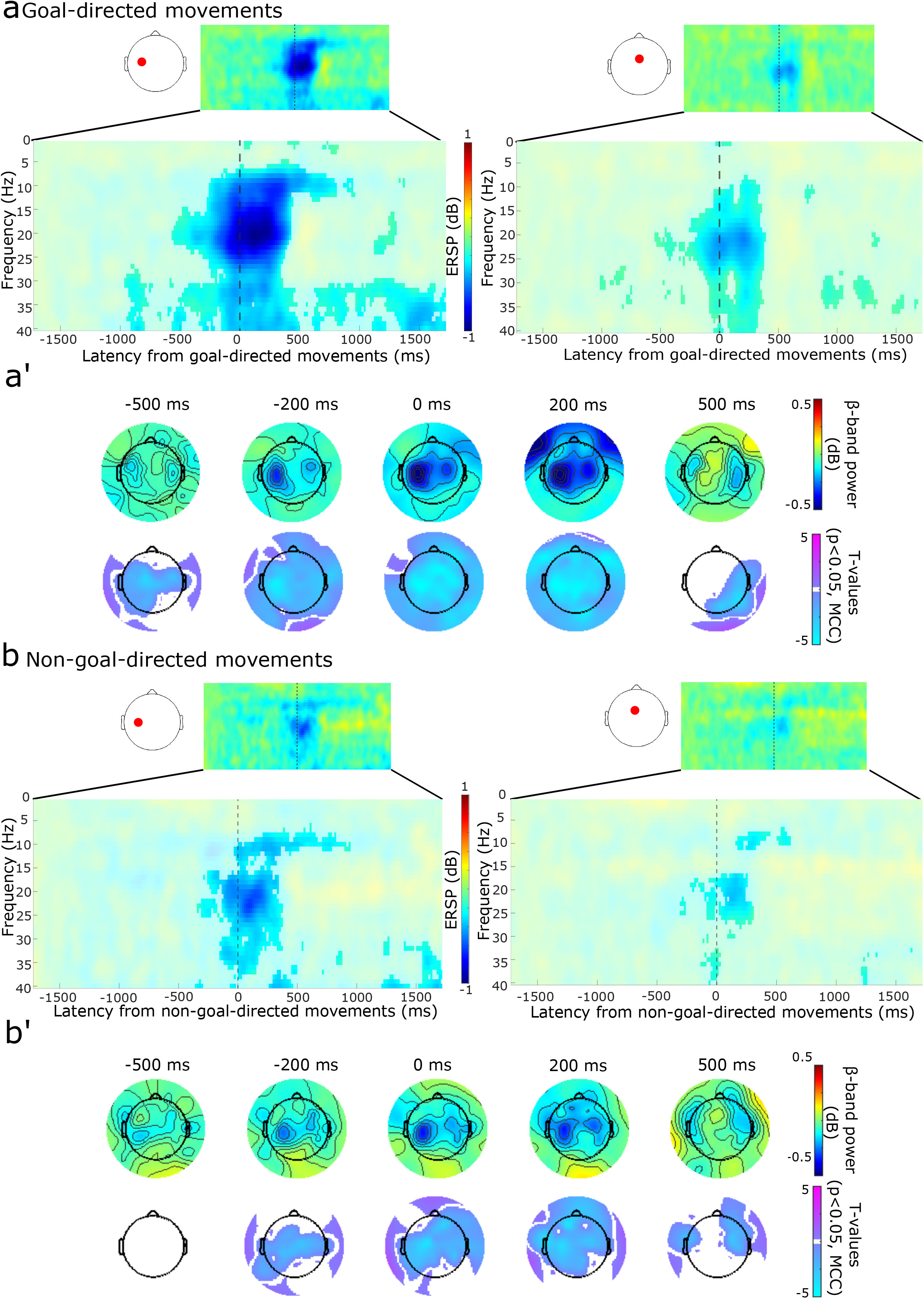
Event-related spectral analysis for goal and non-goal-directed movements. **(a-a’)** Goal-directed movements. Desynchronization was recorded over the left sensorimotor cortex and mid-frontal electrodes. Same legend as in figure 1b-b’. **(b)** Surrounding non-goal-directed movements, highly constrained beta and gamma-band desynchronization were recorded at the electrodes over the left sensorimotor cortex and midline frontal areas. **(b’)** The beta-band desynchronization was spatially and temporally constrained. For alpha-band topologies see Supplementary Figure 5. For paired sample *t*-tests see Supplementary Figure 3. For full statistical outcomes of the event-related spectral analysis see Supplementary Movie 8 (one-sample *t*-test, alpha-band, goal-directed movements), Supplementary Movie 9 (one-sample *t*-test, beta-band, goal-directed movements), Supplementary Movie 10 (one-sample *t*-test, gamma-band, goal-directed movements), Supplementary Movie 11 (one-sample *t*-test, beta-band, non-goal-directed movements), Supplementary Movie 12 (one-sample *t*-test, gamma-band, non-goal-directed movements), Supplementary Movie 13 (one-sample *t*-test, alpha-band, non-goal-directed movements), Supplementary Movie 14 (paired *t*-test goal-directed vs. non-goal-directed movements).

### Sensorimotor cortical response to tactile stimulation during non-goal directed movements

The dampened cortical signals during non-goal-directed movements, in contrast to the prominent cortical signals observed during the goal-directed movements, raise the possibility that sensorimotor cortical information processing of sensory inputs from the thumb is suppressed during the non-goal-directed movements. Alternatively, the dampening may be specific to movements and the cortex may continue to process sensory inputs from the hand. To test these ideas, we analyzed the voltage signals stemming from artificial tactile stimulations to the thumb tip coinciding with goal-directed movements (occurring within ± 500 ms) in contrast to those stimulations coinciding with non-goal-directed movements. Notably, the tactile stimulations resulted in strong event-related signals in both conditions over the contralateral sensorimotor cortex (**Figure 4,** goal-directed artificial touches - Supplementary Movie 15, non-goal-directed artificial touches - Supplementary Movie 16). In sum, although signals are dampened during non-goal-directed movements the sensorimotor cortex remained available for tactile information processing.

**Figure 4.**
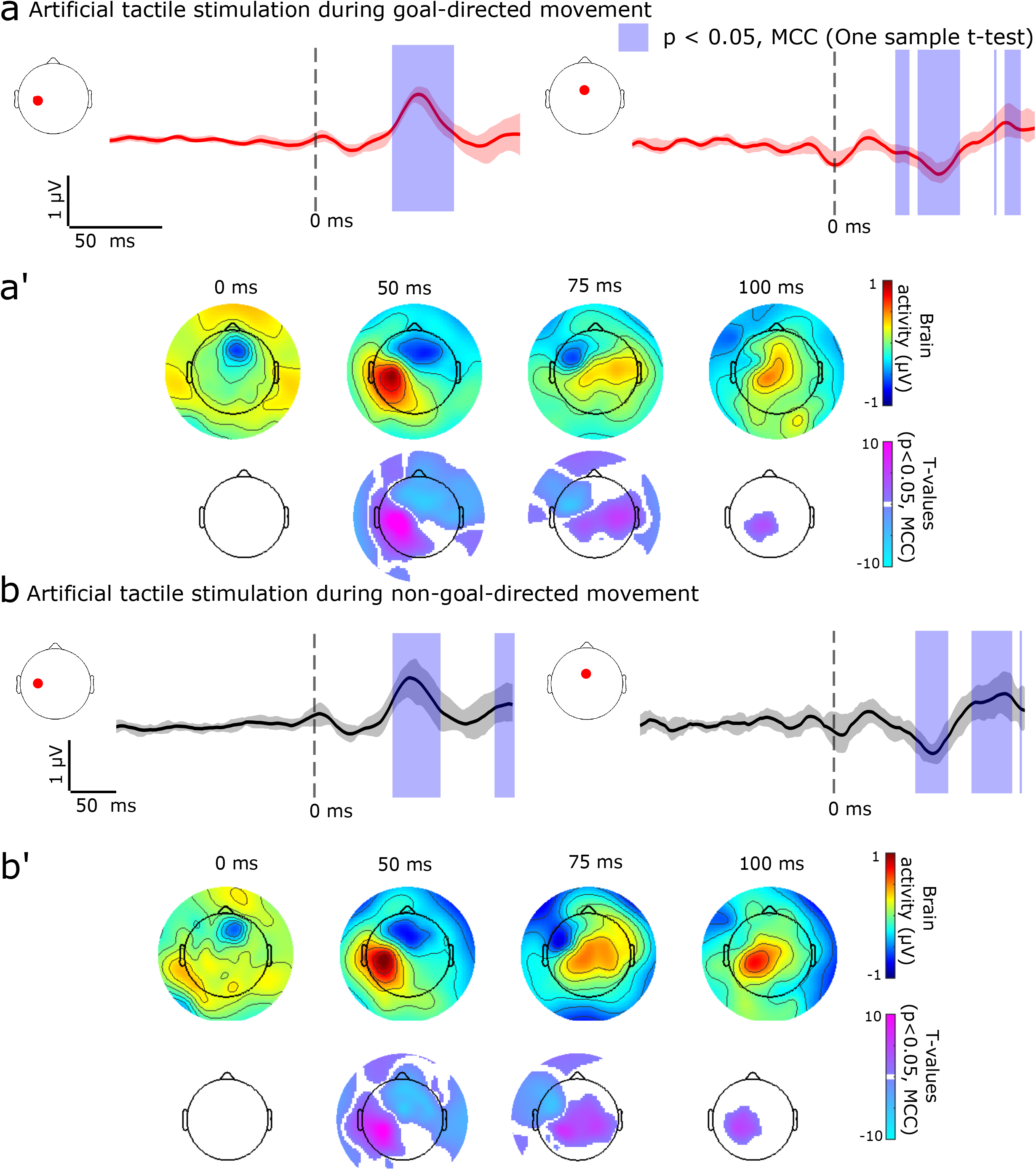
Event-related potentials for artificial tactile stimulations coinciding with the goal and non-goal-directed movements. **(a)** Statistically significant clusters (denoted with purple overlay) involving a positive component were observed after the artificial tactile stimulation during goal-directed movement over the left sensorimotor cortex (left plot, red dot). A negative component was observed at the mid-frontal electrode. **(a’)** Scalp topologies show statistically significant clusters over the sensorimotor cortex. (**b**-**b’**) A near-identical pattern of signals was visible when the stimulations coincided with the non-goal-directed movement. Significant statistical clusters were identified using one-sample *t-*tests and multiple comparisons corrected (MCC, p < 0.05). For full statistical outcomes see Supplementary Movie 15 (one-sample *t*-test, goal-directed movements coinciding with artificial touches), Supplementary Movie 16 (one-sample *t*-test, non-goal-directed movements coinciding with artificial touches)

## Discussion

Through time-locking the EEG signals to discrete temporal landmarks associated with smartphone behavior we identified a range of neural processes, and some of these processes were peculiar to smartphone use. Strikingly, when engaged on the smartphone, not all of the generated movements resulted in touchscreen interactions. These non-goal-directed movements were processed differently by the brain as opposed to the goal-directed movements. Our findings provide a comprehensive overview of how the brain engages in smartphone interactions and highlights the importance of studying real-world behaviors to discover novel neural processes.

The neural signals time-locked to the smartphone touchscreen events revealed peculiar patterns of activity that may not be observed in common laboratory paradigms. First, the events were proceeded by a slow build-up of positivity over the contralateral sensorimotor electrodes. In conventional paradigms a similar positivity – albeit over central electrodes and inconsistently observed – correlates with voluntary action inhibition (16, 17). Our findings where this putative inhibitory signature is followed by negativity raise the possibility that motor outputs emerge from a competitive process that requires overcoming underlying neural inhibition (18). Second, motor preparation of the conventional artificial tasks – from reaction time to instructed voluntary key presses – show bilateral negativity over the sensorimotor electrodes (and this was confirmed here in a subset of the participants generating smartphone-like voluntary thumb movements upon instruction), whereas only a contralateral (to movement) negativity was found surrounding the smartphone touch (11, 19). This ipsilateral disengagement for motor control could stem from the day-to-day repetition of smartphone interactions resulting in a highly optimized circuitry negating inter-hemispheric interactions for motor control, or may reflect top-down processes that suppress ipsilateral activity to promote motor learning (20, 21). Either way, the ipsilateral disengagement must be highly context-dependent as, when performing an artificial task, the ipsilateral hemisphere was vividly engaged for the same type of thumb movements.

The touchscreen events did evoke some familiar neural signals associated with visual processing – i.e., bilateral negativity over the occipital electrodes and a subsequent (~250 ms after the visual response) frontal-to-central positive wave. While the former probably indicates the visual processing of the new content on the screen triggered by the touch, the latter may indicate subsequent information consolidation involving memory processes (22). According to the time-frequency analysis, the touchscreen events were surrounded by robust beta and alpha-band desynchronizations. These desynchronizations are commonly reported for voluntary movements and indicate increased excitability in the populations engaged in movement-related sensorimotor processing (23). A rebound of beta-oscillations was observed at ~500 ms after the touchscreen event, which may reflect an inhibited motor cortical state and signal action completion to other brain areas (24).

Our findings suggest that much of the neural processing during smartphone behavior is dedicated to non-goal-directed movements, and such movements have been long ignored as conventional tasks fixate on highly instructed movements. Still, the familiar signal features – based on artificial tasks – provided some hints on the neural underpinnings of these unnecessary movements. The movements were associated with a rising positivity over the contralateral sensorimotor electrodes – suggesting that the speculative action inhibition processes must be overcome for non-goal-directed movements as well. The subsequent negativity over the contralateral sensorimotor electrodes, and the beta-band desynchronization, were substantially diminished for the non-goal-directed movements compared to the goal-directed movements. These findings on the beta-band oscillations are akin to the correlates of non-goal-directed eye movements recently captured in non-human primates and humans aimlessly performing finger movements (14, 15). As suggested in non-human primates based on invasive neural recordings, these diminished signals may either stem from greater trial-to-trial neural variability or due to more limited recruitment of neural populations (14). We found a negative signal over the ipsilateral sensorimotor electrodes for the non-goal-directed movements. Perhaps these activations are related to inter-hemispheric interactions needed to absorb the broad consequences of the non-goal-directed movements.

The activation at the visual electrodes and the consolidation-oriented signals were absent for the non-goal-directed movements. It remains unclear if the brain pre-empts the inconsequential nature of the non-goal-directed movements at the time of motor preparation to suppress visual processing, or if the absent visual processing can be explained by the lack of a trigger in the form of a tactile event or visual content change. The brain may even keep track of the outcomes of the non-goal directed movements, and signal these erroneous occurrences to the downstream processes to impact visual processing. Indeed, we discovered frontal negativity (mimicking the error-related negativity) following the non-goal-directed movements but this was not observed for the goal-directed movements (25, 26).

While our results indicate that distinct neural processes underly non-goal-directed actions, why they occur at all is not clear. These movements may be a by-product of the cortical-sub-cortical interactions. For instance, subcortical structures may help prepare multiple actions in parallel, and some of these may be released as non-goal-directed movements (27). Furthermore, the role of the cortex in these movements is also not clear. The dampened signals could stem from the engagement of deep, scattered, or highly variable neural signal sources. Resolving the sources of the non-goal-directed movements offers exciting avenues for future research. For instance, using resolved sources we can address if the different movements involve overlapping neural populations or if these distinct movements originate from distinct neural computations. Furthermore, it would become possible to address how the motor and sensory areas orchestrate such that the non-goal-directed movements do not interfere with processing the smartphone information. The neural signals time-locked to the smartphone events in themselves provide a new and highly accessible way to study how various neural processes combine to enable real-world behavior. We anticipate this will enable real-world focused research well beyond fundamental cognitive science, for instance to discover markers of neurological dysfunctions. In conclusion, a combination of data-driven behavioral models in conjunction with neural recordings, and prior research using artificial tasks, make the complex neural signals underlying real-world behavior interpretable and possibly broadly useful.

## Methods

### Participants

Healthy right-handed volunteers (self-declared) were recruited by using on-campus advertisements for a large study deploying multiple sensors to improve the fundamental understanding of smartphone behavior. From this recruitment drive, 106 subjects participated in measurements containing the sensor data required for this study (56 females) from 18 to 46 (median age of 24). All participants provided written and informed consent, and the study was approved by the Institute of Psychology Ethics Committee at Leiden University. For an overview of participants and the different stages of technical elimination please see Supplementary Methods.

### Instructed movements on a smartphone-like box

To capture the common movement-related EEG signals associated with artificial tasks, we instructed participants (twenty-seven of the recruited subjects) to touch a dummy smartphone-like box, attached to a force sensor (Interlink Electronics, Camarillo). They were instructed to touch a target on the box whenever they felt like in 5 seconds (with a 5-second clock visible to the user). The force sensor signals served as a representation of the touchscreen interactions. The touches on the force sensor did not yield any digital feedback, and only touches that occurred within the 2 mm target perimeter were recorded. The force sensor output was gathered using a USB 6008 DAQ (National Instruments, Austin).

### Smartphone data collection

Participants installed the TapCounter app (QuantActions Ltd, Lausanne) before the laboratory visit. The app operated in the background and gathered the timestamps of each interaction with a millisecond resolution, and with a typical error of 0 ms (5). Participants were instructed to use the two most used social and two non-social apps, based on their behavior gathered before the laboratory measure (21) (See Supplementary Methods for an overview of the apps used according to the frequency of goal and non-goal-directed movements). Participants were instructed to only use their right thumb during the laboratory measure and this was further verified online by using video recordings (For an illustrative video of the experimental setup see Supplementary Movie 17). The usage sessions lasted for ~ 1 h and participants were encouraged to take short (< 1 min) breaks every 10 minutes.

### Artificial tactile stimulation

Compact solenoid tactile stimulators (Tactor, Dancer Design, Merseyside) were attached to the thumb tip by using double-sided stickers and further wrapped with a conductor such that participants could freely interact with the capacitive smartphone touchscreen. The solenoid was activated by using square wave pulses (10 ms) spaced by a uniform distribution of intervals spanning 0.75 s to 1 s. A copy of the stimulation trigger (TTL) was registered by the EEG equipment. The thumb was further covered with a conductive surface (common aluminum foil) ensuring that all the touches were translated to touchscreen events and that the same part of the thumb was used to target the screen.

### Movement sensor recordings

The right thumb flexions were tracked using a movement sensor (Flex Sensor, 112 mm, Digi-Key, Thief River Falls). The sensor was attached to the thumb (dorsum) using a custom-built jacket that allowed the sensor to bend within the jacket without the sensor being pulled. The analog signals from the sensor were digitized at 1 kHz using Labview via the USB 6008 DAQ (National Instruments, Austin, USA). The same DAQ was also used to power the sensor. The thumb was able to freely move on the touchscreen under this configuration. As the EEG and the movement sensors operated on different clocks, they were synchronized using common TTL pulse bursts generated by using an IBM T 42 computer running MATLAB. The movement signals were bandpass filtered in the range of 1 to 10 Hz (for an example of the recorded kinematic signals see Supplementary Methods).

### Alignment of smartphone data to the common laboratory clock

We formulated a data-driven method to align the smartphone data – recorded using the smartphone operating system (Android) clock – to the common laboratory clock (used for EEG, force sensor and movement-sensor recordings). As TTL pulses could not be injected into the smartphone (without software adjustments), we trained a model to link the movement sensor signal to the force sensor signals. The model was a global bidirectional LSTM (BI-LSTM) regression model that used movement sensor values and 100 equally weighted extracted moving averages values (calculated over a sliding window of 10 ms) as inputs (28). The model predicted the force of touch, we inferred that the high model predicted force emulated a touchscreen interaction. Mean squared error was chosen as the model cost function. The architecture consisted of 2 BI-LSTM layers followed by a fully connected layer. Before training the model, each participant’s data was split into the train (80 %), validation (10 %), and test (10 %) sets. The model was trained in batches of 10 (z-score normalized) samples obtained by randomly selecting sequences of length 1000 ms. Following the training, we obtained a mean squared error (capturing the difference between real force vs. predicted force) of 0.1120 on the test set, 0.1097 on the train set, and 0.0877 on the validation set. The alignment was performed by correcting for the delay between the touchscreen interactions and the model-predicted force. As the model relied on movement sensor signal fluctuations, subjects without systematic movement sensor signals surrounding the smartphone touches, and subjects where the signals appeared misaligned were eliminated resulting in 68 participants for further consideration (see Supplementary Methods).

### Identification of goal and non-goal-directed movements

After the technical alignment, we identified goal and non-goal-directed movements based on the outputs of a (separate from the one used for alignment) artificial neural network (ANN) trained to identify smartphone interactions using kinematic inputs. This involved two methodological steps. First, at the level of each individual a classification ANN was trained with touchscreen interactions and z-score normalized approximate integrals extracted from processed movement sensor signals. To decrease class imbalance, touchscreen interactions were padded with ± 30 samples and the datasets were undersampled by a factor of 10 before training. Second, the model predictions were contrasted against the real outputs and non-goal-directed movements were identified based on a predicted interaction that did not coincide with real touchscreen interaction (false-positive errors, Supplementary Figure 1a-a’).

The ANN model contained a 1-D Convolutional layer with 100 kernels for automatic feature extraction. Followed by three BI-LSTM layers. Three Dropout layers, with a dropout rate of 0.5, were applied in between each BI-LSTM layer to prevent overfitting. Finally, a fully connected layer with sigmoid activation was used for the classification. Binary cross entropy was chosen as the model cost function. The ANNs were trained in batches of 10 samples obtained by randomly selecting sequences of length 200 ms. After undersampling of the data, a sequence of length 200 represented 2 s of the raw data. Considering that the movement generally occurred within a duration of 2 s (Supplementary Figure 2), the chosen sequence length was enough to capture a movement. For one pass of the training dataset (epoch), the number of generated batches was calculated as the total length of the train data divided by 200 ms. A maximum of 100 epochs was set for training. However, the ANNs were stopped after 30 epochs with no improvement in the number of true positive predictions in the validation set. Furthermore, the learning rate was adjusted by a factor of 0.1 after no improvement for 5 epochs on the number of true positives in the validation set. For the validation set, no shuffling or random sampling was used, essentially keeping the validation data across epochs the same allowing for a direct comparison between the number of true positives in the validation set. For the final evaluation, we used the F2 score due to its higher emphasis on recall as opposed to precision (see Supplementary Methods for model hyperparameters).

After training, goal and non-goal-directed movements were identified based on the model predictions. The ANN made continuous predictions of the probability of a class label, a value between 0 and 1. The final predictions of each model were selected by comparing each output probability to a threshold (between 0 and 1) and by assigning class 1 when the value was above the selected threshold and class 0 otherwise. The F2 score was calculated for every possible threshold and the threshold that yielded the highest F2 score was selected. For each participant, the model predictions above the threshold with the highest F2 score were used to identify the goal and non-goal-directed movements. The ANN was trained with a window of touchscreen interactions. Consequently, the predicted peaks were expected approximately around (and not exactly at) the interaction. Any predicted peak around +/- 100 ms of the touchscreen interaction was considered a correct prediction (goal-directed movement). The value of +/- 100 ms was selected based on the peak width of the model predictions averaged across all participants (Supplementary Figure 4). Finally, a non-goal-directed movement was identified with a model prediction where there was no touchscreen interaction in the vicinity (false-positive errors). This was defined as any peak of a model prediction further than +/- 1 s from the interaction, based on the typical movement completion durations.

### EEG data collection and pre-processing

EEG data were collected while subjects were comfortably seated in a faraday cage. Sixty-four channel EEG caps with equidistant electrodes were used (Easycap GmbH, Wörthsee, Germany) in conjunction with ABRALYT HiCl electrode gel. The data was gathered using the 64-channel DC amplifier BrainAmp (Brain Products GmbH, Gilching). The signals were recorded and digitized at 1 kHz. All of the EEG data processing was performed offline using EEGLAB running on MATLAB (MathWorks, Natick)(29). All channels with impedances higher than 10 kΩ were removed and then subsequently replaced by interpolation. Furthermore, we used Independent Component Analysis to remove the blink-related artifacts (30). Towards the analysis of event-related potentials, the data were bandpass filtered between 0.5 Hz and 30 Hz, and for event-related spectral analysis, the data were bandpass filtered between 0.5 Hz and 45 Hz. The two ocular electrodes placed under the eyes were removed from the statistical analyses.

Towards the event-related analysis surrounding the movements and touchscreen interactions, the data was epoched using a –2 s to + 2 s window surrounding the event, and the period between −2 s to −1.5 s was used as a baseline. Trials crossing ± 80 μV were rejected as measurement artifacts. Further statistical analysis was performed on participants with greater than 50 remaining trials per movement type. Towards the analysis surrounding the artificial touches, the data was epoched using a –100 ms to 100 ms window surrounding the event, and the period between – 100 ms to – 25 ms was used as the baseline. The data was bandpass filtered between 1 Hz and 45 Hz. The spectrograms were estimated at each electrode using continuous wavelet transform (Frequencies 1 to 40 Hz, Morlet wavelet, 1 cycle-wavelet expanding to 70%).

### Statistical analysis

The event-related signals were analyzed using one-sample and paired *t*-tests (goal vs. non-goal directed movements) across all electrodes and time points (and frequency range of 1 to 40 Hz) using the mass univariate linear modeling toolbox LIMO EEG (31). Towards follow-up analysis, the same toolbox was used at the level of each individual – with movement categories and neural network model predicted peaks as a covariate – to obtain ANCOVA outputs. These outputs were then used toward population-level one-sample *t*-tests. In this follow-up applied to the frequency analysis, for computational efficiency, only the beta-band was considered (12 – 30 Hz). The statistics were based on trimmed means (20 percent). The time range considered for statistical analysis was identical to the epoching windows. The statistics were corrected for multiple comparisons by using spatiotemporal clustering as implemented in LIMO EEG (α = 0.05, 1000 bootstraps).

## Author contributions

A.G. conceived the study. A.G., R.K. and E.C. designed the study. R.K. analyzed the data aided by E.C. and A.G.. A.G. drafted the report aided by L.G. and R.K. All authors helped edit the manuscript.

## Acknowledgment

The authors would like to thank the student assistants who contributed to the data collection at Leiden University. The authors appreciate the discussions with Dr. Arnoud Visser at the University of Amsterdam and the feedback from Sebo Uithol and Gerd Tinkenhauser. We thank Evert Dekker for his assistance in developing the movement sensors used in this study.

## Data availability

All processed EEG data used in this study are shared on dataverse.nl within a month of publication, according to the Leiden University Institute of Psychology guidelines on data sharing. (Temporary link: https://surfdrive.surf.nl/files/index.php/s/ytAqMbZHTD6QqiH)

## Code availability

Codes used for processing the kinematic and EEG signals including the model settings are shared via GitHub (https://github.com/CODELABLEIDEN/Non_goal_directed_smartphone_2022).

## Related Supplementary Figures

**Supplementary Figure 1.** Identification of goal and non-goal-directed movements. **(a)** An artificial neural network was trained to identify touchscreen interactions based on movement sensor signals. The movement signal approximate integrals over 1 ms windows were used as input. **(a’)** The model well-identified movements surrounding the touchscreen interactions (goal-directed movements), but also identified highly similar movements which did not result in a touchscreen interaction (non-goal-directed movements). **(b)** The probability density of the number of events per ms for both goal and non-goal-directed movements (log10 normalized) shows that the two movement types occurred at a similar rate. **(c)** The probability density of inter-event intervals for goal and non-goal-directed movements shows a relatively broad distribution for the latter (log10 normalized). **(d)** Probability density of distance to non-goal-directed movements (log10 normalized). Non-goal-directed movements were more likely to occur after a goal-directed movement.

**Supplementary Figure 2.** Movement signal traces for goal (red, left) and non-goal-directed (black, right) movements time-locked to the predicted events for all participants (Z-score normalized for visualization). Overlaid are the median movement signals. The similarity between the movement types is indicated by the Pearson R.

**Supplementary Figure 4.** Grand averages of artificial neural network predictions (larger than optimal F2 score threshold) and probability distribution of the model predicted peaks in the sampled population.

**Supplementary figure 3.** Paired samples *t-*tests for goal and non-goal-directed movements for event-related (spectral) analysis. **(a)** Significant differences between movements for event-related analysis occurred mostly after time-locked events, shown with scalp topographies of T-values after multiple comparison corrections (MCC, p < 0.05). **(b)** Statistically, significant clusters show different event-related spectral desynchronization between the movements in the beta-band (13 –30 Hz). Approximate times were used for scalp topographies as time information was adjusted due to continuous wavelet transform.

**Supplementary figure 5.** Scalp topologies of event-related changes in the alpha-band for **(a)** goal-directed and **(b)** non-goal-directed movements. Significant statistical clusters were identified using one-sample *t*-tests and multiple comparisons corrected (MCC, p < 0.05).

## Supplementary Movies

Temporary link: https://surfdrive.surf.nl/files/index.php/s/yzBs7IWz6c4ECUP

**[Supplementary Movie 1](https://leidenuniv1-my.sharepoint.com/:v:/g/personal/kockrmd_vuw_leidenuniv_nl/EZ_8VBraS5dGtmijuoAvcGgBhPhyLcxRrOCEsAeHMtHlyA?e=TGjzap).** Event-related potentials surrounding smartphone touchscreen interactions. Statistics corresponding to one-sample *t*-test corrected for multiple comparisons (p < 0.05).

**[Supplementary Movie 2](https://leidenuniv1-my.sharepoint.com/:v:/g/personal/kockrmd_vuw_leidenuniv_nl/EVZQYcQWuA9MiqU9kRRVzesBESfJb_tBtaFPd7tEVKlFVw?e=Oy9vlU).** Event-related potentials surrounding touches on a smartphone-like surface.

**[Supplementary Movie 3](https://leidenuniv1-my.sharepoint.com/:v:/g/personal/kockrmd_vuw_leidenuniv_nl/EfQwhSEXeuRJlYP_CVpD0oMBD07-d9kvrnC_1krtZu_STQ?e=B3fwmw).** Event-related spectral perturbations surrounding smartphone touchscreen interactions collapsed across the beta-band (12 to 30 Hz) by estimating the 20% trimmed means at each time point. Statistics corresponding to one-sample *t*-test corrected for multiple comparisons, masked T-values are collapsed by using the maximum absolute amplitude).

**[Supplementary Movie 4](https://leidenuniv1-my.sharepoint.com/:v:/g/personal/kockrmd_vuw_leidenuniv_nl/EQrgOi4kRlVLge3_AbdNkYUBexs7-bGGDEb1EaUQhaVjkg?e=1Pla14).** Event-related spectral perturbations surrounding smartphone touchscreen interactions collapsed across the alpha-band (8 to 11 Hz).

**[Supplementary Movie 5](https://leidenuniv1-my.sharepoint.com/:v:/g/personal/kockrmd_vuw_leidenuniv_nl/EUsEcEtHR2xLmYAWKbxPuGsBDCp47MtrW_aPXW2UCtwILQ?e=h72P0j).** Event-related potentials surrounding goal-directed movements.

**[Supplementary Movie 6](https://leidenuniv1-my.sharepoint.com/:v:/g/personal/kockrmd_vuw_leidenuniv_nl/EYnFNFmAr5NDhkmePVLQCGYBFb5qMgdByAqR1MsCxMoLSg?e=hm84YD)**. Event-related potentials surrounding non-goal-directed movements.

**[Supplementary Movie 7](https://leidenuniv1-my.sharepoint.com/:v:/g/personal/kockrmd_vuw_leidenuniv_nl/ERwCI52erWVGh7vF_k1mIwABBSzaanc-ZJeKrw58iWj1PA?e=vZKADl).** Paired-sample *t*-test comparing event-related potentials of goal and non-goal-directed movements. Statistics corrected for multiple comparisons.

**[Supplementary Movie 8](https://leidenuniv1-my.sharepoint.com/:v:/g/personal/kockrmd_vuw_leidenuniv_nl/Eb7qPiRLgP5HuoZVt5CkbncBqyfc2Of_DPaU0HunA8o1EQ?e=1J1alg).** Event-related spectral perturbations surrounding goal-directed movements collapsed across the alpha-band (8 to 11 Hz).

**[Supplementary Movie 9](https://leidenuniv1-my.sharepoint.com/:v:/g/personal/kockrmd_vuw_leidenuniv_nl/ESwJdL4qzkdAvDFlEf3718sBiTDenebnUHJnJsTqu9XIqQ?e=6XdQjz).** Event-related spectral perturbations surrounding goal-directed movements collapsed across the beta-band (12 to 30 Hz).

**[Supplementary Movie 10](https://leidenuniv1-my.sharepoint.com/:v:/g/personal/kockrmd_vuw_leidenuniv_nl/EbDMK1q5wElJp7T2nVUiBHoBC95YQOkpOkAYrwP01zWltg?e=X7aZ5p).** Event-related spectral perturbations surrounding goal-directed movements collapsed across the gamma-band (31 to 40 Hz).

**[Supplementary Movie 11](https://leidenuniv1-my.sharepoint.com/:v:/g/personal/kockrmd_vuw_leidenuniv_nl/ERcaZ5rk9lFEkco8dr69BxMB8PqO65MkRbbhzhrwwz07_A?e=jeOHOP).** Event-related spectral perturbations surrounding non-goal-directed movements collapsed across the beta-band (12 to 30 Hz).

**[Supplementary Movie 12](https://leidenuniv1-my.sharepoint.com/:v:/g/personal/kockrmd_vuw_leidenuniv_nl/ES-YsGaRDx9KmPrTWA99NMkBGa1lJ2rKXuEB_sd94Q4utQ?e=xXNnoo).** Event-related spectral perturbations surrounding non-goal-directed movements collapsed across the gamma-band (31 to 40 Hz).

**[Supplementary Movie 13](https://leidenuniv1-my.sharepoint.com/:v:/g/personal/kockrmd_vuw_leidenuniv_nl/EVvTy-z3Ln9LoFWW-XNyOooBrGUq6OhmUKD19_C3cBE6yg?e=syqRKg).** Event-related spectral perturbations surrounding non-goal-directed movements collapsed across the alpha-band (8 to 11 Hz).

**[Supplementary Movie 14](https://leidenuniv1-my.sharepoint.com/:v:/g/personal/kockrmd_vuw_leidenuniv_nl/EYimdQTwcsNDtW5PxMttnQ4BnXXmdQb4Lh5pwjRr_w3aeQ?e=SxeLJg).** Paired-sample *t*-test comparing event-related spectral perturbations for goal and non-goal-directed movements. Statistics corrected for multiple comparisons.

**[Supplementary Movie 15](https://leidenuniv1-my.sharepoint.com/:v:/g/personal/kockrmd_vuw_leidenuniv_nl/EU5cgDVaB_lPsApjAmb9JGsBQRFswFaS8zRnU514-ZjKlA?e=P50dqG).** Event-related potentials surrounding goal-directed artificial touches.

**[Supplementary Movie 16](https://leidenuniv1-my.sharepoint.com/:v:/g/personal/kockrmd_vuw_leidenuniv_nl/Ef5DJ3kcjMJLiURkBMuldDIBXYXzNtkBGKC4-b4ygJ839g?e=d13f27).** Event-related potentials surrounding non-goal-directed artificial touches.

**[Supplementary Movie 17](https://leidenuniv1-my.sharepoint.com/:v:/g/personal/kockrmd_vuw_leidenuniv_nl/EeYsHV4U4J5Om9o_wR5WOMMBsIy4GpmIaDyx_5Gq69DGfA?e=tsInXP).** Illustrative video of the experimental setup showing participant scrolling on a smartphone with the movement sensor attached to the right thumb.

## References

1. J. N. Ingram, K. P. Körding, I. S. Howard, D. M. Wolpert, The statistics of natural hand movements. Exp. Brain Res. 188, 223–236 (2008).

2. C. Kayser, K. P. Körding, P. König, Processing of complex stimuli and natural scenes in the visual cortex. Curr. Opin. Neurobiol. 14, 468–473 (2004).

3. S. Sonkusare, M. Breakspear, C. Guo, Naturalistic Stimuli in Neuroscience: Critically Acclaimed. Trends Cogn. Sci. 23, 699–714 (2019).

4. J. N. Ingram, D. M. Wolpert, “Chapter 1 - Naturalistic approaches to sensorimotor control” in Progress in Brain Research, Enhancing Performance for Action and Perception., A. M. Green, C. E. Chapman, J. F. Kalaska, F. Lepore, Eds. (Elsevier, 2011), pp. 3–29.

5. M. Balerna, A. Ghosh, The details of past actions on a smartphone touchscreen are reflected by intrinsic sensorimotor dynamics. Npj Digit. Med. 1, 4 (2018).

6. B. E. Kilavik, M. Zaepffel, A. Brovelli, W. A. MacKay, A. Riehle, The ups and downs of beta oscillations in sensorimotor cortex. Exp. Neurol. 245, 15–26 (2013).

7. R. Qing Cui, L. Deecke, High Resolution DC-EEG Analysis of the Bereitschaftspotential and Post Movement Onset Potentials Accompanying Uni-or Bilateral Voluntary Finger Movements. Brain Topogr. 11, 233–249 (1999).

8. R. Kristeva, D. Cheyne, W. Lang, G. Lindinger, L. Deecke, Movement-related potentials accompanying unilateral and bilateral finger movements with different inertial loads. Electroencephalogr. Clin. Neurophysiol. 75, 410–418 (1990).

9. T. W. Picton, The P300 Wave of the Human Event-Related Potential. J. Clin. Neurophysiol. 9, 456–479 (1992).

10. E. Houdayer, S.-J. Lee, M. Hallett, Cerebral preparation of spontaneous movements: An EEG study. Clin. Neurophysiol. 131, 2561–2565 (2020).

11. H. Shibasaki, M. Hallett, What is the Bereitschaftspotential? Clin. Neurophysiol. 117, 2341– 2356 (2006).

12. L. Leocani, C. Toro, P. Zhuang, C. Gerloff, M. Hallett, Event-related desynchronization in reaction time paradigms: a comparison with event-related potentials and corticospinal excitability. Clin. Neurophysiol. 112, 923–930 (2001).

13. A. Mehrabian, S. L. Friedman, An analysis of fidgeting and associated individual differences. J. Pers. 54, 406–429 (1986).

14. N. Sendhilnathan, D. Basu, M. E. Goldberg, J. D. Schall, A. Murthy, Neural correlates of goal-directed and non–goal-directed movements. Proc. Natl. Acad. Sci. 118 (2021).

15. J. Pereira, P. Ofner, A. Schwarz, A. I. Sburlea, G. R. Müller-Putz, EEG neural correlates of goal-directed movement intention. NeuroImage 149, 129–140 (2017).

16. E. Misirlisoy, P. Haggard, Veto and Vacillation: A Neural Precursor of the Decision to Withhold Action. J. Cogn. Neurosci. 26, 296–304 (2014).

17. H. Shibasaki, M. Kato, Movement-associated cortical potentials with unilateral and bilateral simultaneous hand movement. J. Neurol. 208, 191–199 (1975).

18. J. Duque, I. Greenhouse, L. Labruna, R. B. Ivry, Physiological Markers of Motor Inhibition during Human Behavior. Trends Neurosci. 40, 219–236 (2017).

19. A. Urbano, C. Babiloni, P. Onorati, F. Babiloni, Dynamic functional coupling of high resolution EEG potentials related to unilateral internally triggered one-digit movements. Electroencephalogr. Clin. Neurophysiol. 106, 477–487 (1998).

20. M. G. Lacourse, E. L. R. Orr, S. C. Cramer, M. J. Cohen, Brain activation during execution and motor imagery of novel and skilled sequential hand movements. NeuroImage 27, 505–519 (2005).

21. M. Kobayashi, H. Théoret, A. Pascual-Leone, Suppression of ipsilateral motor cortex facilitates motor skill learning. Eur. J. Neurosci. 29, 833–836 (2009).

22. M. Kobayashi, H. Théoret, A. Pascual-Leone,Updating P300: An integrative theory of P3a and P3b. Clin. Neurophysiol. 118, 2128–2148 (2007).

23. C. Neuper, G. Pfurtscheller, Event-related dynamics of cortical rhythms: frequency-specific features and functional correlates. Int. J. Psychophysiol. 43, 41–58 (2001).

24. E. Heinrichs-Graham, M. J. Kurz, J. E. Gehringer, T. W. Wilson, The functional role of post-movement beta oscillations in motor termination. Brain Struct. Funct. 222, 3075–3086 (2017).

25. H. T. van Schie, R. B. Mars, M. G. H. Coles, H. Bekkering, Modulation of activity in medial frontal and motor cortices during error observation. Nat. Neurosci. 7, 549–554 (2004).

26. A. Riesel, The erring brain: Error-related negativity as an endophenotype for OCD—A review and meta-analysis. Psychophysiology 56, e13348 (2019).

27. P. Cisek, J. F. Kalaska, Neural Mechanisms for Interacting with a World Full of Action Choices. Annu. Rev. Neurosci. 33, 269–298 (2010).

28. A. Graves, J. Schmidhuber, Framewise phoneme classification with bidirectional LSTM and other neural network architectures. Neural Netw. 18, 602–610 (2005).

29. A. Delorme, S. Makeig, EEGLAB: an open source toolbox for analysis of single-trial EEG dynamics including independent component analysis. J. Neurosci. Methods 134, 9–21 (2004).

30. M. B. Pontifex, V. Miskovic, S. Laszlo, Evaluating the efficacy of fully automated approaches for the selection of eyeblink ICA components. Psychophysiology 54, 780–791 (2017).

31. C. R. Pernet, N. Chauveau, C. Gaspar, G. A. Rousselet, LIMO EEG: A Toolbox for Hierarchical LInear MOdeling of ElectroEncephaloGraphic Data. Comput. Intell. Neurosci. (2011) https://doi.org/10.1155/2011/831409 (January 25, 2019).

