## Supplementary figures and images for "Neural processing of goal and non-goal-directed movements on the smartphone"

### Supplementary Figure 1

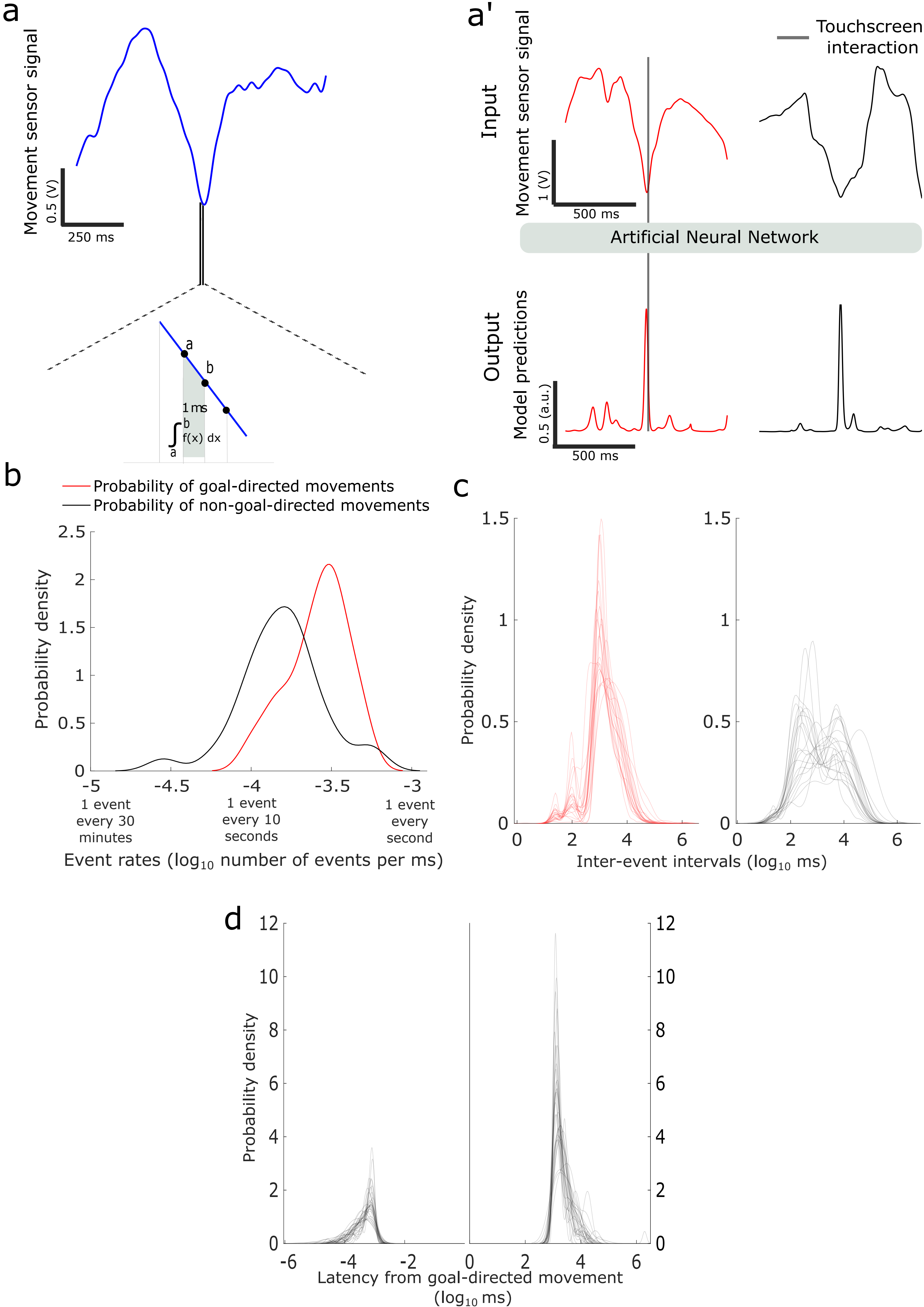

### Supplementary Figure 3

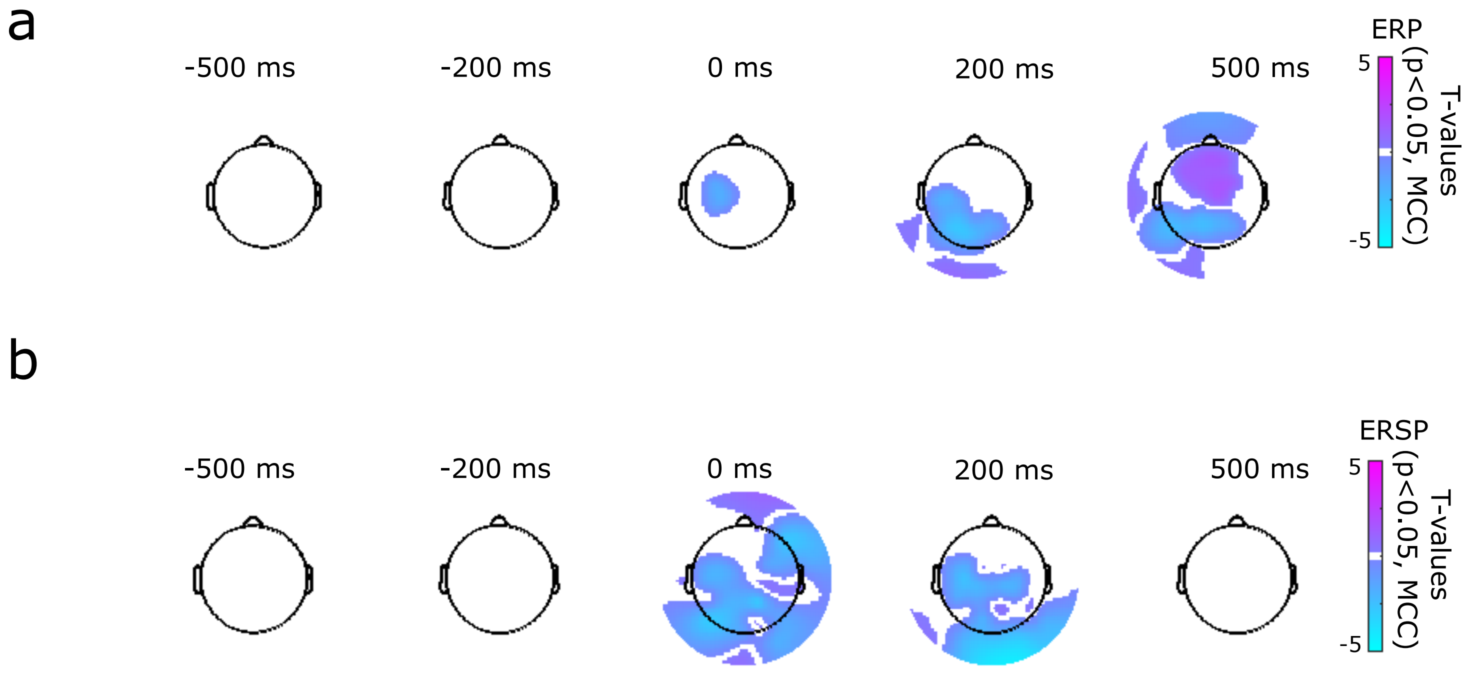

### Supplementary Figure 4

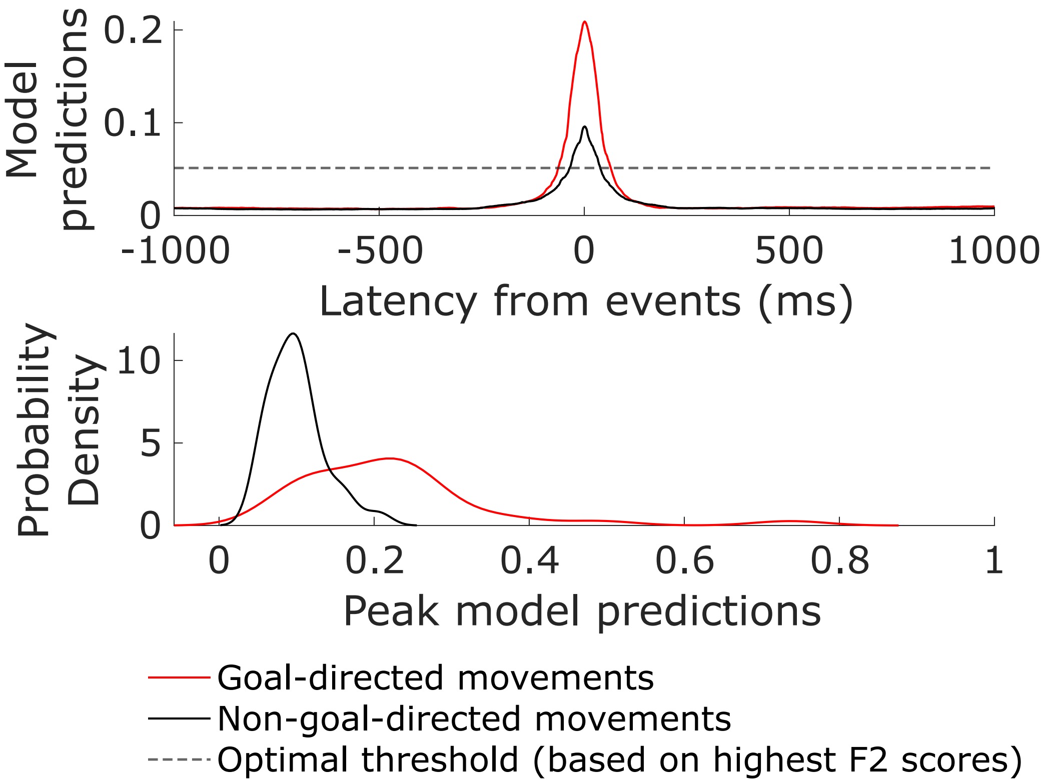

### Supplementary Figure 5

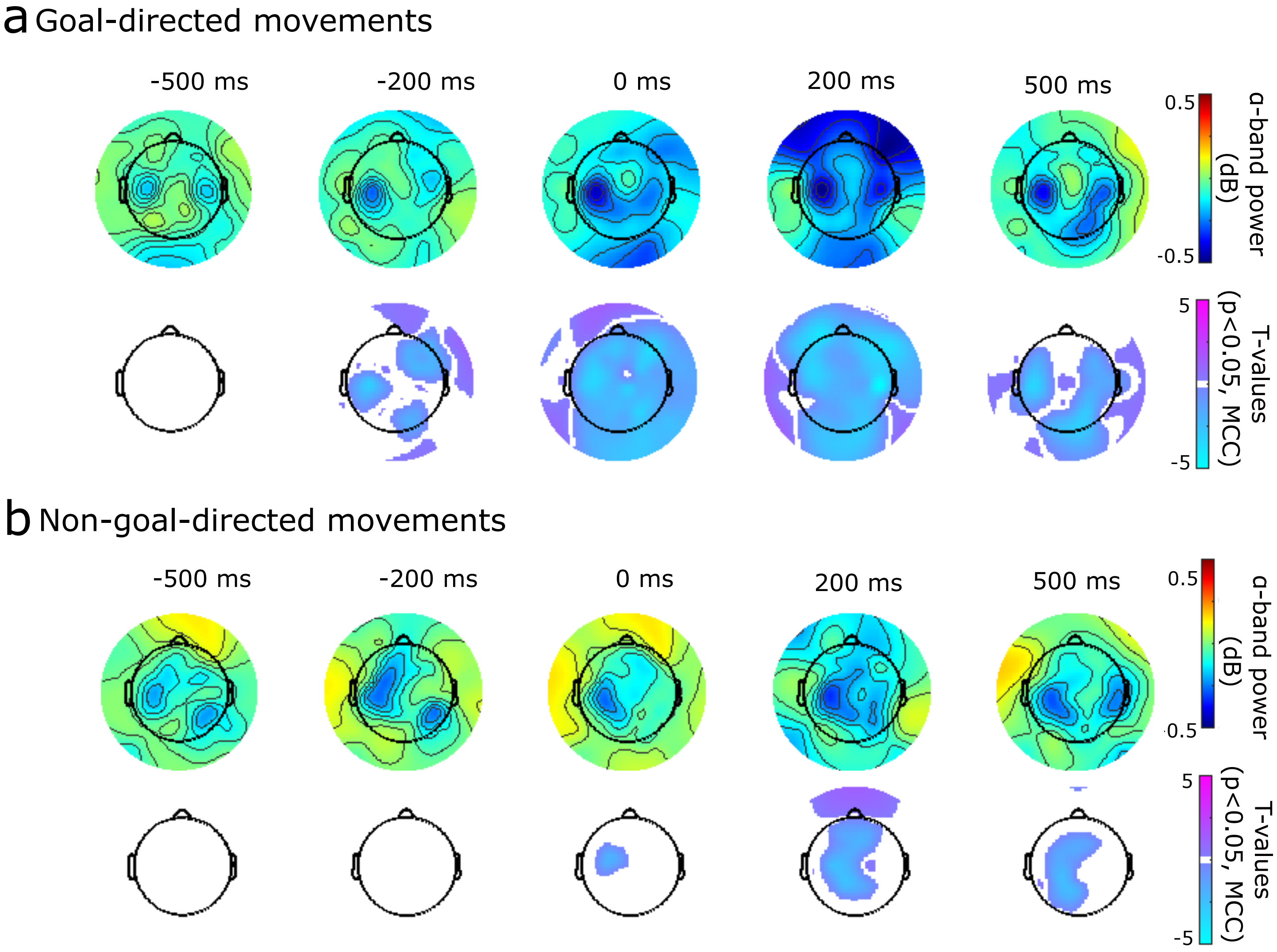
