## Supplementary Figure 2 for "Neural processing of goal and non-goal-directed movements on the smartphone"

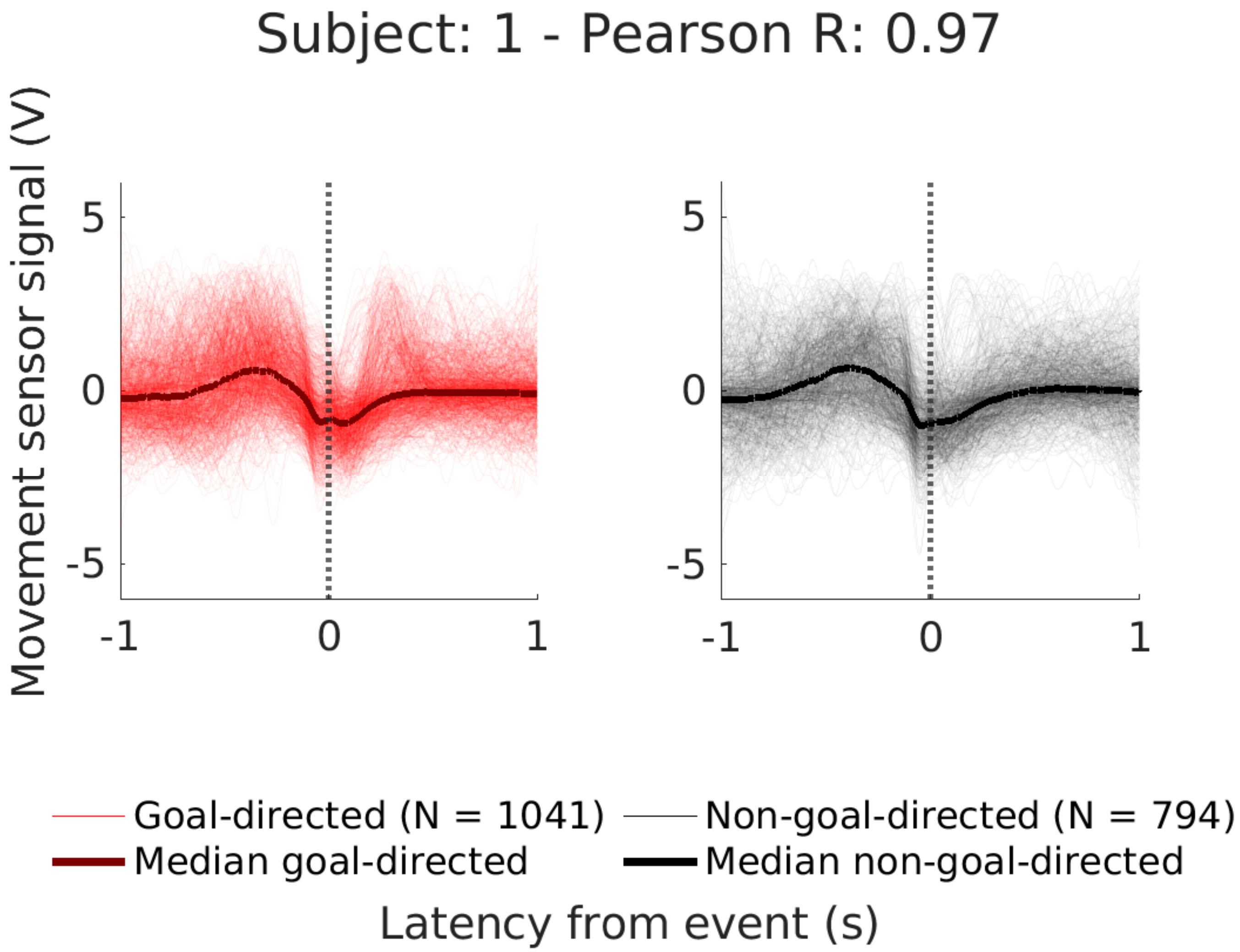
Subject: 1 - Pearson R: 0.97
Movement sensor signal (V)
5
0
-5
-1
0
1
5
0
-5
-1
0
1
Goal-directed (N = 1041)
Non-goal-directed (N = 794)
Median goal-directed
Median non-goal-directed
Latency from event (s)

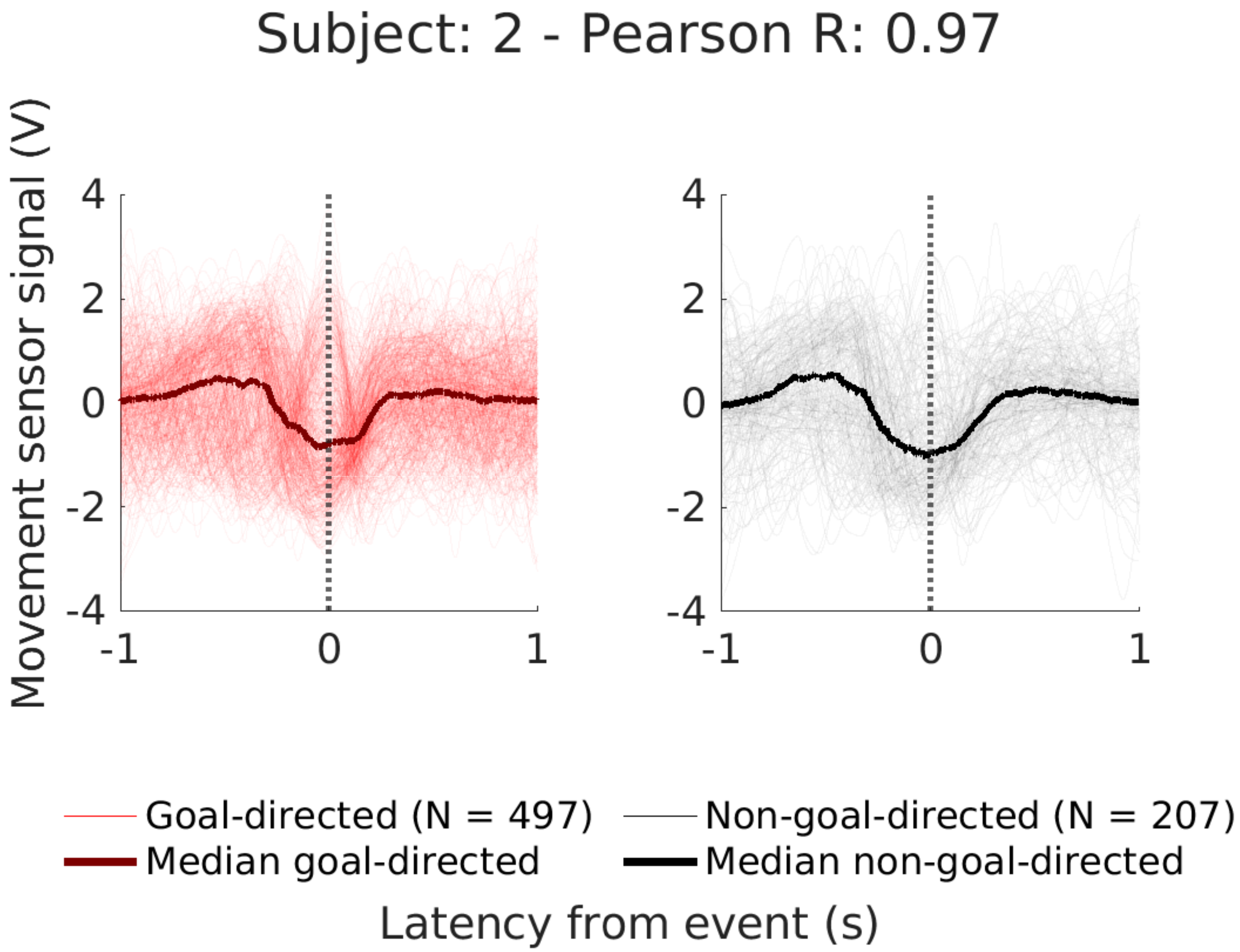

Subject: 2 - Pearson R: 0.97
Movement sensor signal (V)
4
2
0
-2
-4
-1
0
1
Goal-directed (N = 497)
Median goal-directed
Non-goal-directed (N = 207)
Median non-goal-directed
Latency from event (s)

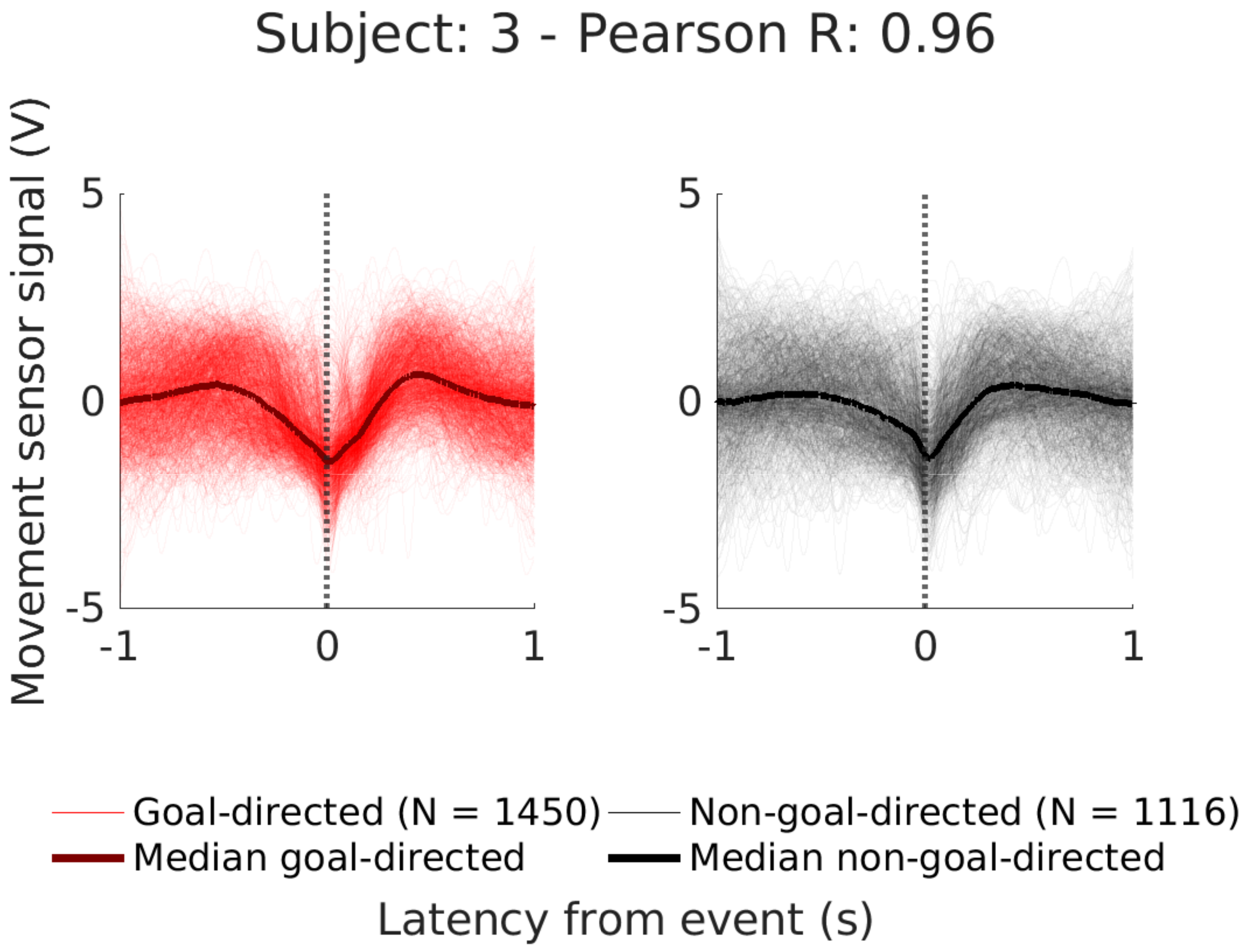
Subject: 3 - Pearson R: 0.96
Movement sensor signal (V)
5
0
-5
-1
0
1
5
0
-5
-1
0
1
Goal-directed (N = 1450)
Non-goal-directed (N = 1116)
Median goal-directed
Median non-goal-directed
Latency from event (s)

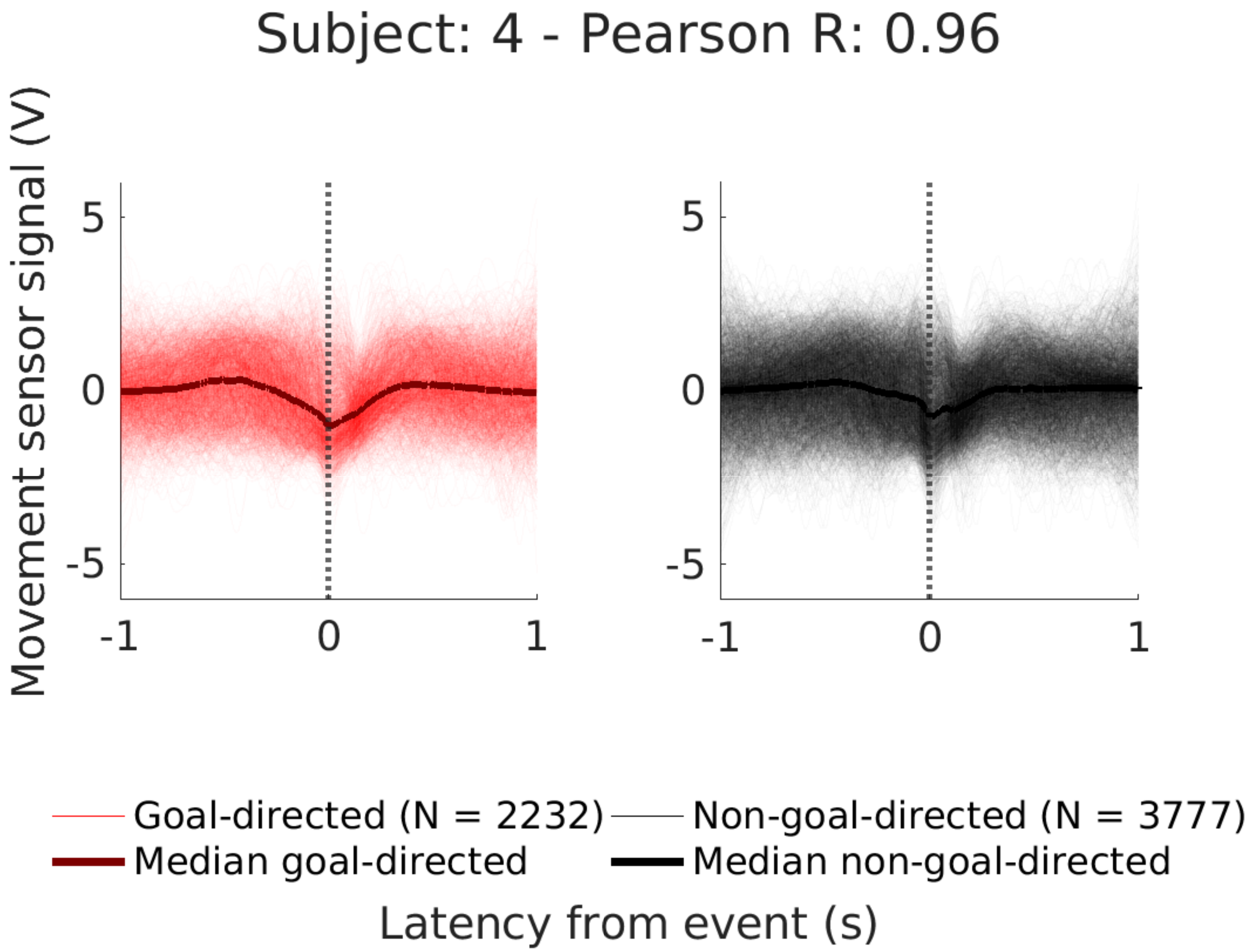
Subject: 4 - Pearson R: 0.96
Movement sensor signal (V)
5
0
-5
-1
0
1
5
0
-5
-1
0
1
Goal-directed (N = 2232)
Non-goal-directed (N = 3777)
Median goal-directed
Median non-goal-directed
Latency from event (s)

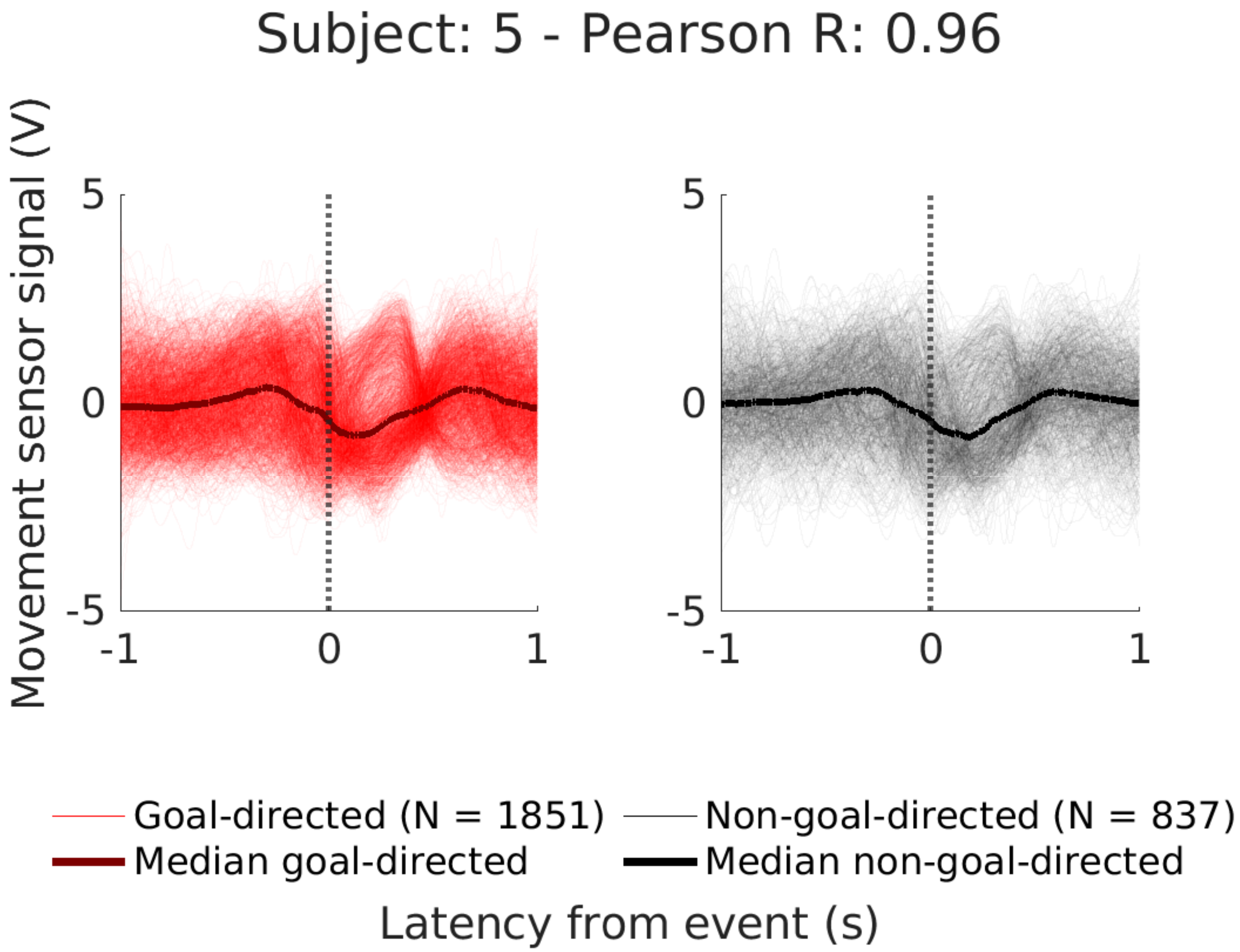

Subject: 5 - Pearson R: 0.96
Movement sensor signal (V)
5
0
-5
-1
0
1
5
0
-5
-1
0
1
Goal-directed (N = 1851)
Median goal-directed
Non-goal-directed (N = 837)
Median non-goal-directed
Latency from event (s)

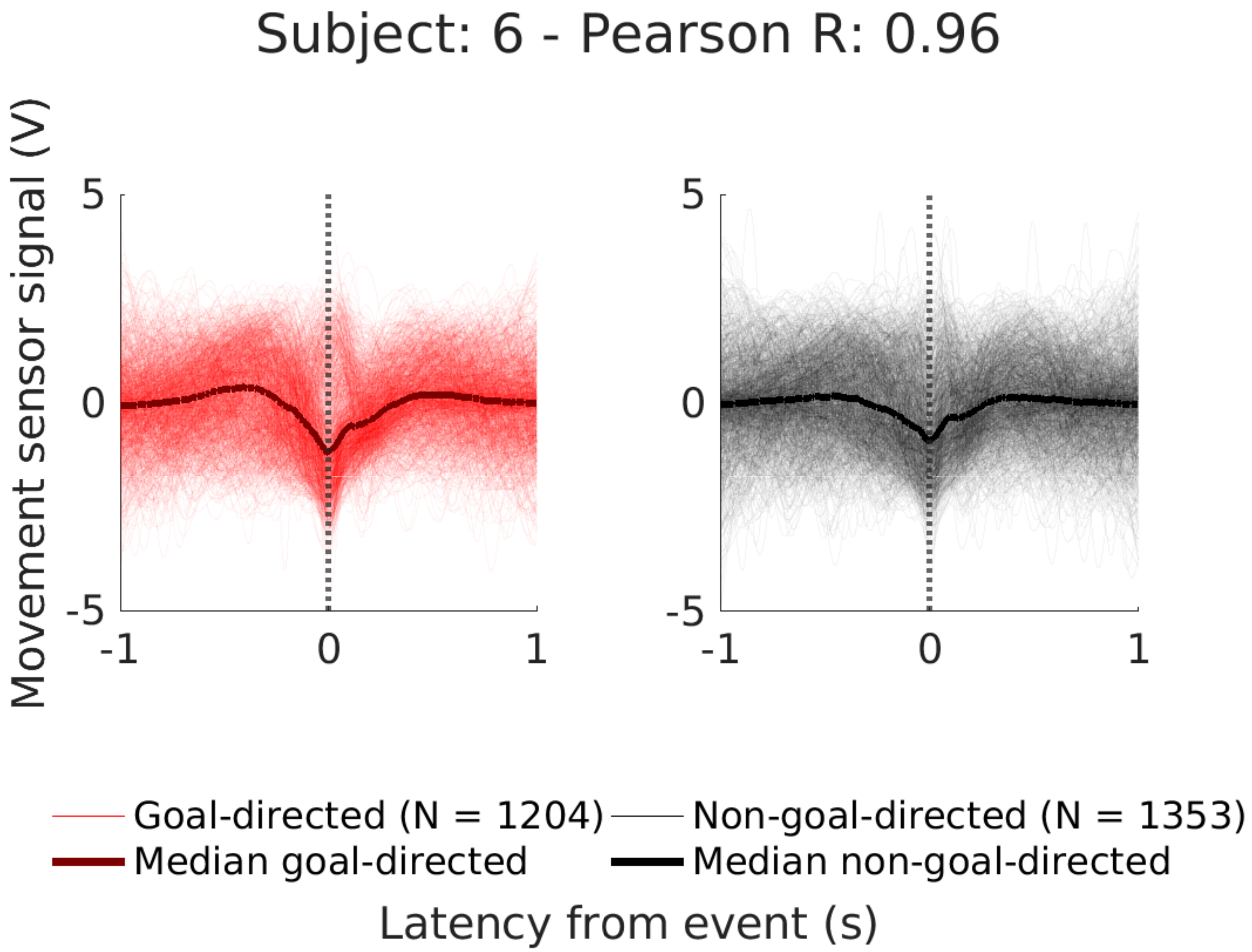
Subject: 6 - Pearson R: 0.96
Movement sensor signal (V)
5
0
-5
-1
0
1
Goal-directed (N = 1204)
Non-goal-directed (N = 1353)
Median goal-directed
Median non-goal-directed
Latency from event (s)

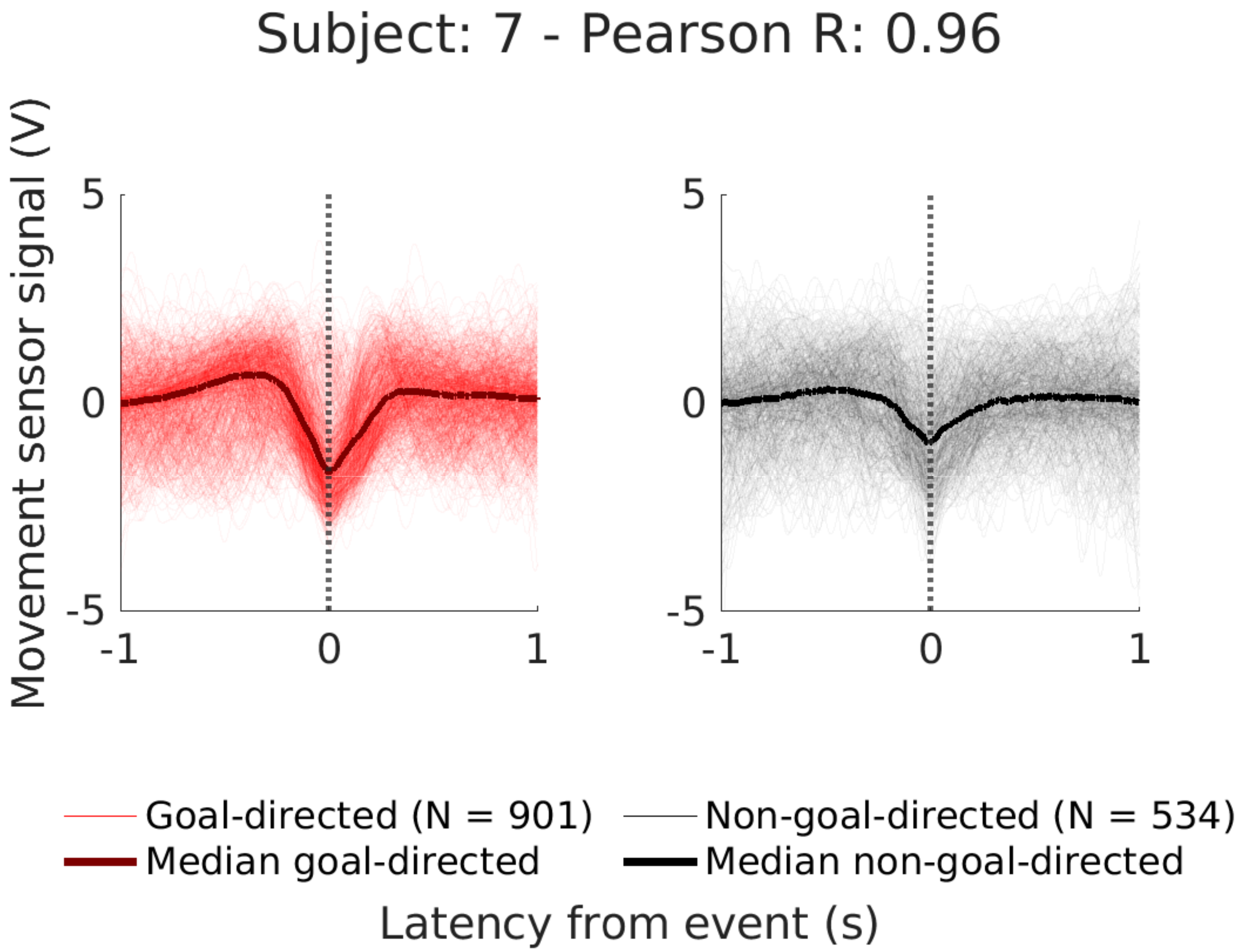

Subject: 7 - Pearson R: 0.96
Movement sensor signal (V)
5
0
-5
-1
0
1
5
0
-5
-1
0
1
Goal-directed (N = 901)
Median goal-directed
Non-goal-directed (N = 534)
Median non-goal-directed
Latency from event (s)

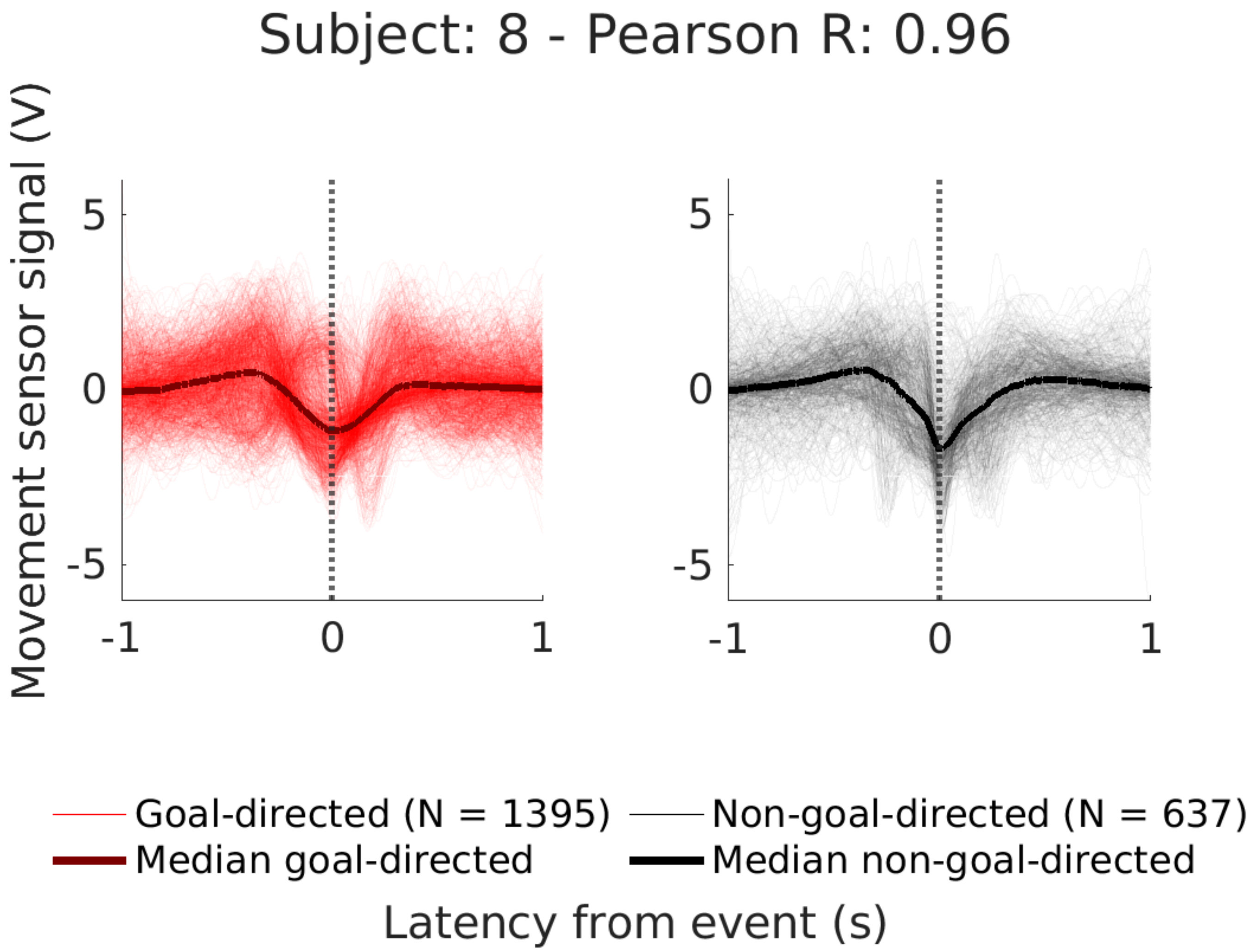

Subject: 8 - Pearson R: 0.96
Movement sensor signal (V)
5
0
-5
-1
0
1
5
0
-5
-1
0
1
Goal-directed (N = 1395)
Median goal-directed
Non-goal-directed (N = 637)
Median non-goal-directed
Latency from event (s)

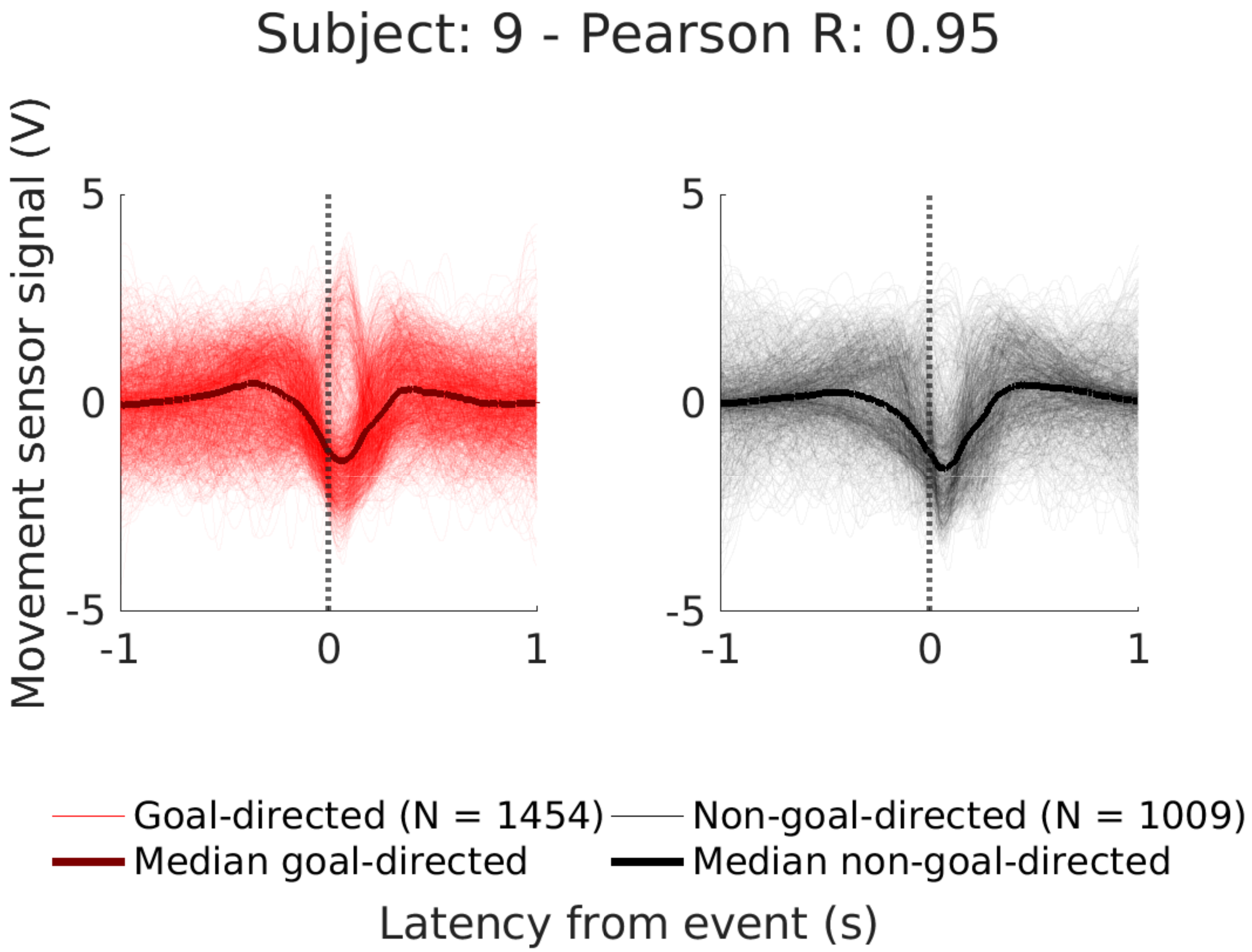
Subject: 9 - Pearson R: 0.95
Movement sensor signal (V)
5
0
-5
-1
0
1
5
0
-5
-1
0
1
Goal-directed (N = 1454)
Non-goal-directed (N = 1009)
Median goal-directed
Median non-goal-directed
Latency from event (s)

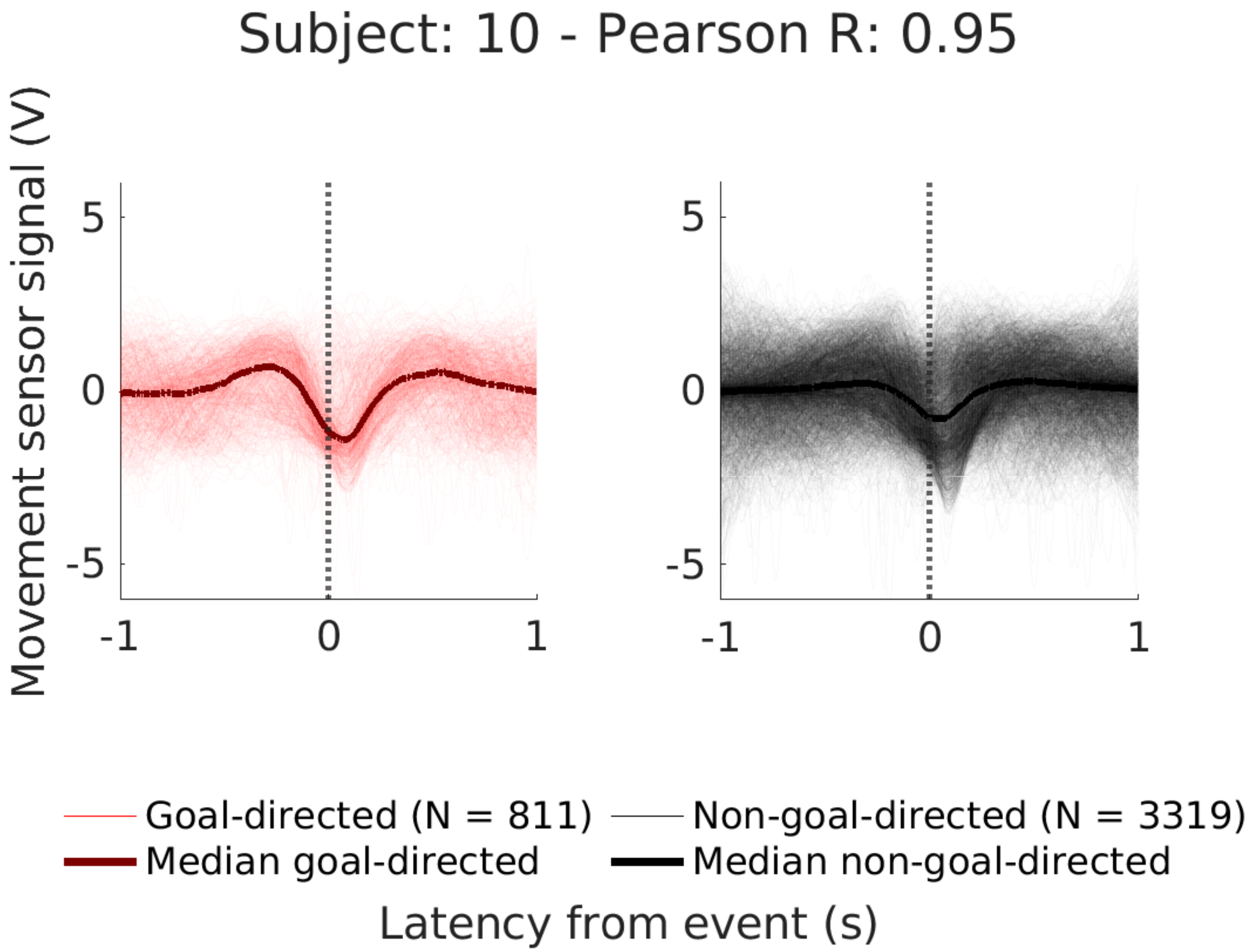

Subject: 10 - Pearson R: 0.95
Movement sensor signal (V)
5
0
-5
-1
0
1
5
0
-5
-1
0
1
Goal-directed (N = 811)
Non-goal-directed (N = 3319)
Median goal-directed
Median non-goal-directed
Latency from event (s)

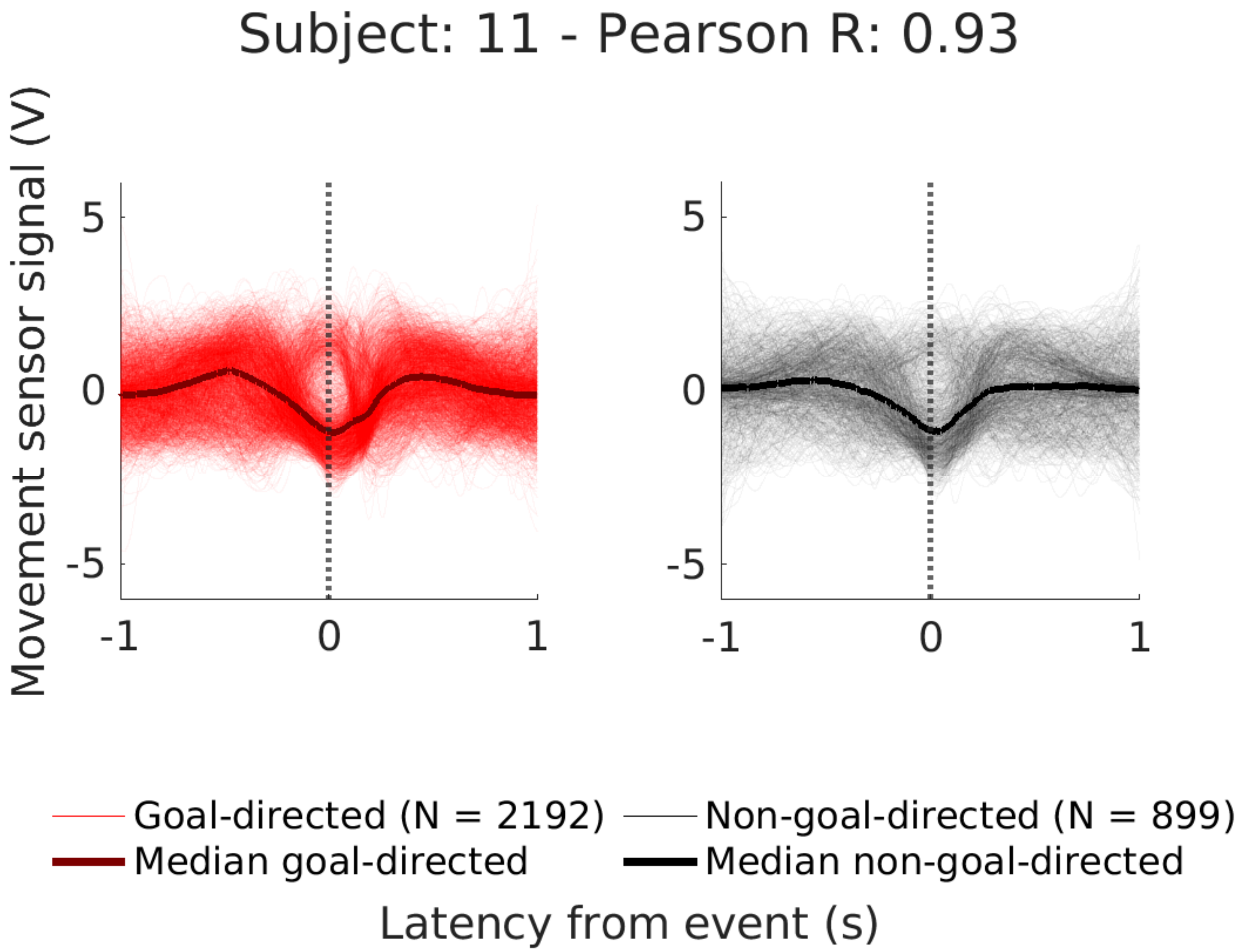
Subject: 11 - Pearson R: 0.93
Movement sensor signal (V)
5
0
-5
-1
0
1
5
0
-5
-1
0
1
Goal-directed (N = 2192)
Non-goal-directed (N = 899)
Median goal-directed
Median non-goal-directed
Latency from event (s)

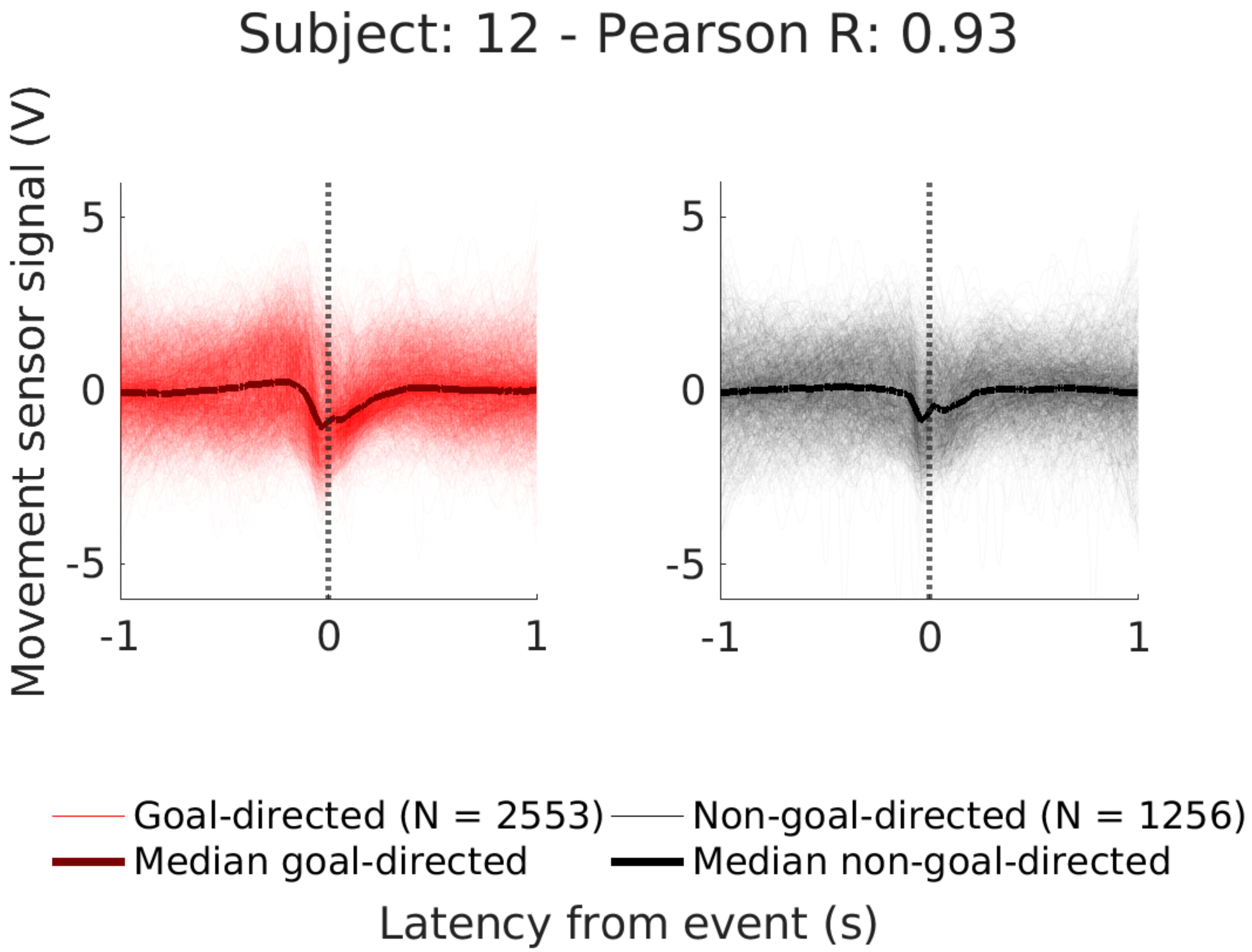

Subject: 12 - Pearson R: 0.93
Movement sensor signal (V)
5
0
-5
-1
0
1
Goal-directed (N = 2553)
Non-goal-directed (N = 1256)
Median goal-directed
Median non-goal-directed
Latency from event (s)

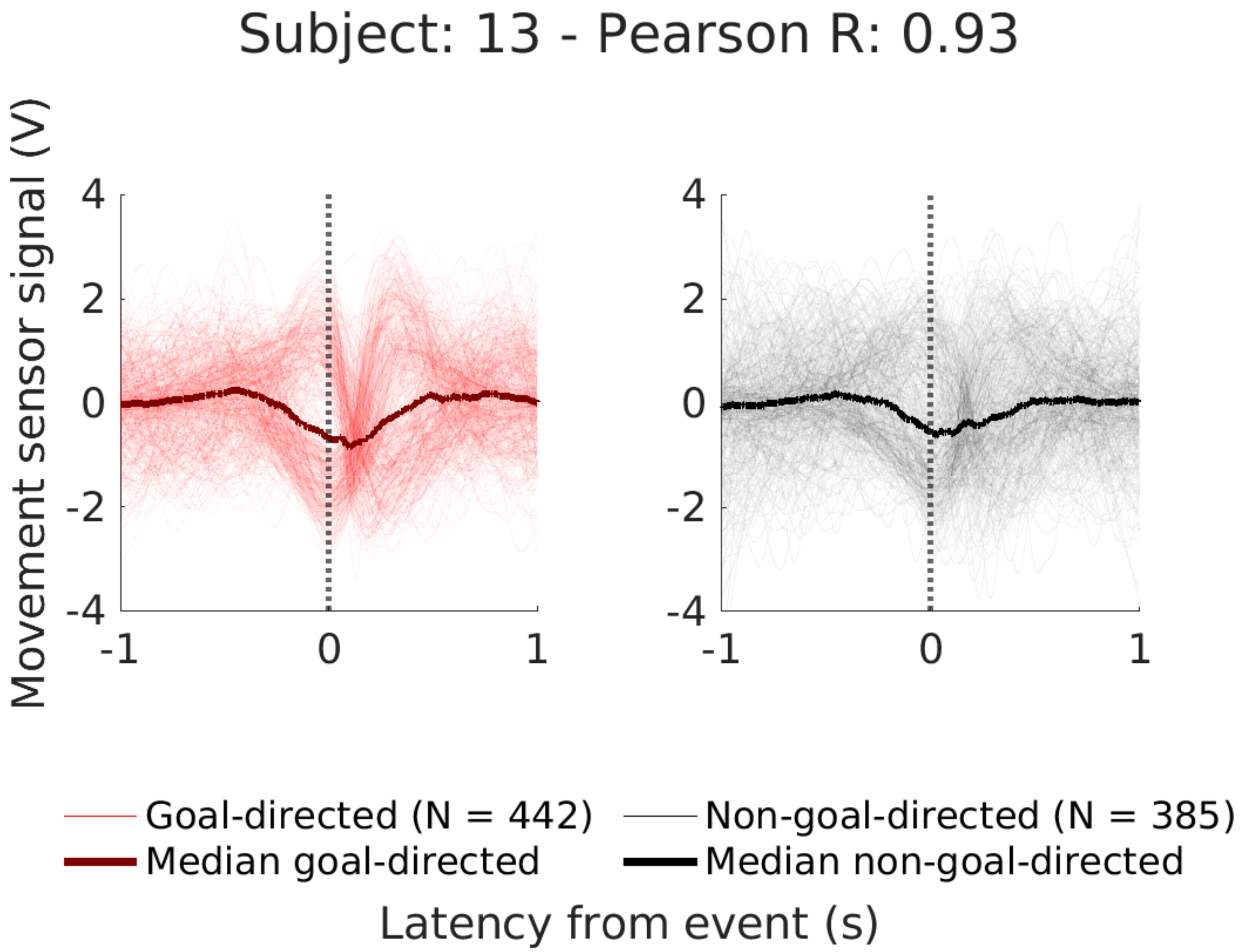

Subject: 13 - Pearson R: 0.93
Movement sensor signal (V)
4
2
0
-2
-4
-1
0
1
Goal-directed (N = 442)
Median goal-directed
Non-goal-directed (N = 385)
Median non-goal-directed
Latency from event (s)

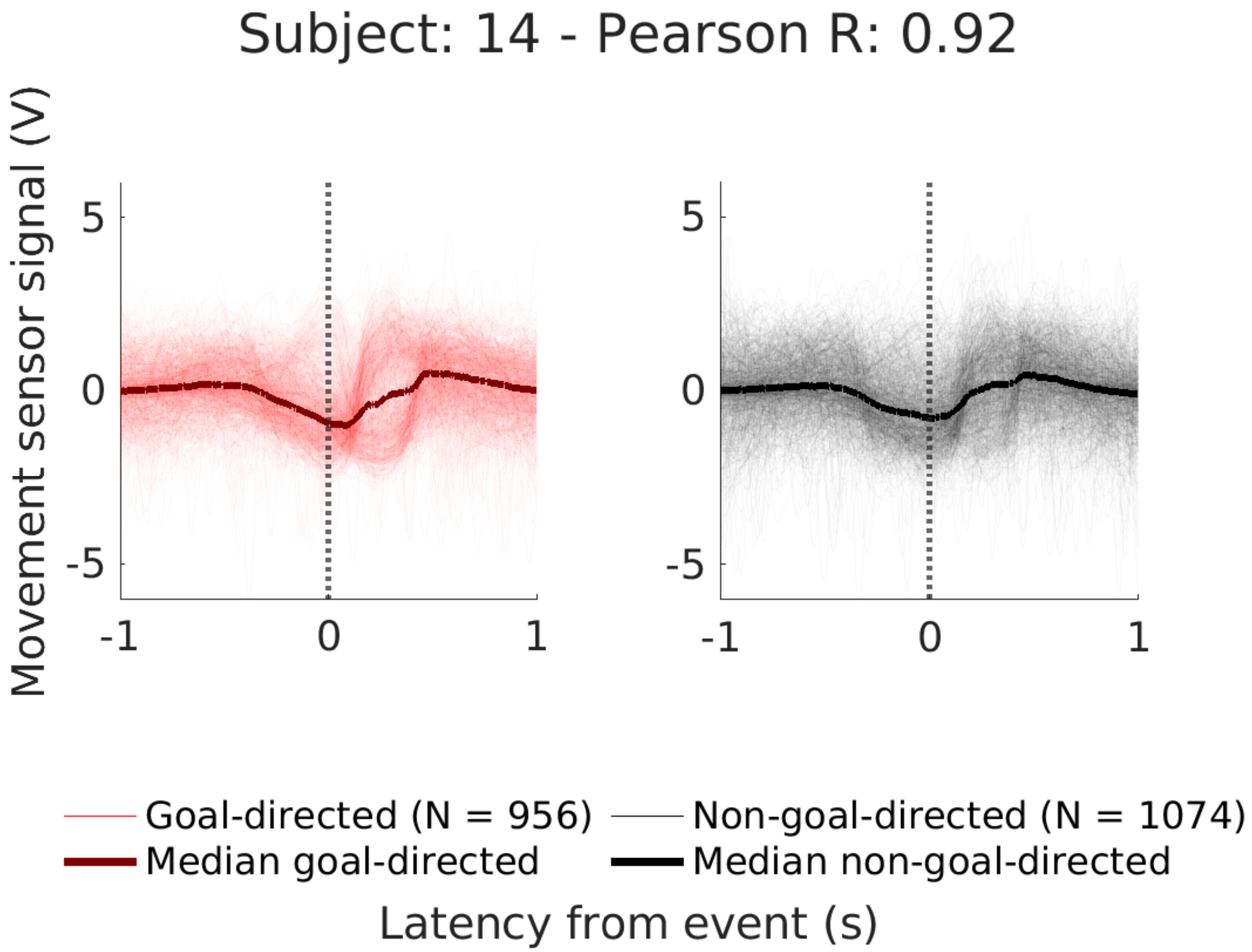

Subject: 14 - Pearson R: 0.92
Movement sensor signal (V)
5
0
-5
-1
0
1
Goal-directed (N = 956)
Non-goal-directed (N = 1074)
Median goal-directed
Median non-goal-directed
Latency from event (s)

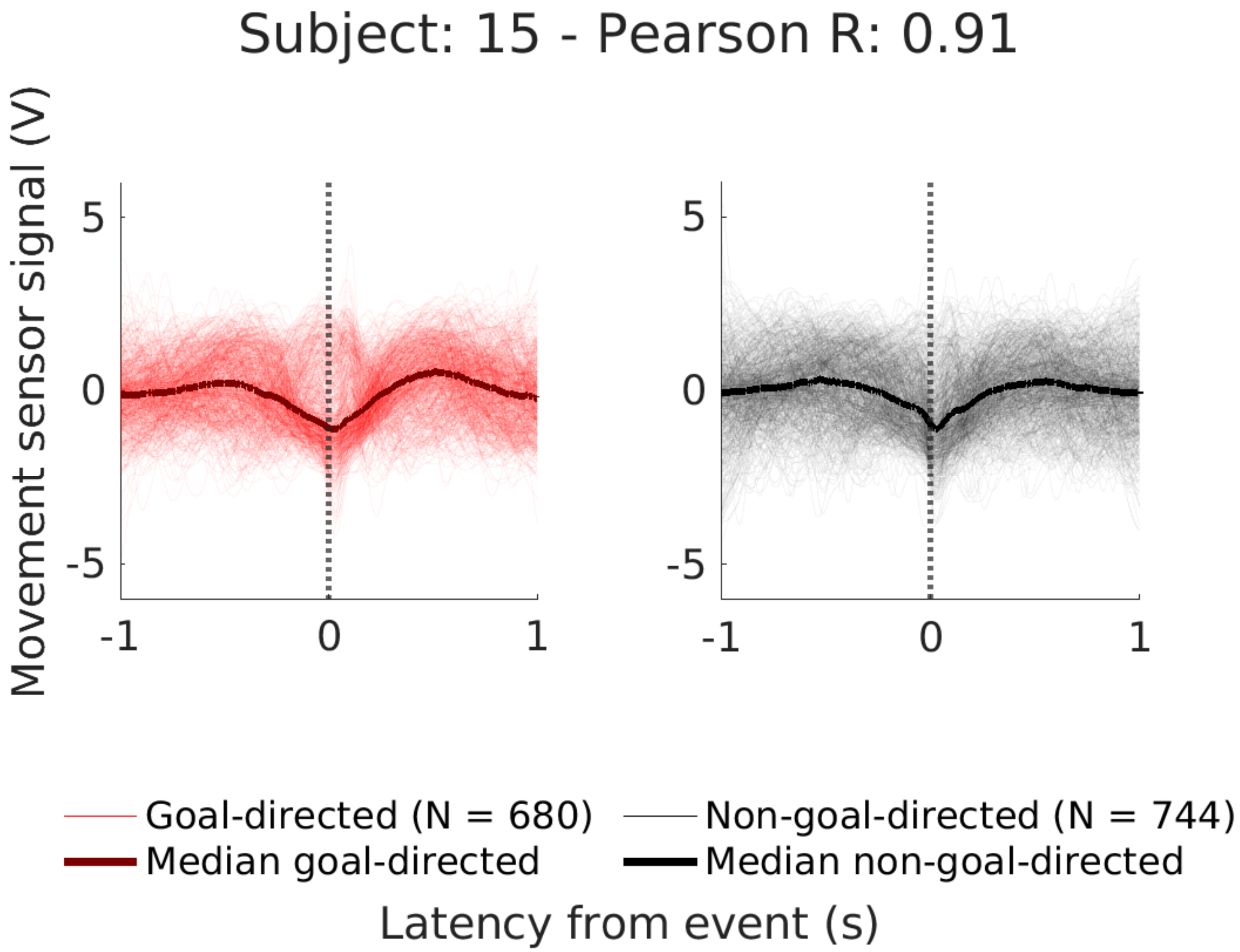
Subject: 15 - Pearson R: 0.91
Movement sensor signal (V)
5
0
-5
-1
0
1
5
0
-5
-1
0
1
Goal-directed (N = 680)
Median goal-directed
Non-goal-directed (N = 744)
Median non-goal-directed
Latency from event (s)

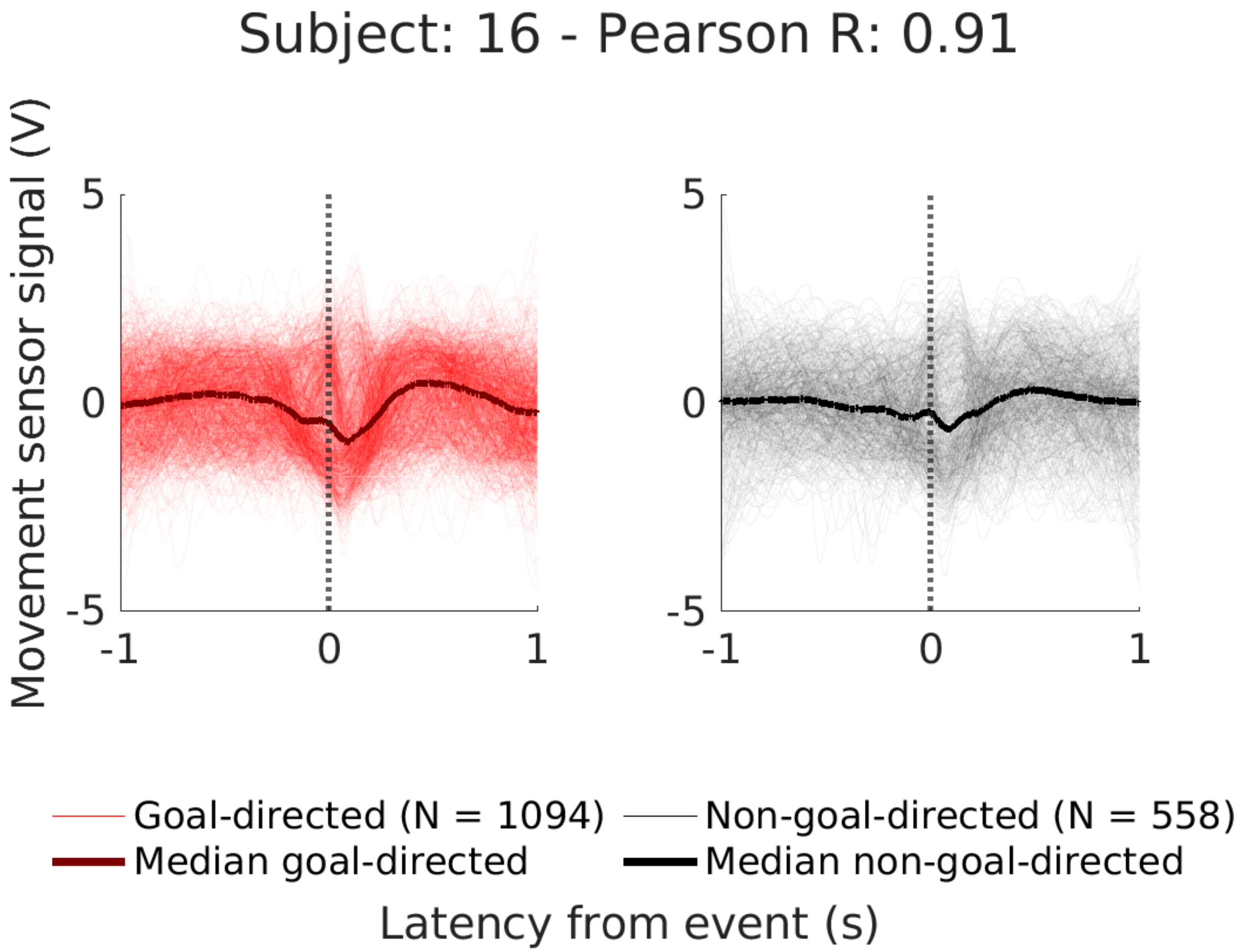

Subject: 16 - Pearson R: 0.91
Movement sensor signal (V)
5
0
-5
-1
0
1
Goal-directed (N = 1094)
Median goal-directed
Non-goal-directed (N = 558)
Median non-goal-directed
Latency from event (s)

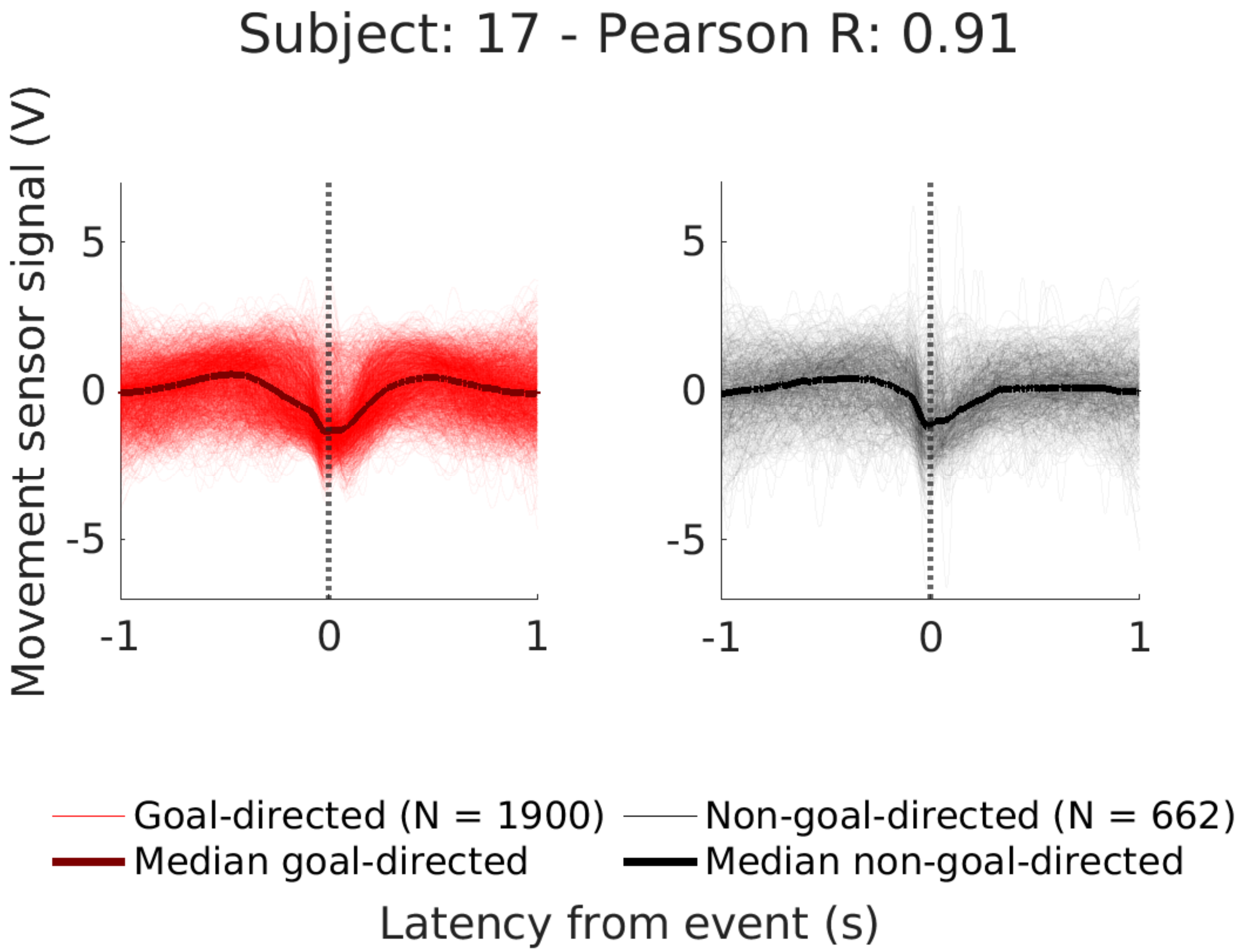
Subject: 17 - Pearson R: 0.91
Movement sensor signal (V)
5
0
-5
-1
0
1
5
0
-5
-1
0
1
Goal-directed (N = 1900)
Non-goal-directed (N = 662)
Median goal-directed
Median non-goal-directed
Latency from event (s)

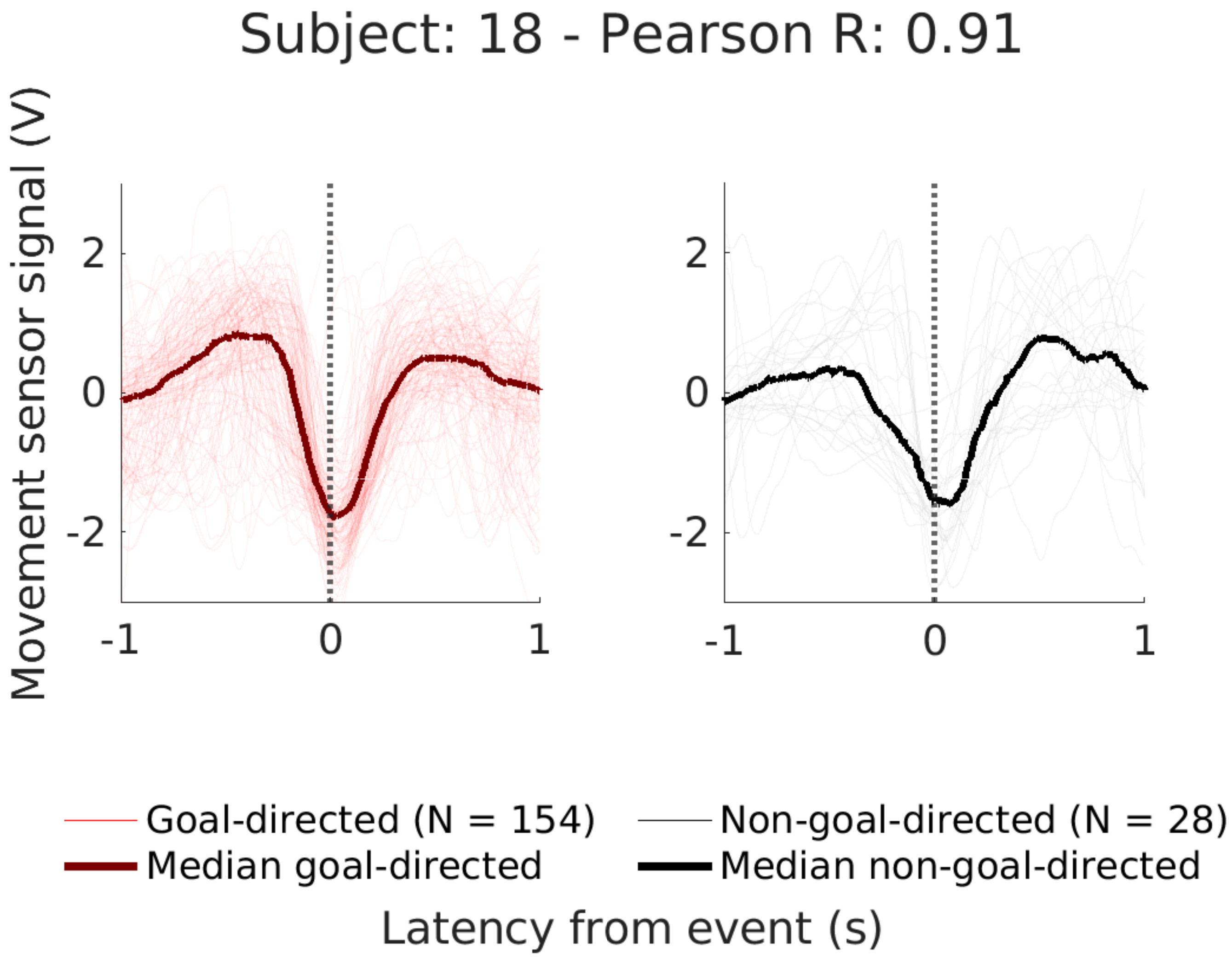
Subject: 18 - Pearson R: 0.91
Movement sensor signal (V)
2
0
-2
-1
0
1
Goal-directed (N = 154)
Median goal-directed
Non-goal-directed (N = 28)
Median non-goal-directed
Latency from event (s)

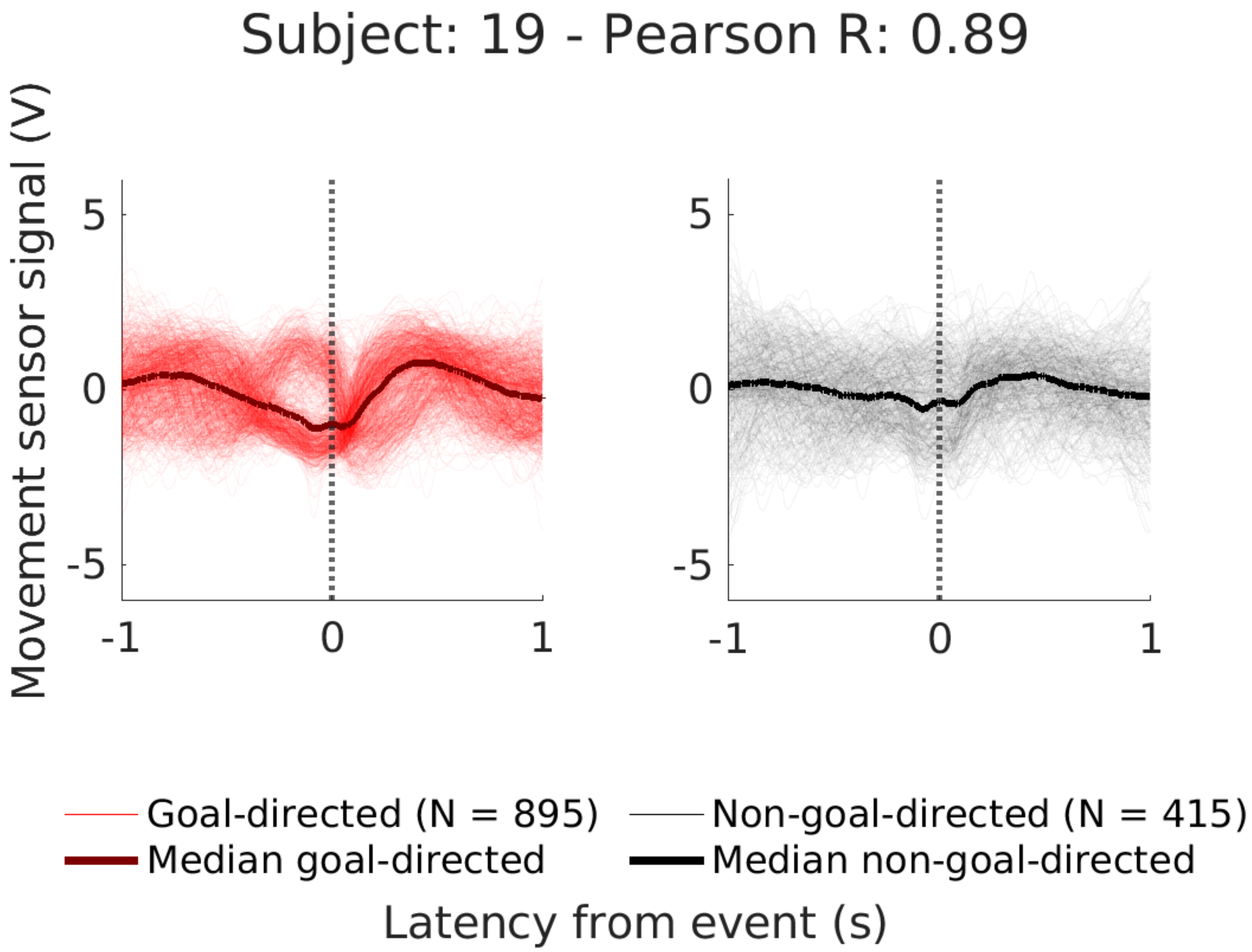
Subject: 19 - Pearson R: 0.89
Movement sensor signal (V)
5
0
-5
-1
0
1
5
0
-5
-1
0
1
Goal-directed (N = 895)
Median goal-directed
Non-goal-directed (N = 415)
Median non-goal-directed
Latency from event (s)

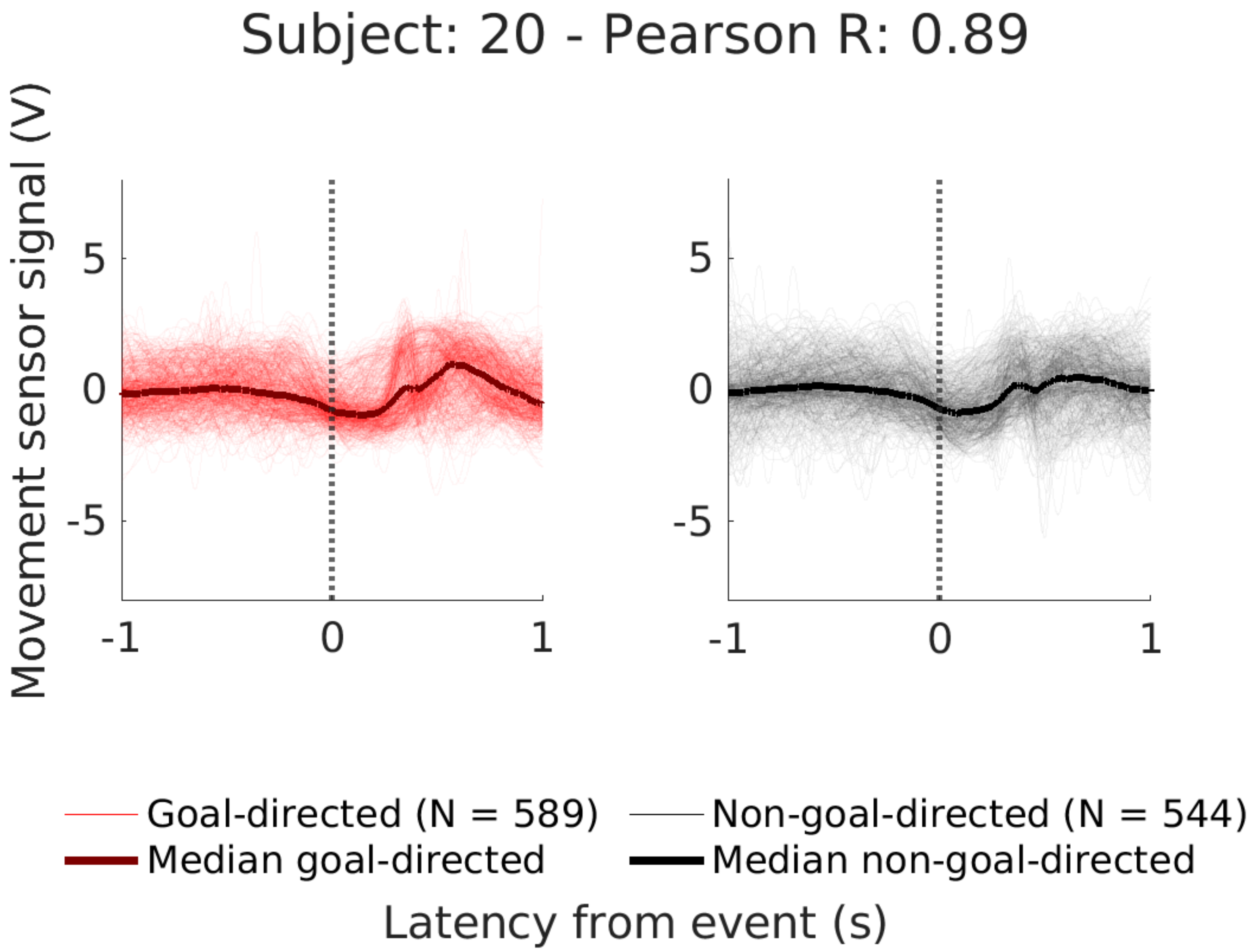

Subject: 20 - Pearson R: 0.89
Movement sensor signal (V)
5
0
-5
-1
0
1
Goal-directed (N = 589)
Non-goal-directed (N = 544)
Median goal-directed
Median non-goal-directed
Latency from event (s)

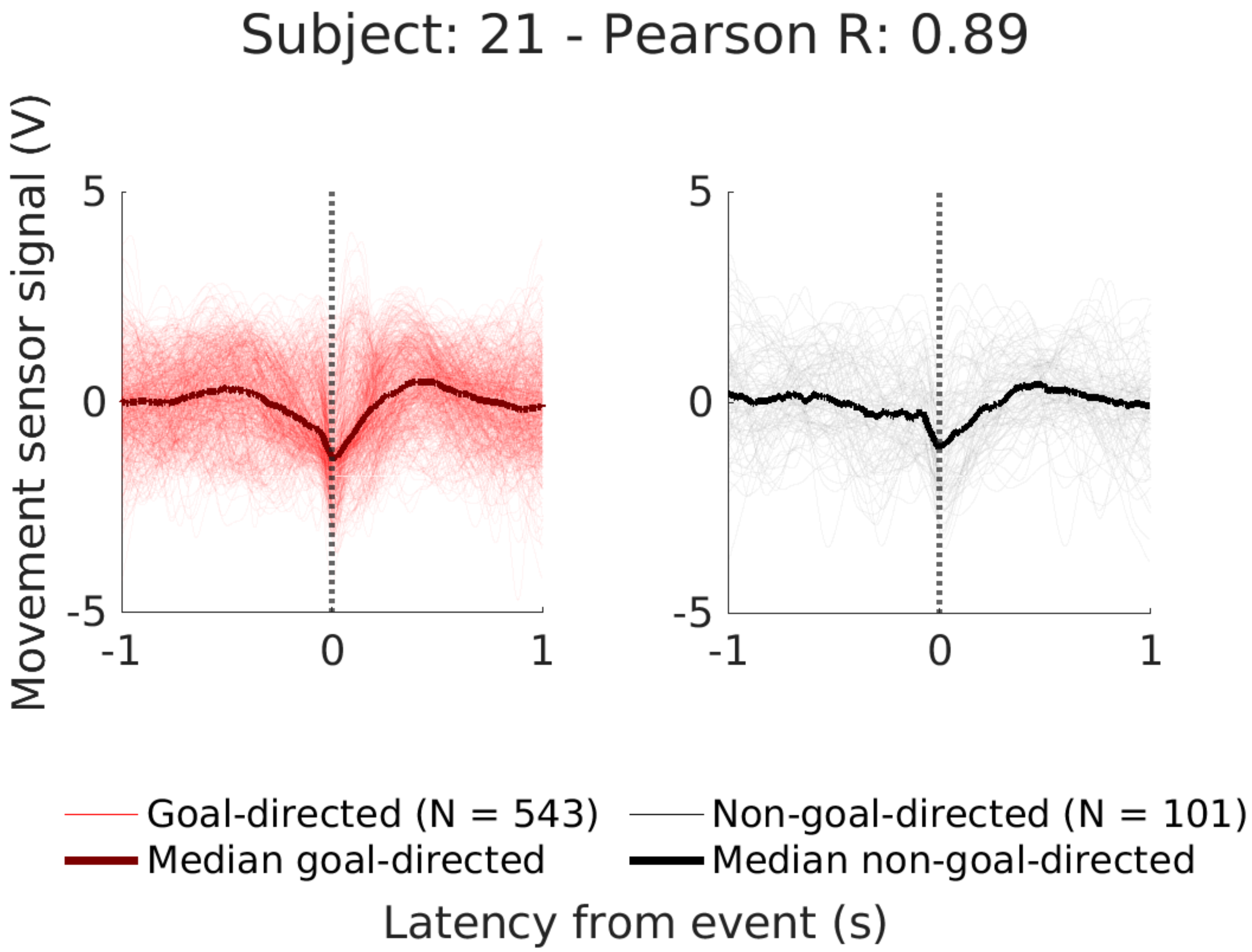

Subject: 21 - Pearson R: 0.89
Movement sensor signal (V)
5
0
-5
-1
0
1
5
0
-5
-1
0
1
Goal-directed (N = 543)
Median goal-directed
Non-goal-directed (N = 101)
Median non-goal-directed
Latency from event (s)

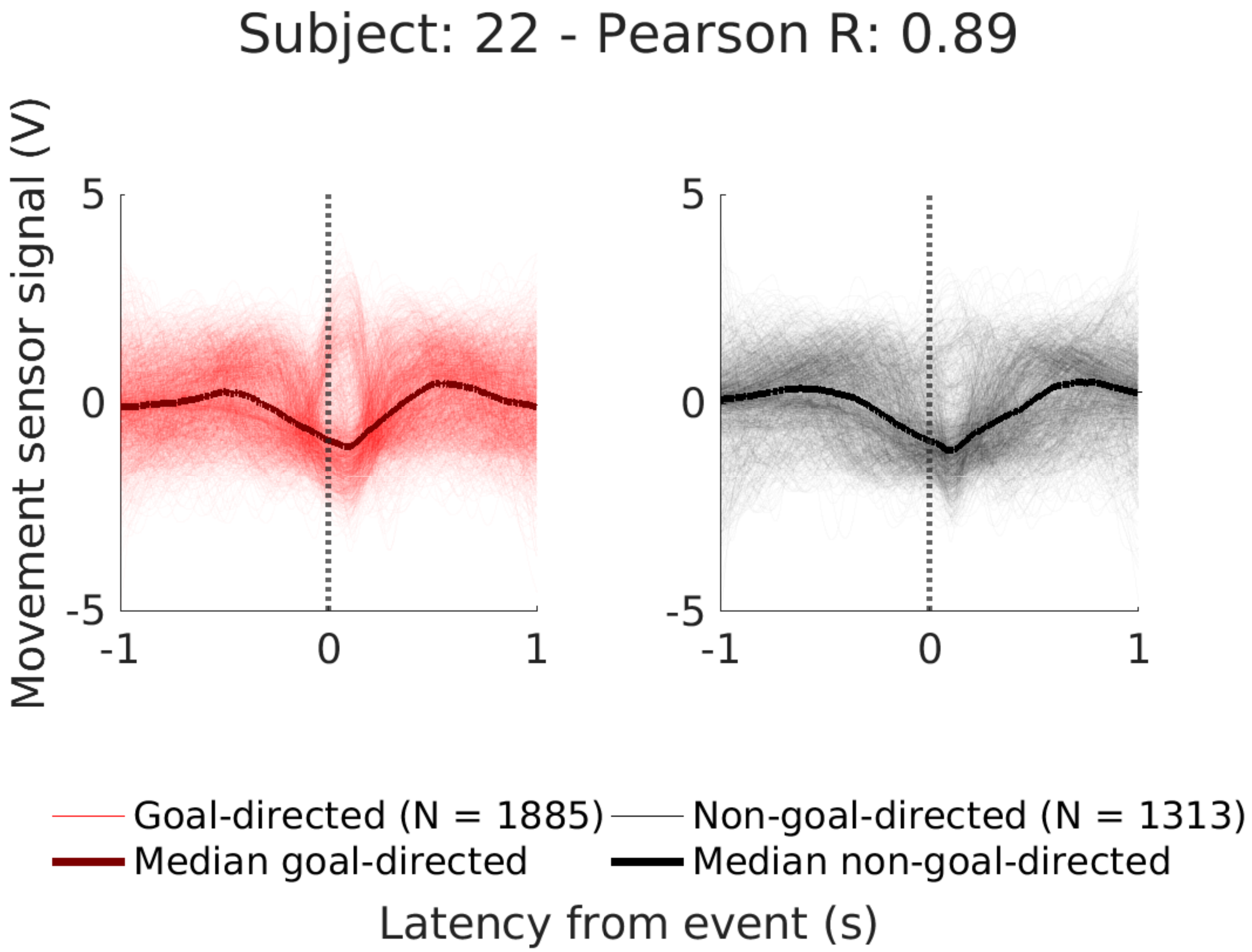
Subject: 22 - Pearson R: 0.89
Movement sensor signal (V)
5
0
-5
-1
0
1
5
0
-5
-1
0
1
Goal-directed (N = 1885)
Non-goal-directed (N = 1313)
Median goal-directed
Median non-goal-directed
Latency from event (s)

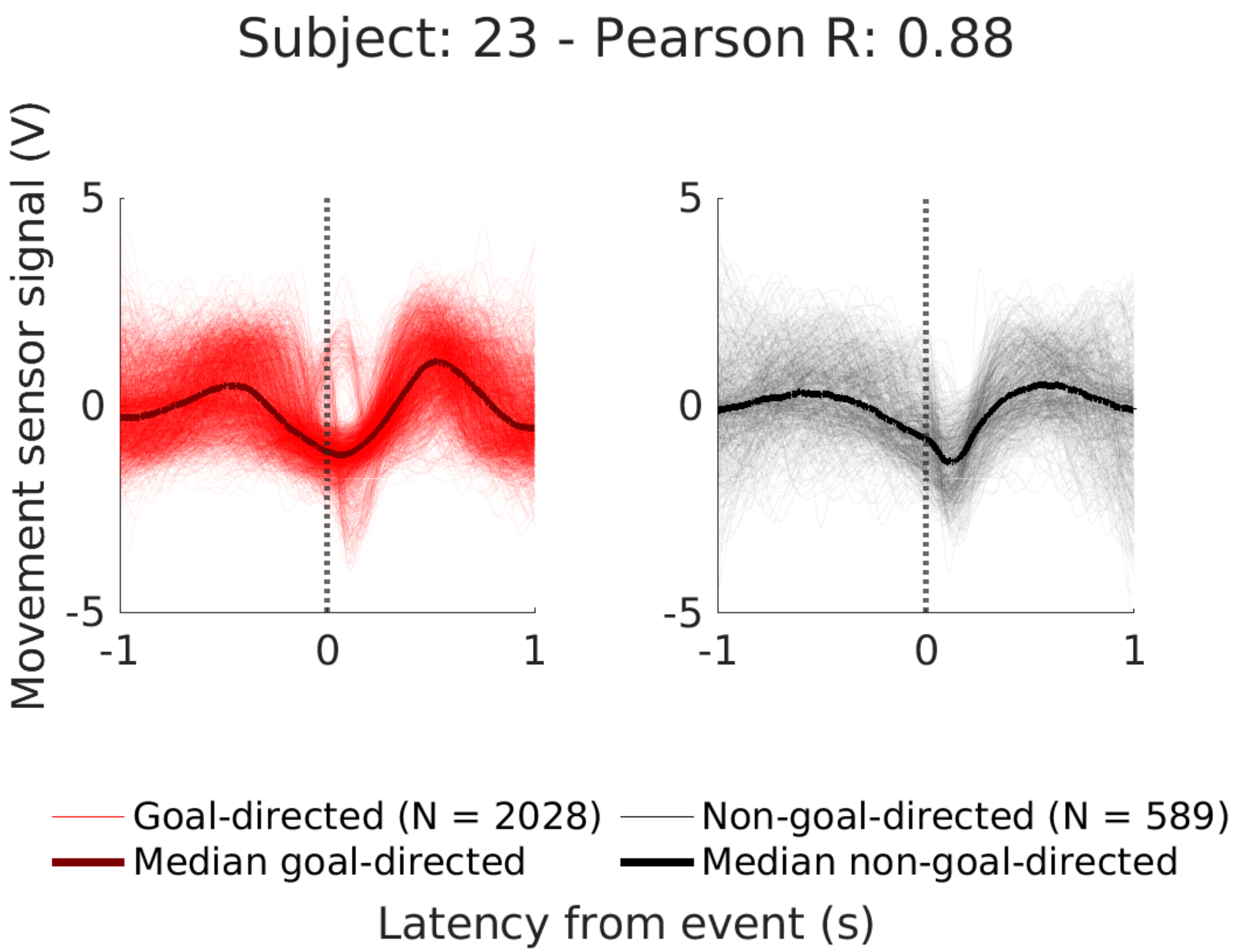

Subject: 23 - Pearson R: 0.88
Movement sensor signal (V)
5
0
-5
-1
0
1
5
0
-5
-1
0
1
Goal-directed (N = 2028)
Median goal-directed
Non-goal-directed (N = 589)
Median non-goal-directed
Latency from event (s)

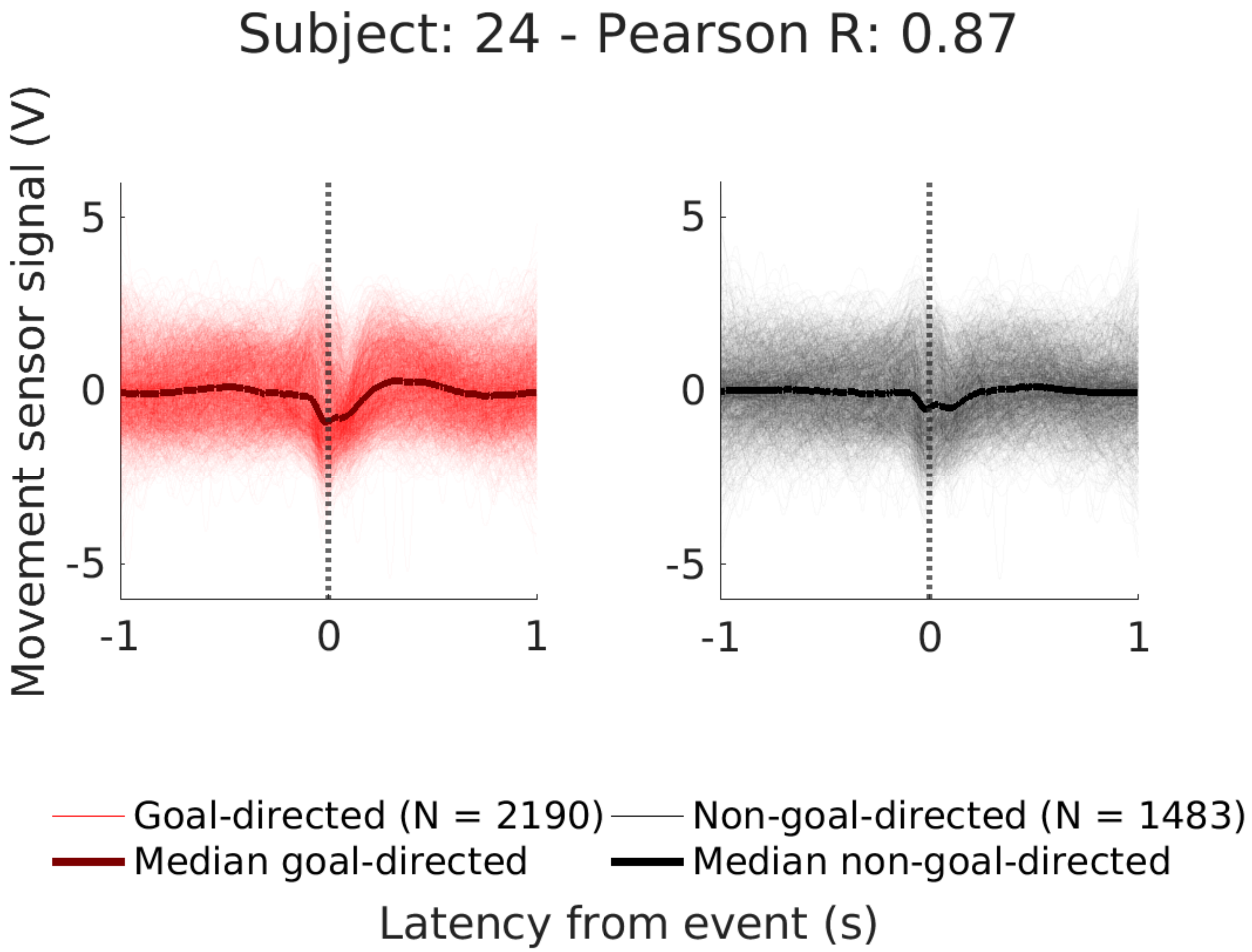

Subject: 24 - Pearson R: 0.87
Movement sensor signal (V)
5
0
-5
-1
0
1
5
0
-5
-1
0
1
Goal-directed (N = 2190)
Non-goal-directed (N = 1483)
Median goal-directed
Median non-goal-directed
Latency from event (s)

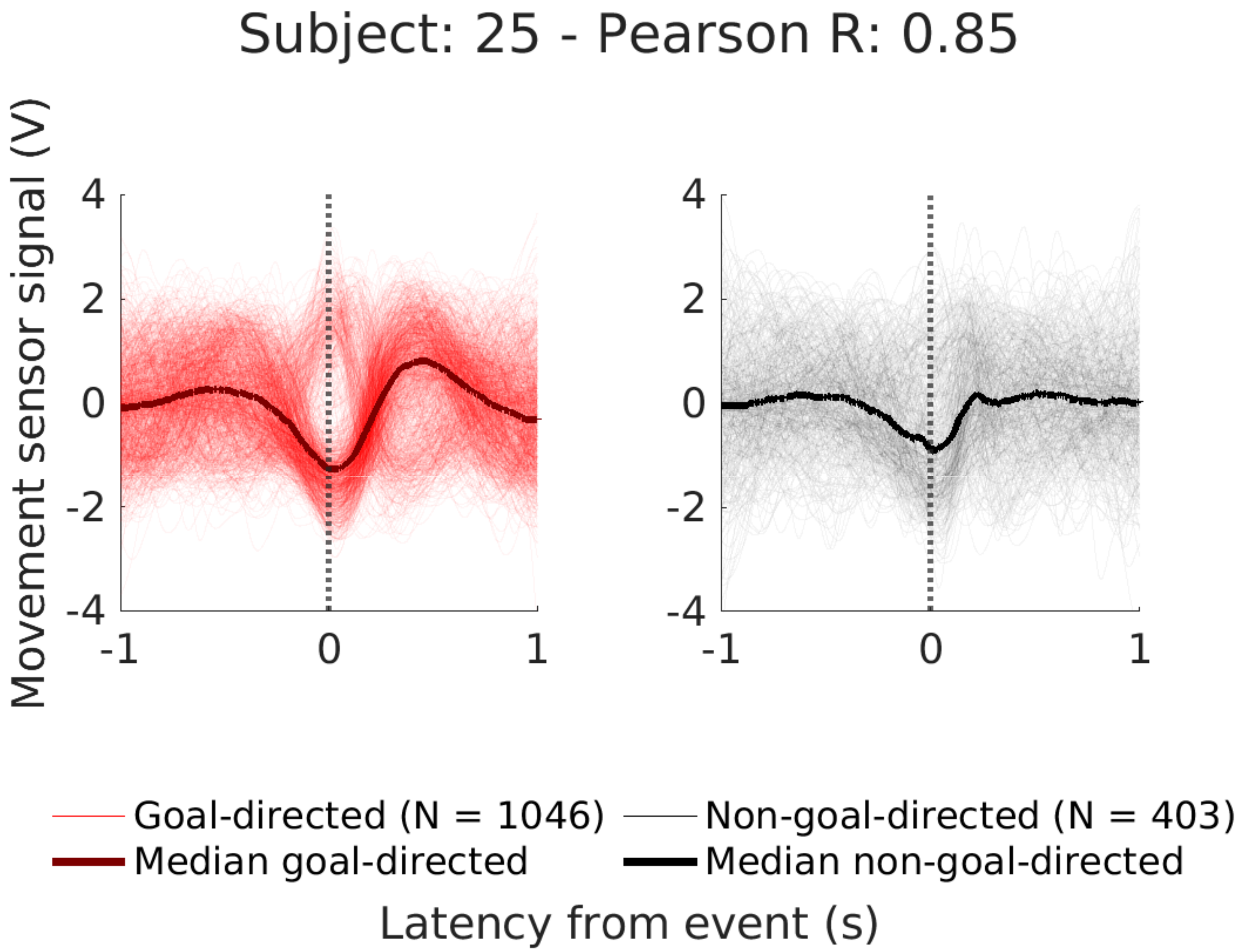

Subject: 25 - Pearson R: 0.85
Movement sensor signal (V)
4
2
0
-2
-4
-1
0
1
Goal-directed (N = 1046)
Median goal-directed
Non-goal-directed (N = 403)
Median non-goal-directed
Latency from event (s)

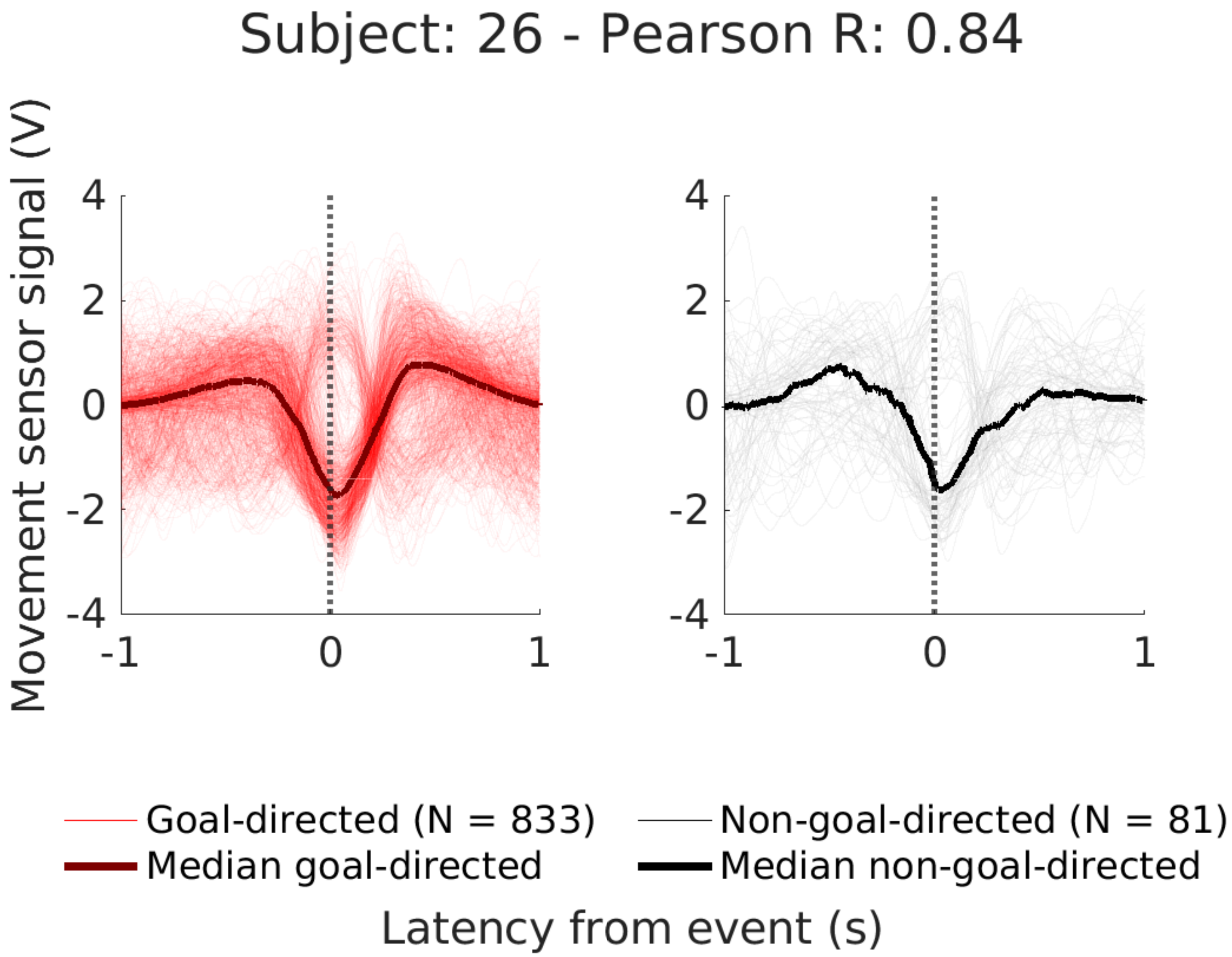
Subject: 26 - Pearson R: 0.84
Movement sensor signal (V)
4
2
0
-2
-4
-1
0
1
Goal-directed (N = 833)
Median goal-directed
Non-goal-directed (N = 81)
Median non-goal-directed
Latency from event (s)

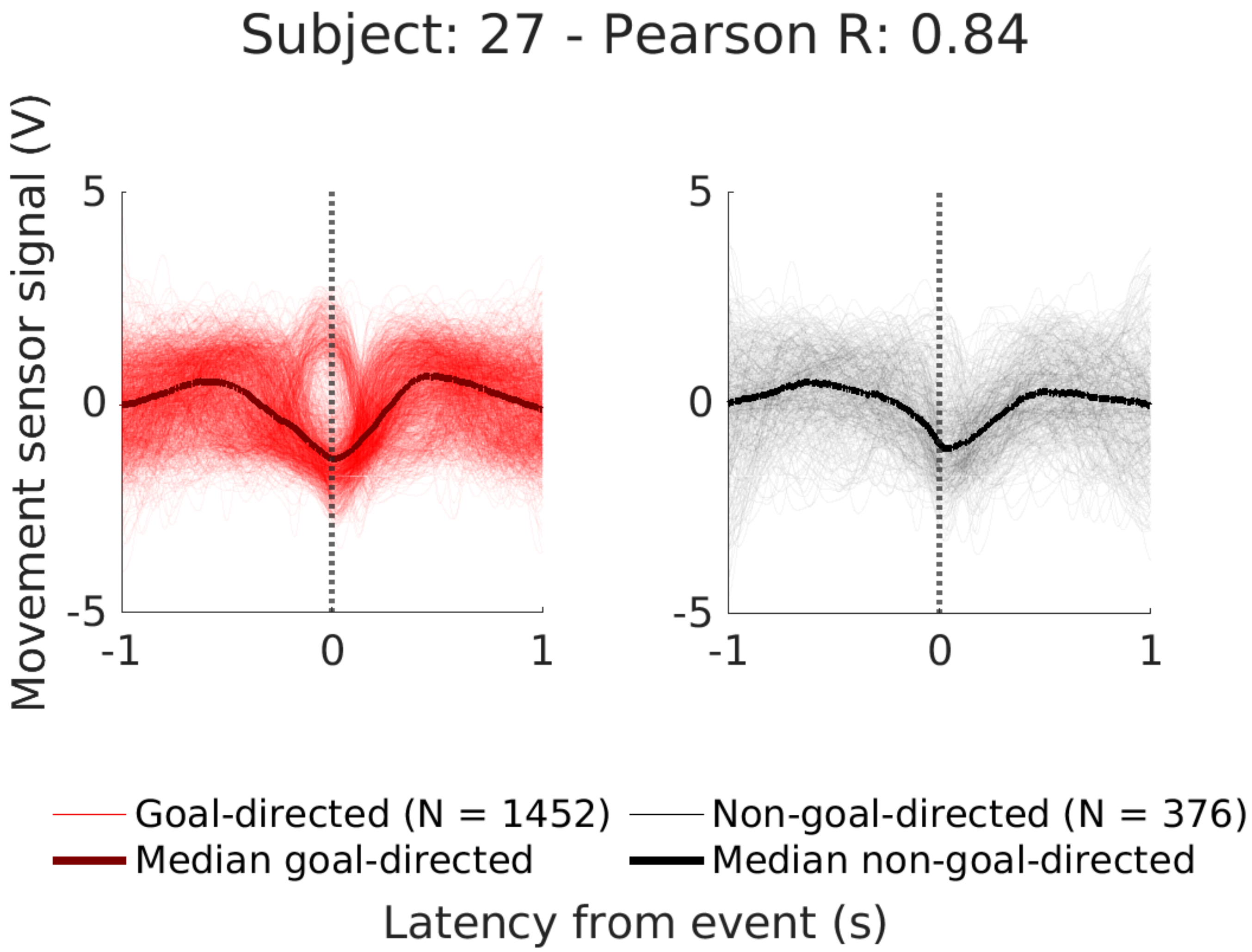
Subject: 27 - Pearson R: 0.84
Movement sensor signal (V)
5
0
-5
-1
0
1
Goal-directed (N = 1452)
Median goal-directed
Non-goal-directed (N = 376)
Median non-goal-directed
Latency from event (s)

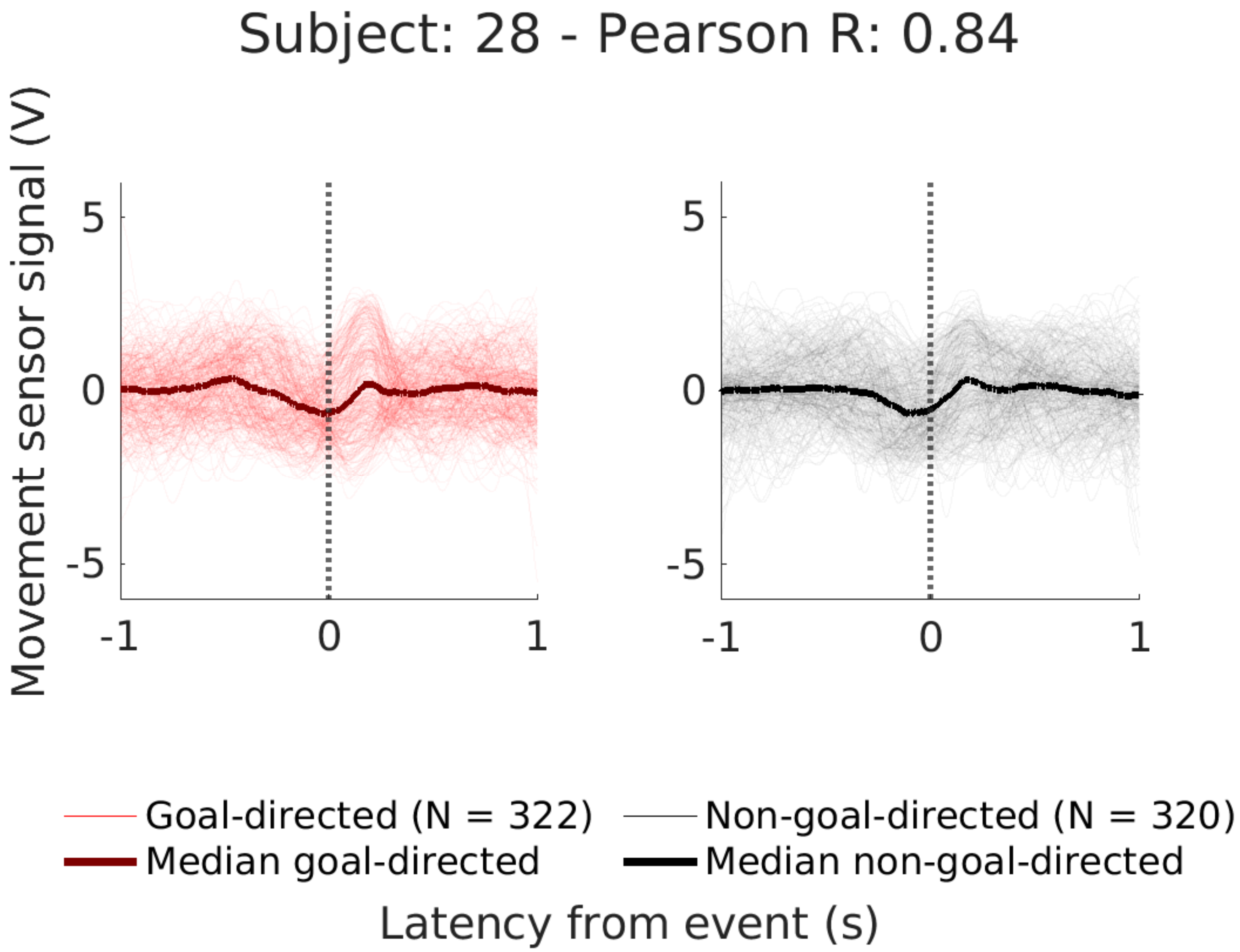
Subject: 28 - Pearson R: 0.84
Movement sensor signal (V)
5
0
-5
-1
0
1
Goal-directed (N = 322)
Median goal-directed
Non-goal-directed (N = 320)
Median non-goal-directed
Latency from event (s)

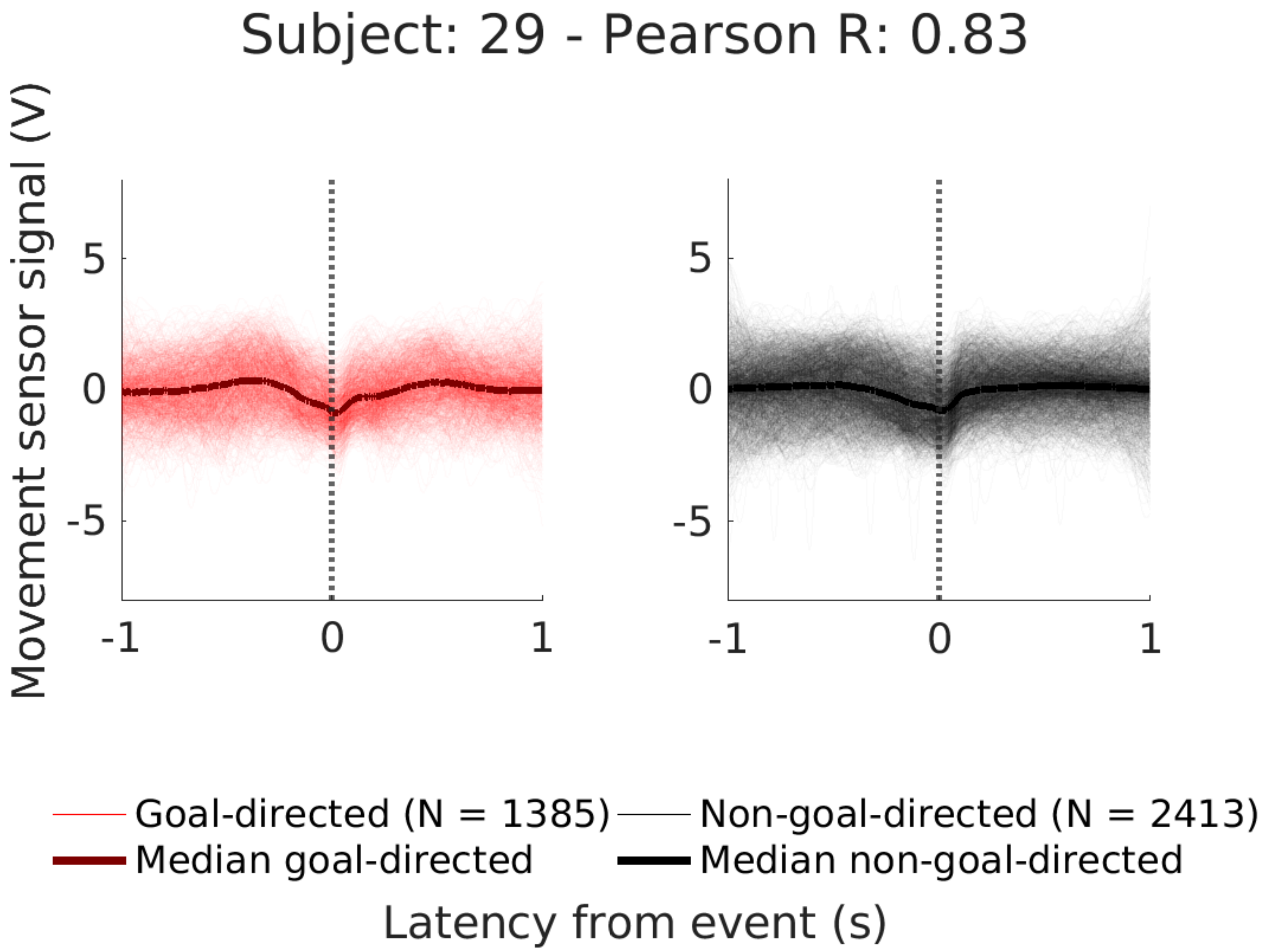

Subject: 29 - Pearson R: 0.83
Movement sensor signal (V)
5
0
-5
-1
0
1
Goal-directed (N = 1385)
Non-goal-directed (N = 2413)
Median goal-directed
Median non-goal-directed
Latency from event (s)

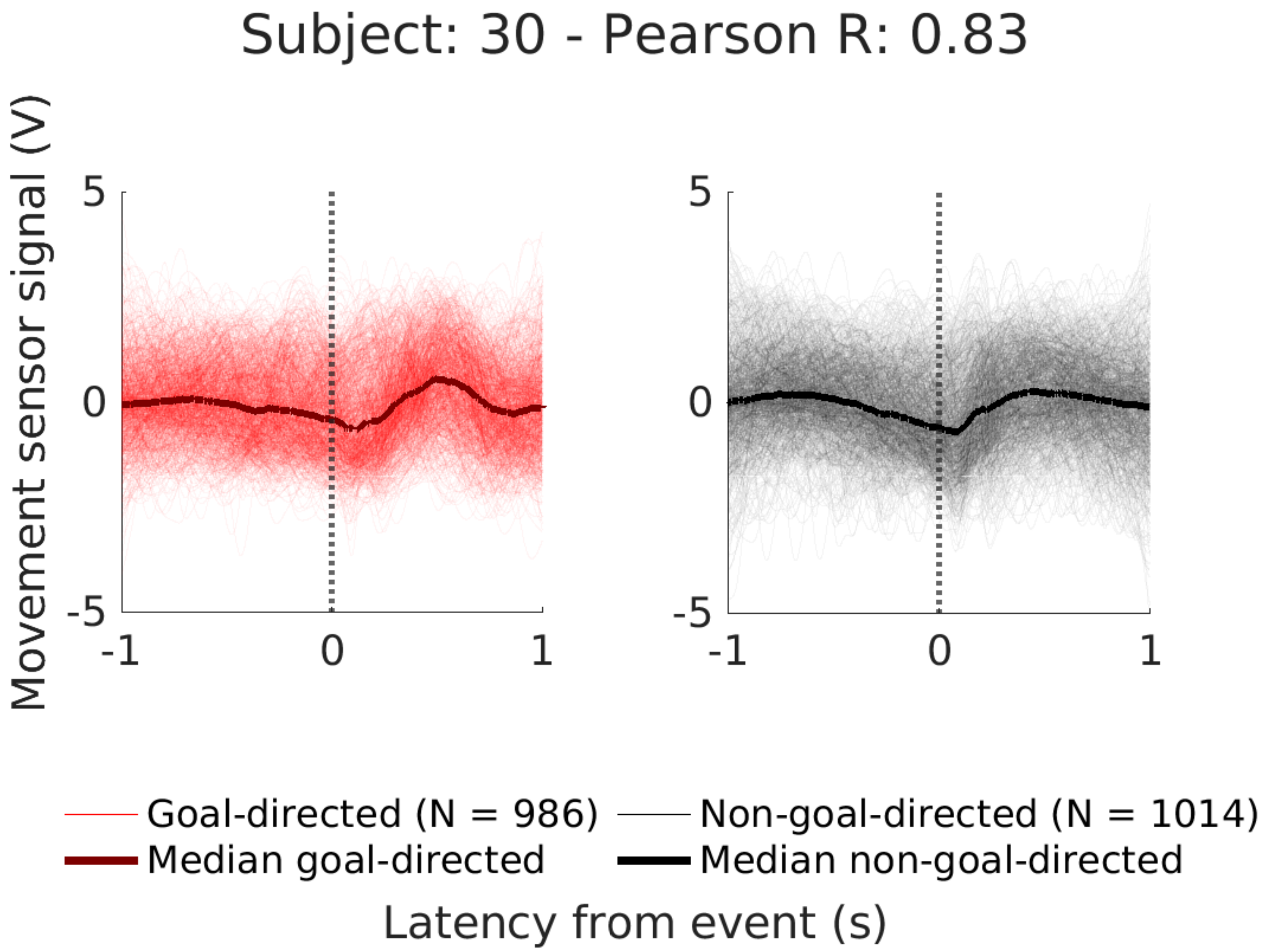
Subject: 30 - Pearson R: 0.83
Movement sensor signal (V)
5
0
-5
-1
0
1
5
0
-5
-1
0
1
Goal-directed (N = 986)
Non-goal-directed (N = 1014)
Median goal-directed
Median non-goal-directed
Latency from event (s)

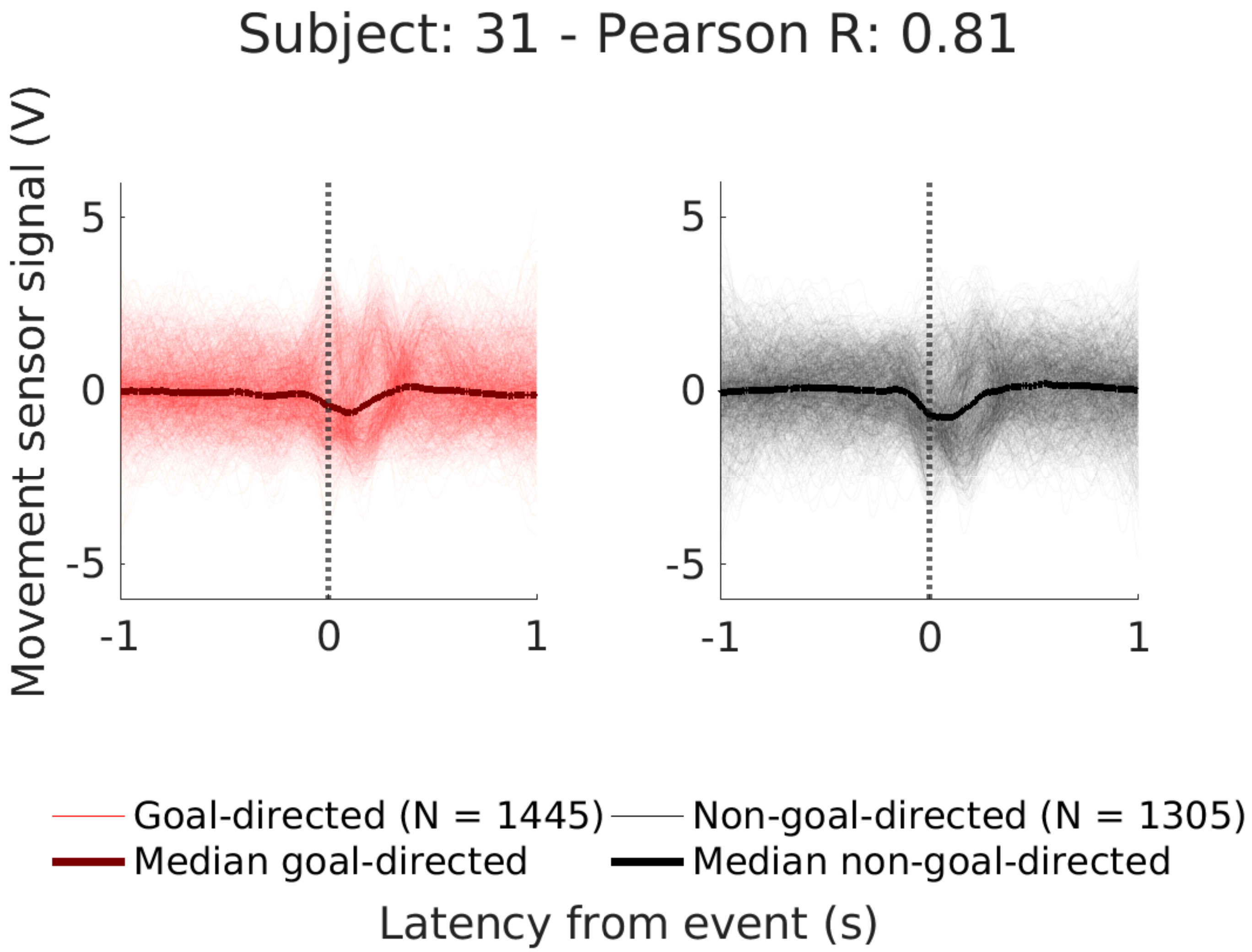
Subject: 31 - Pearson R: 0.81
Movement sensor signal (V)
5
0
-5
-1
0
1
Goal-directed (N = 1445)
Non-goal-directed (N = 1305)
Median goal-directed
Median non-goal-directed
Latency from event (s)

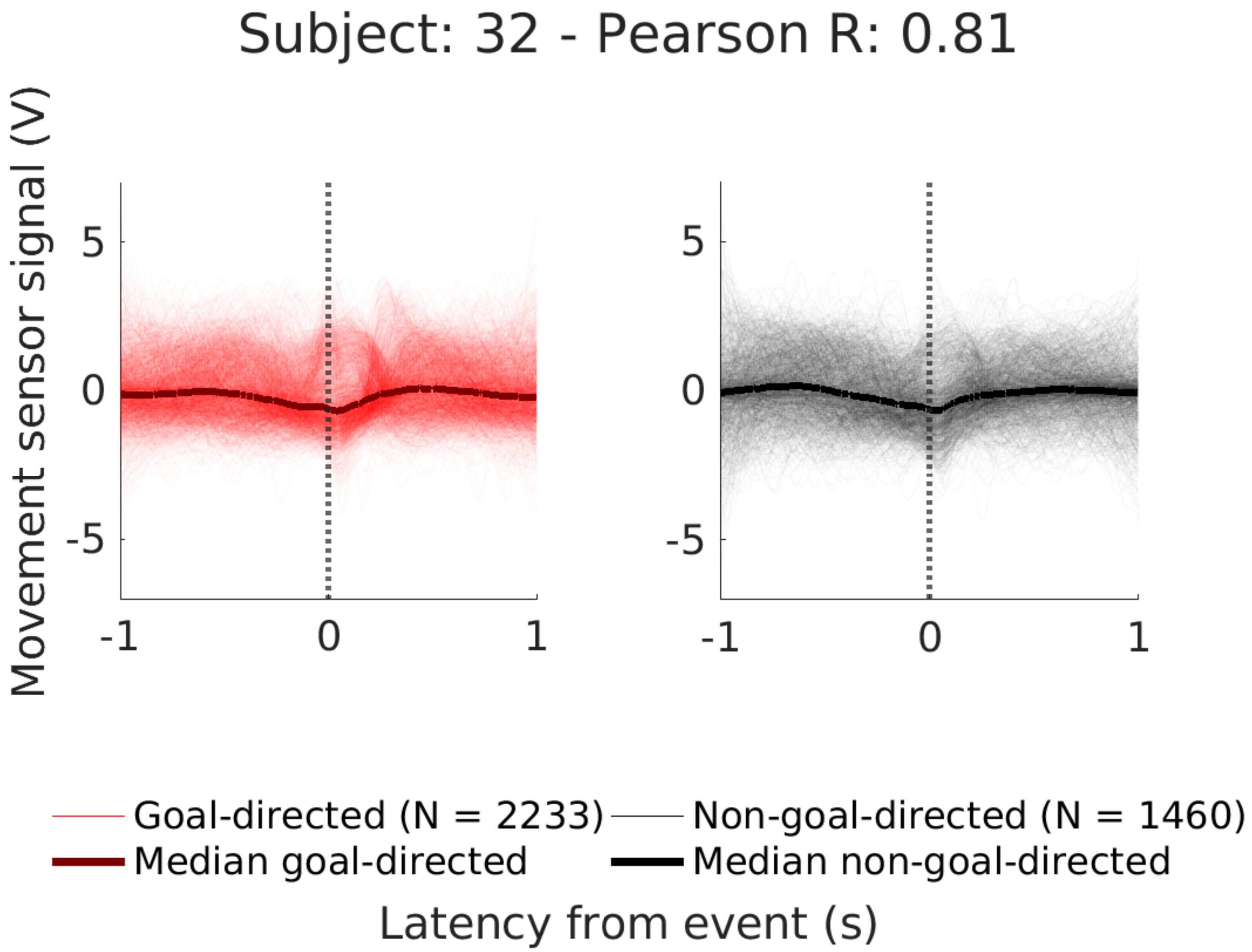
Subject: 32 - Pearson R: 0.81
Movement sensor signal (V)
5
0
-5
-1
0
1
5
0
-5
-1
0
1
Goal-directed (N = 2233)
Non-goal-directed (N = 1460)
Median goal-directed
Median non-goal-directed
Latency from event (s)

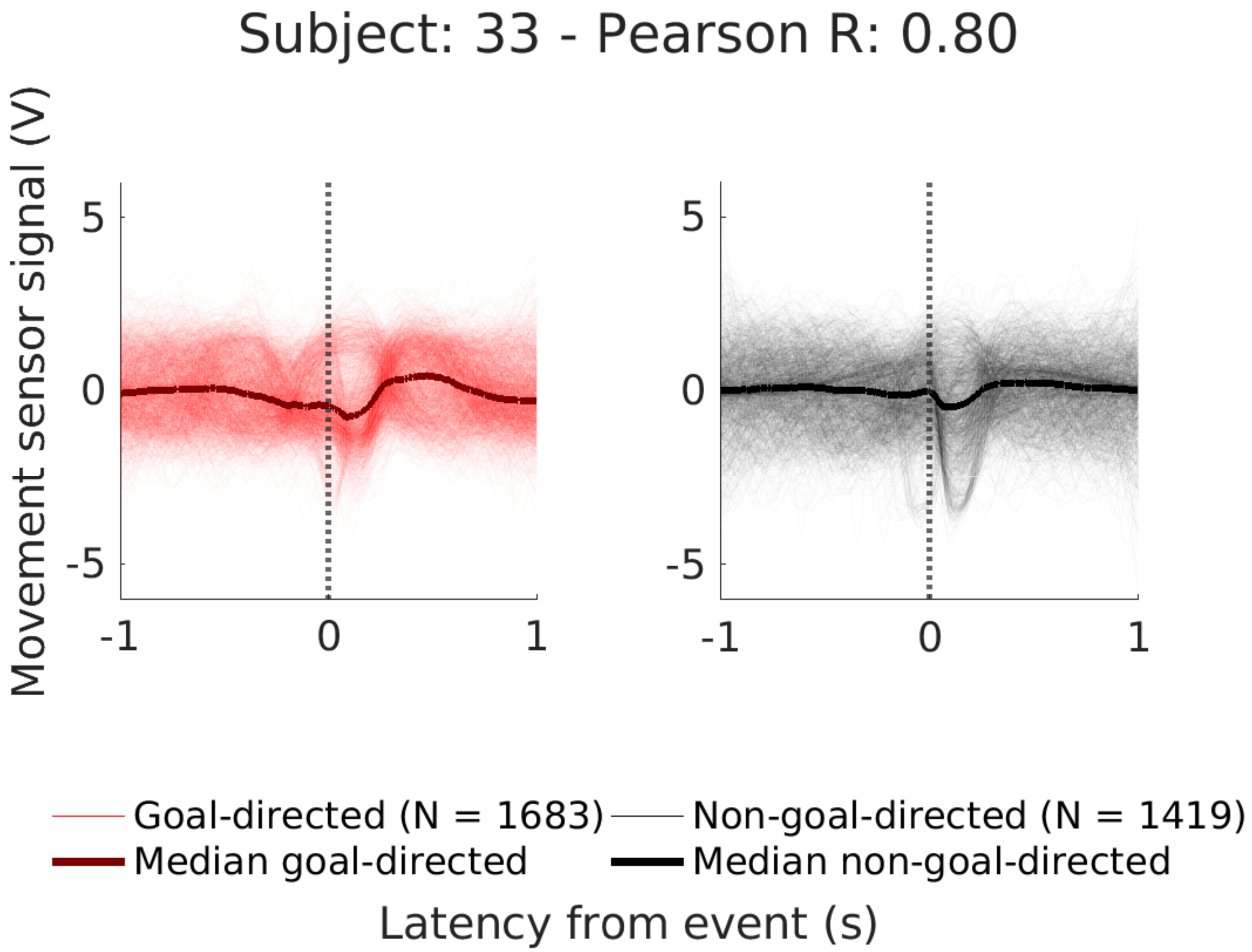
Subject: 33 - Pearson R: 0.80
Movement sensor signal (V)
5
0
-5
-1
0
1
Goal-directed (N = 1683)
Non-goal-directed (N = 1419)
Median goal-directed
Median non-goal-directed
Latency from event (s)

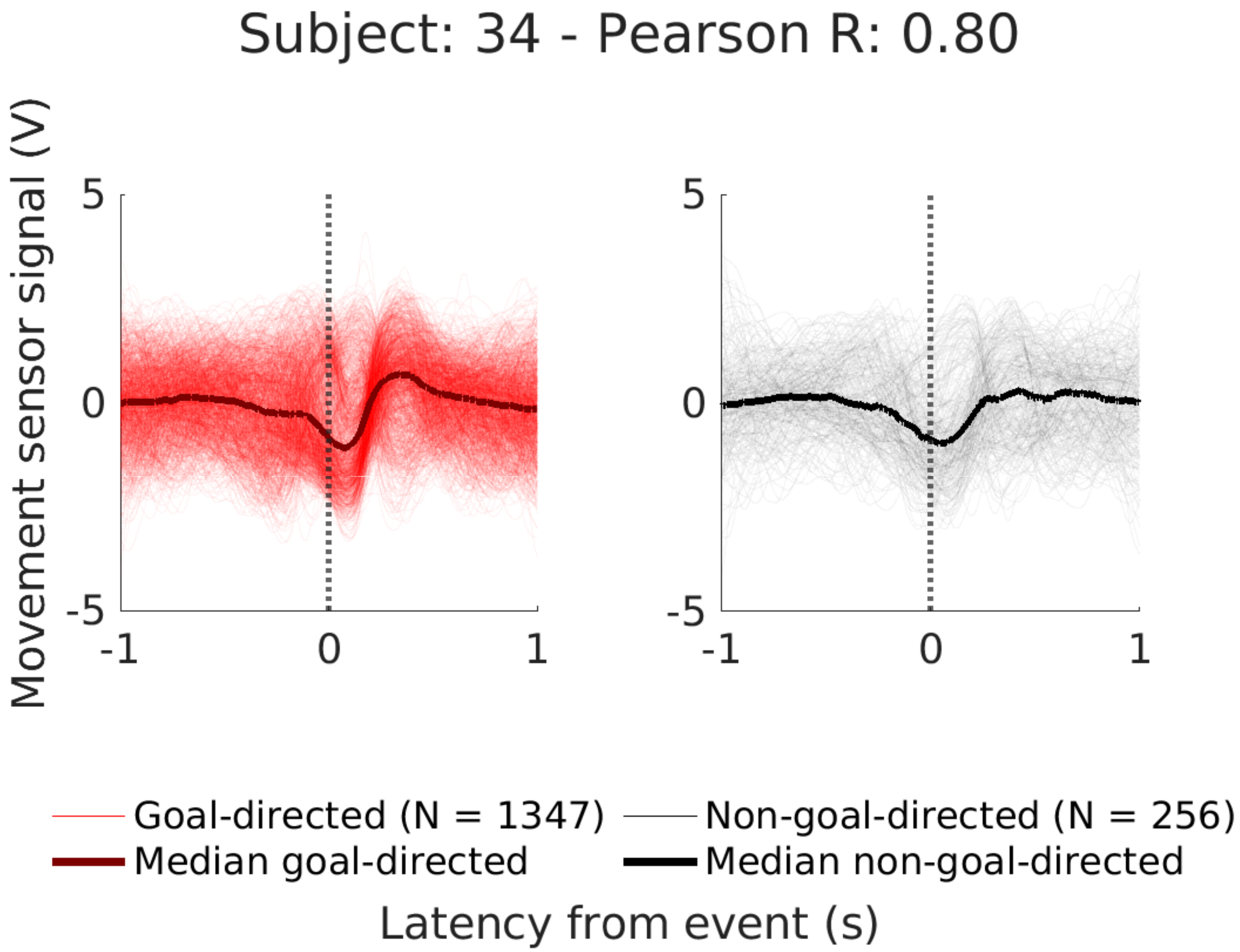

Subject: 34 - Pearson R: 0.80
Movement sensor signal (V)
5
0
-5
-1
0
1
5
0
-5
-1
0
1
Goal-directed (N = 1347)
Non-goal-directed (N = 256)
Median goal-directed
Median non-goal-directed
Latency from event (s)

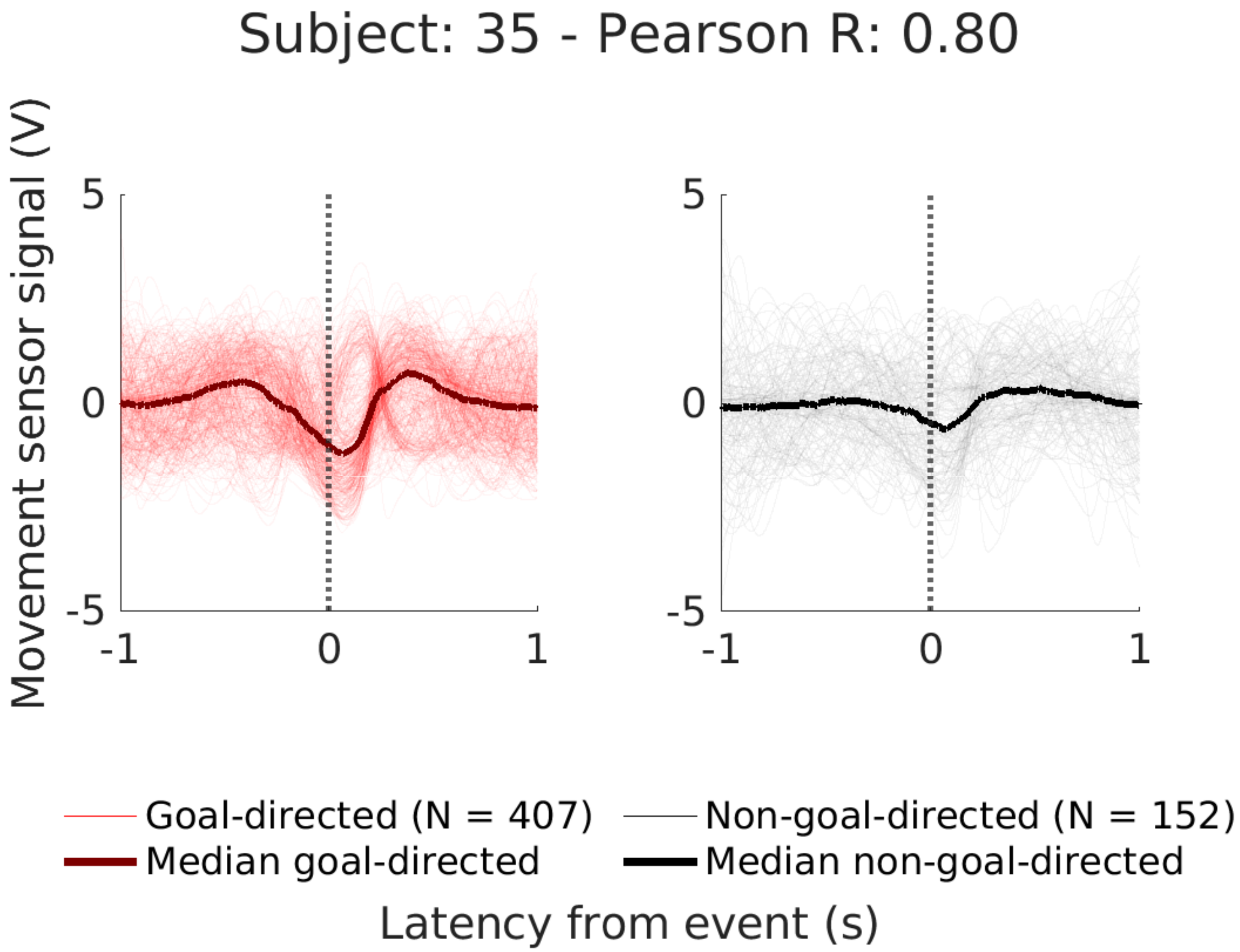

Subject: 35 - Pearson R: 0.80
Movement sensor signal (V)
5
0
-5
-1
0
1
Goal-directed (N = 407)
Median goal-directed
Non-goal-directed (N = 152)
Median non-goal-directed
Latency from event (s)

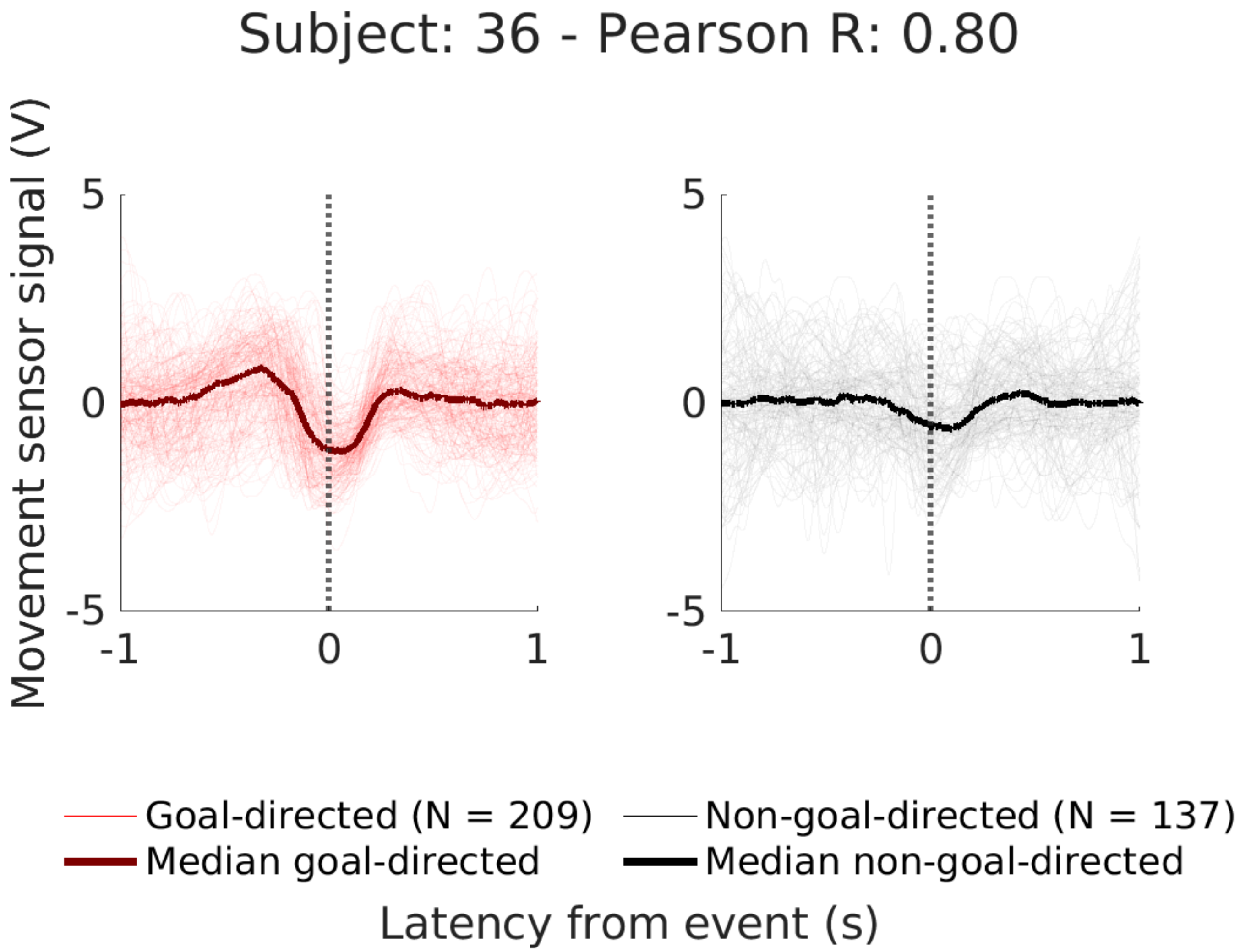
Subject: 36 - Pearson R: 0.80
Movement sensor signal (V)
5
0
-5
-1
0
1
Goal-directed (N = 209)
Non-goal-directed (N = 137)
Median goal-directed
Median non-goal-directed
Latency from event (s)

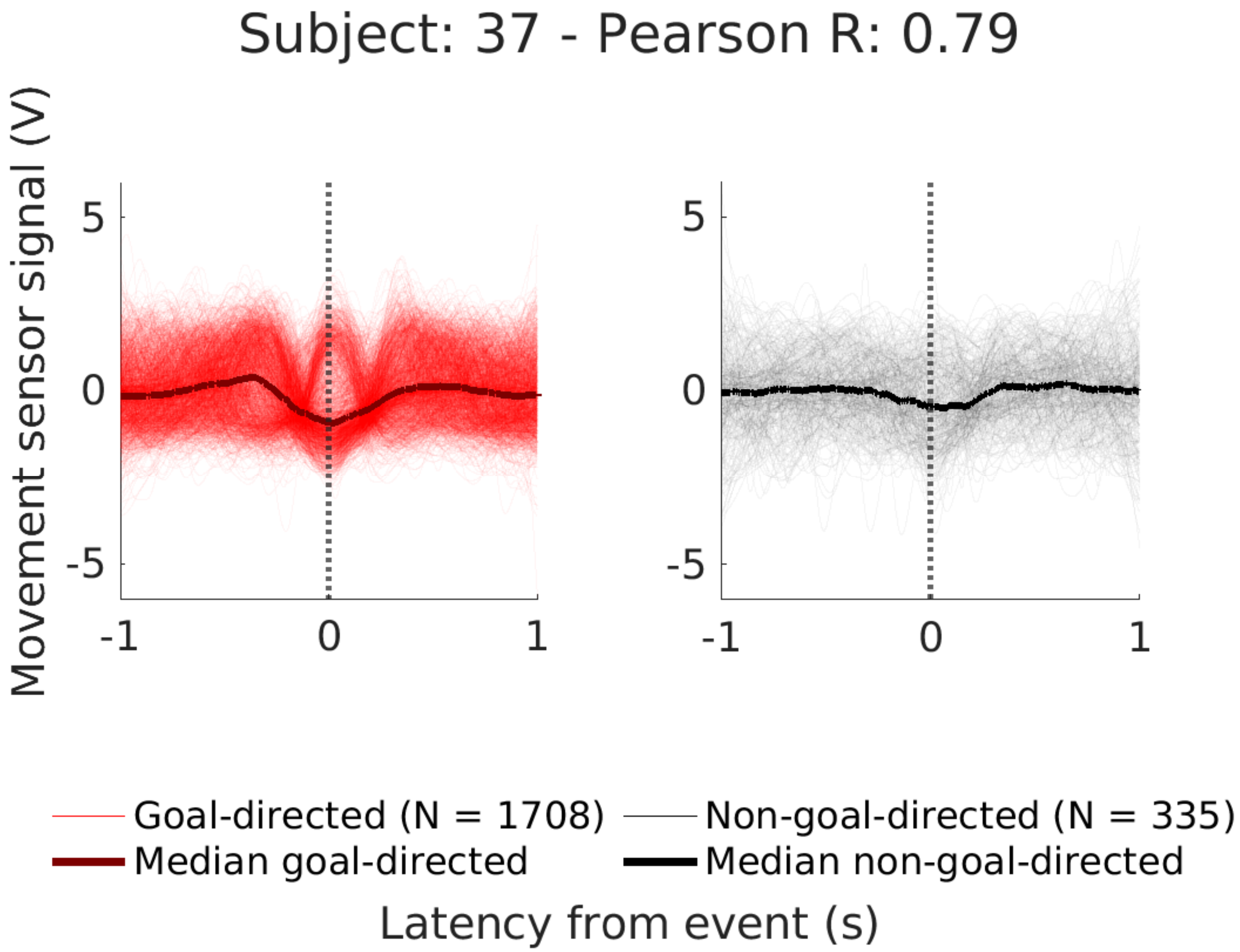
Subject: 37 - Pearson R: 0.79
Movement sensor signal (V)
5
0
-5
-1
0
1
5
0
-5
-1
0
1
Goal-directed (N = 1708)
Non-goal-directed (N = 335)
Median goal-directed
Median non-goal-directed
Latency from event (s)

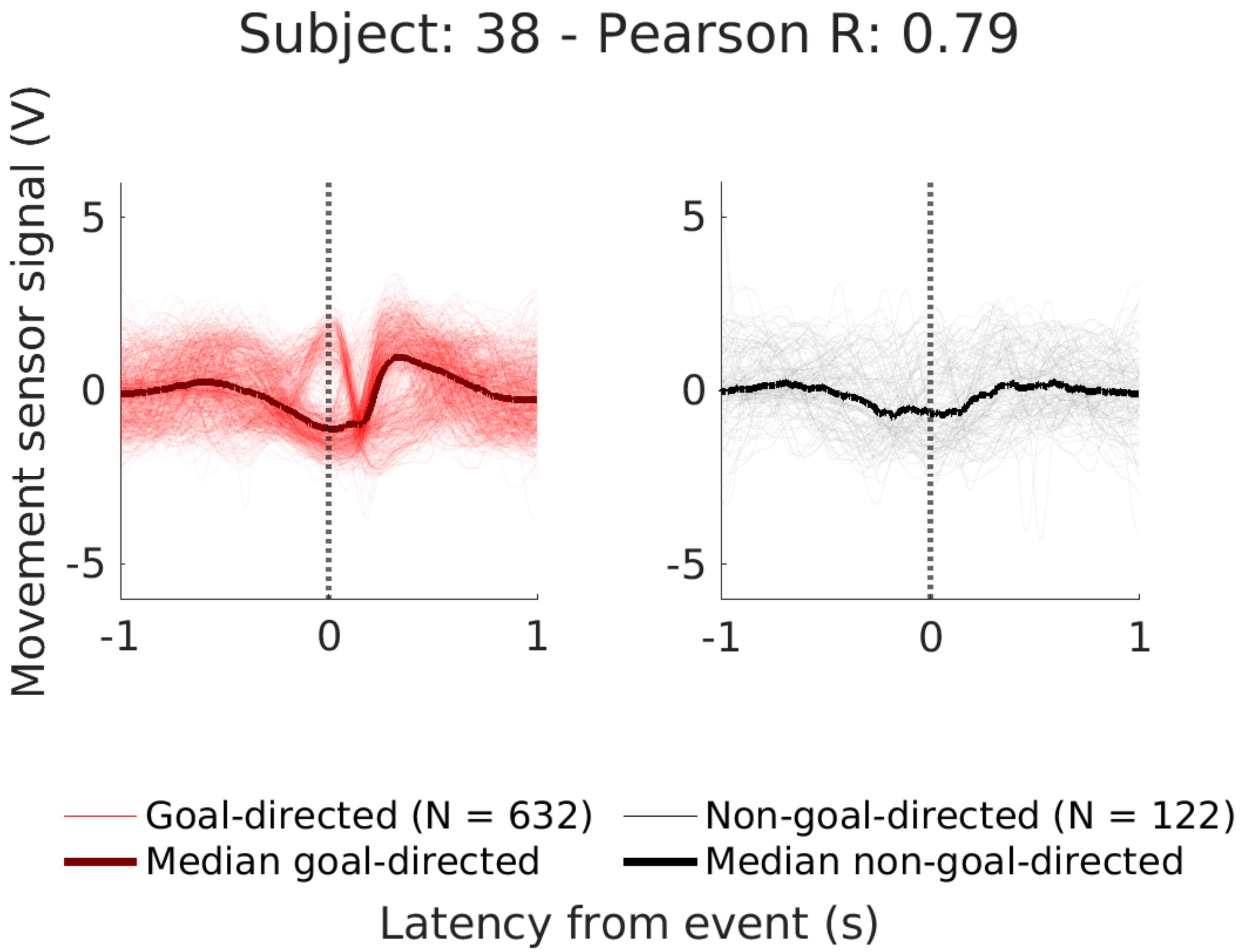
Subject: 38 - Pearson R: 0.79
Movement sensor signal (V)
5
0
-5
-1
0
1
Goal-directed (N = 632)
Median goal-directed
Non-goal-directed (N = 122)
Median non-goal-directed
Latency from event (s)

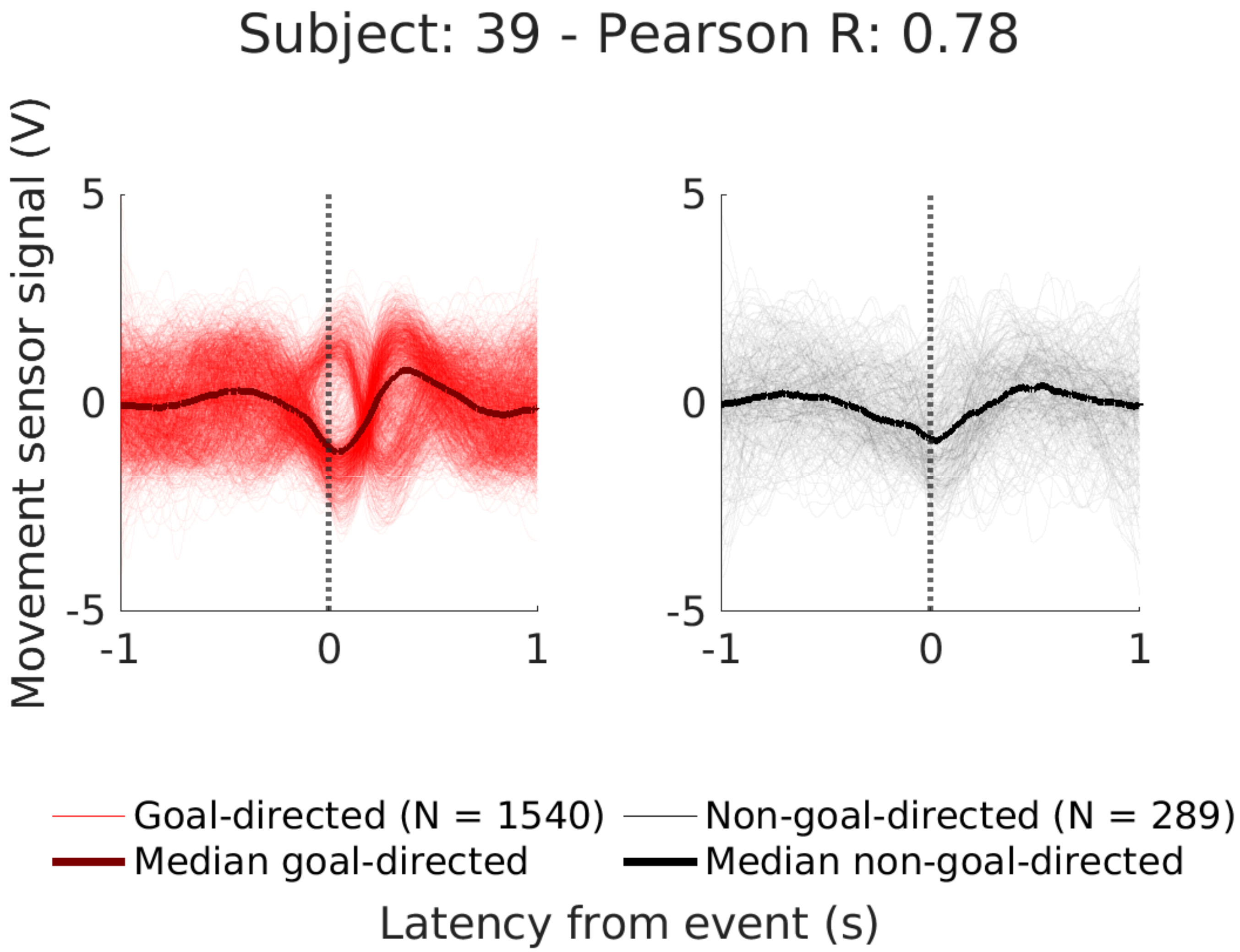

Subject: 39 - Pearson R: 0.78
Movement sensor signal (V)
5
0
-5
-1
0
1
5
0
-5
-1
0
1
Goal-directed (N = 1540)
Median goal-directed
Non-goal-directed (N = 289)
Median non-goal-directed
Latency from event (s)

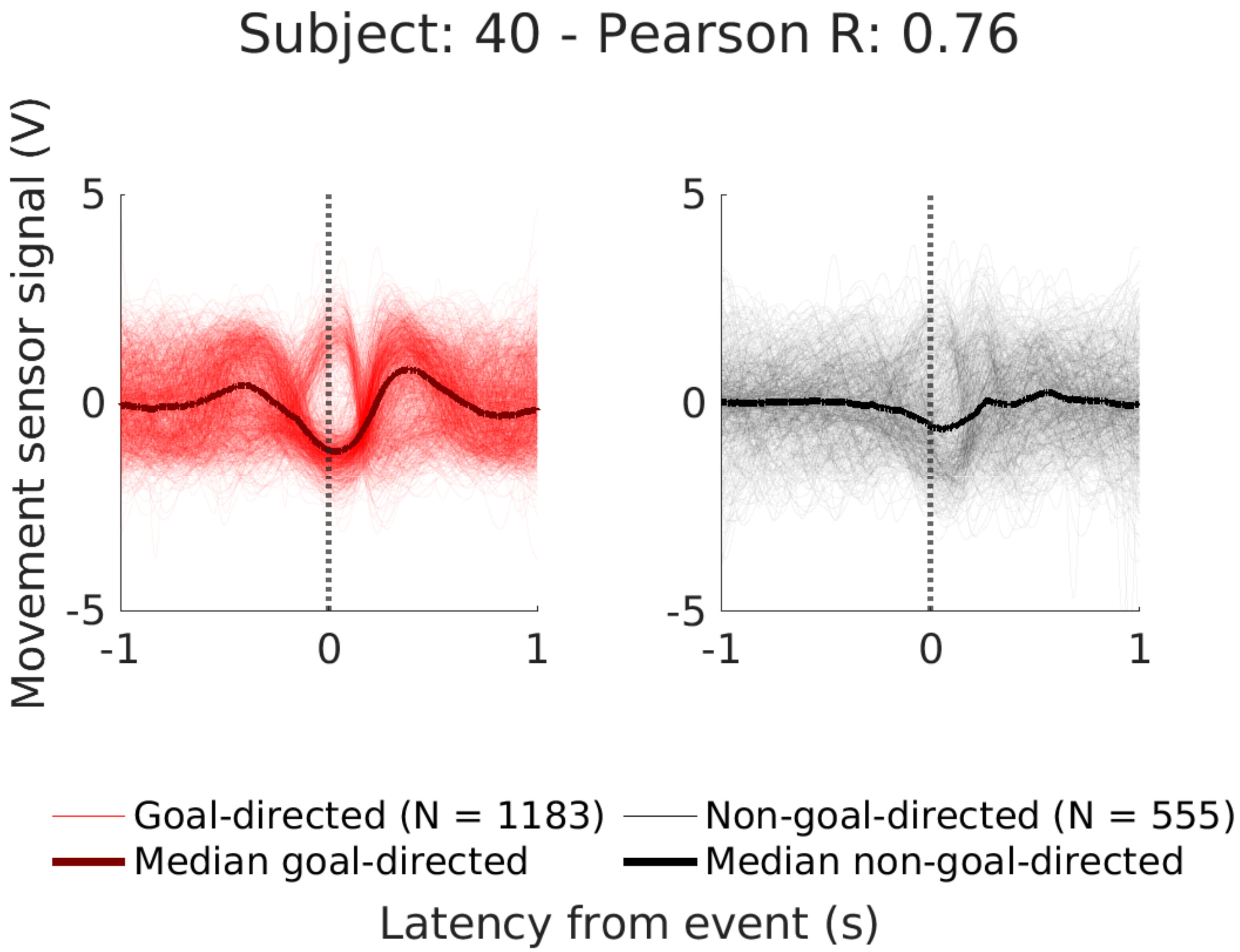
Subject: 40 - Pearson R: 0.76
Movement sensor signal (V)
5
0
-5
-1
0
1
5
0
-5
-1
0
1
Goal-directed (N = 1183)
Median goal-directed
Non-goal-directed (N = 555)
Median non-goal-directed
Latency from event (s)

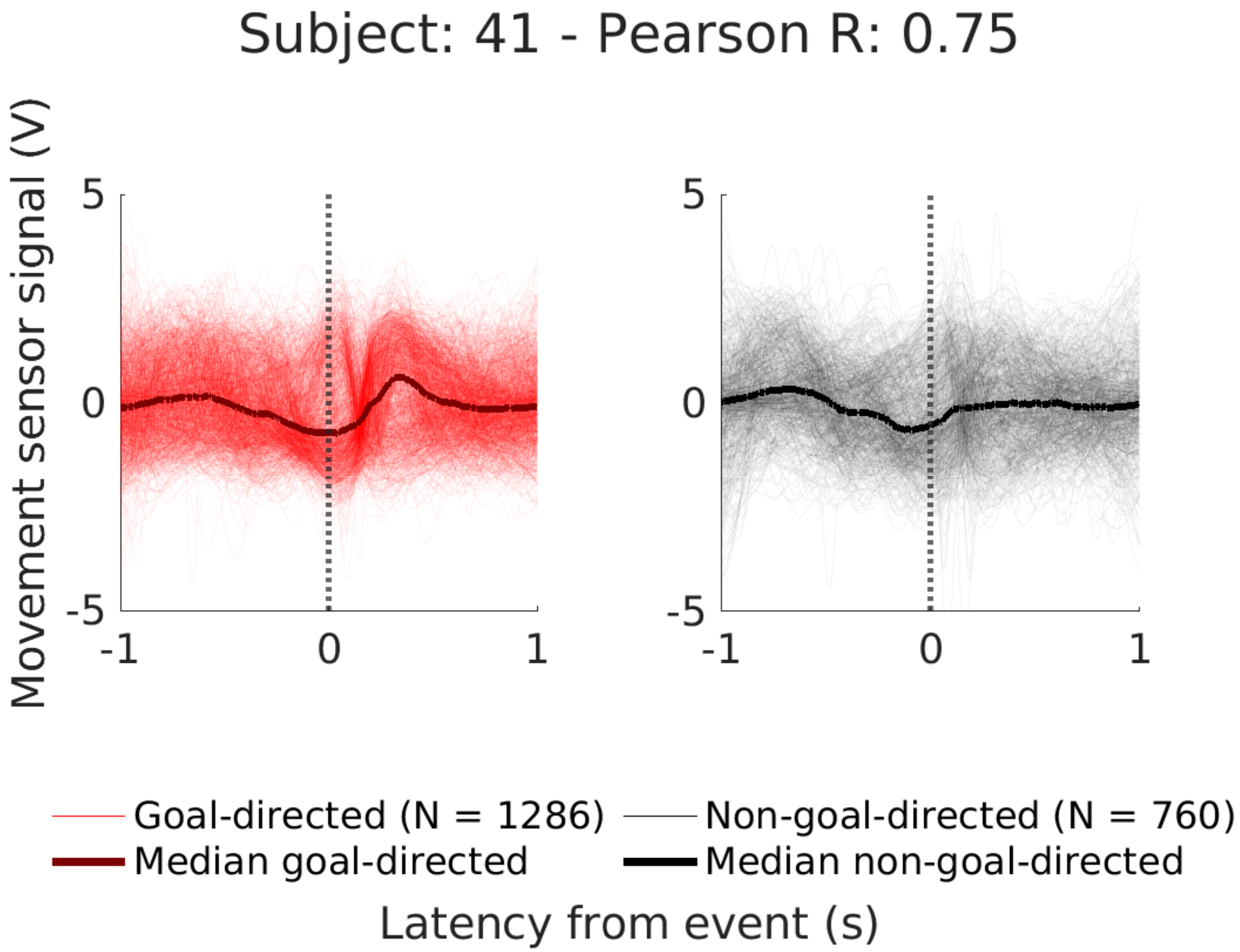
Subject: 41 - Pearson R: 0.75
Movement sensor signal (V)
5
0
-5
-1
0
1
5
0
-5
-1
0
1
Goal-directed (N = 1286)
Median goal-directed
Non-goal-directed (N = 760)
Median non-goal-directed
Latency from event (s)

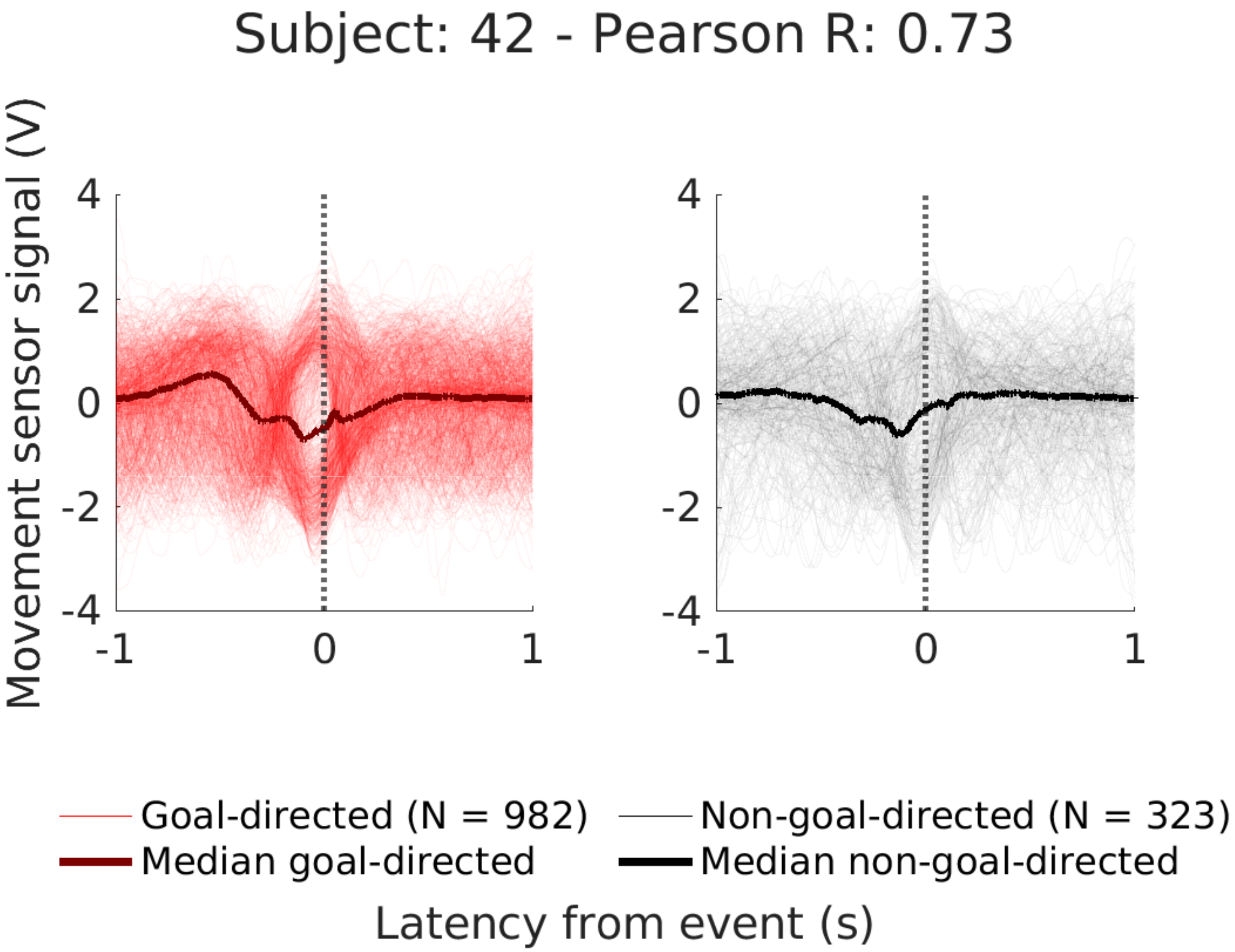
Subject: 42 - Pearson R: 0.73
Movement sensor signal (V)
4
2
0
-2
-4
-1
0
1
Goal-directed (N = 982)
Median goal-directed
Non-goal-directed (N = 323)
Median non-goal-directed
Latency from event (s)

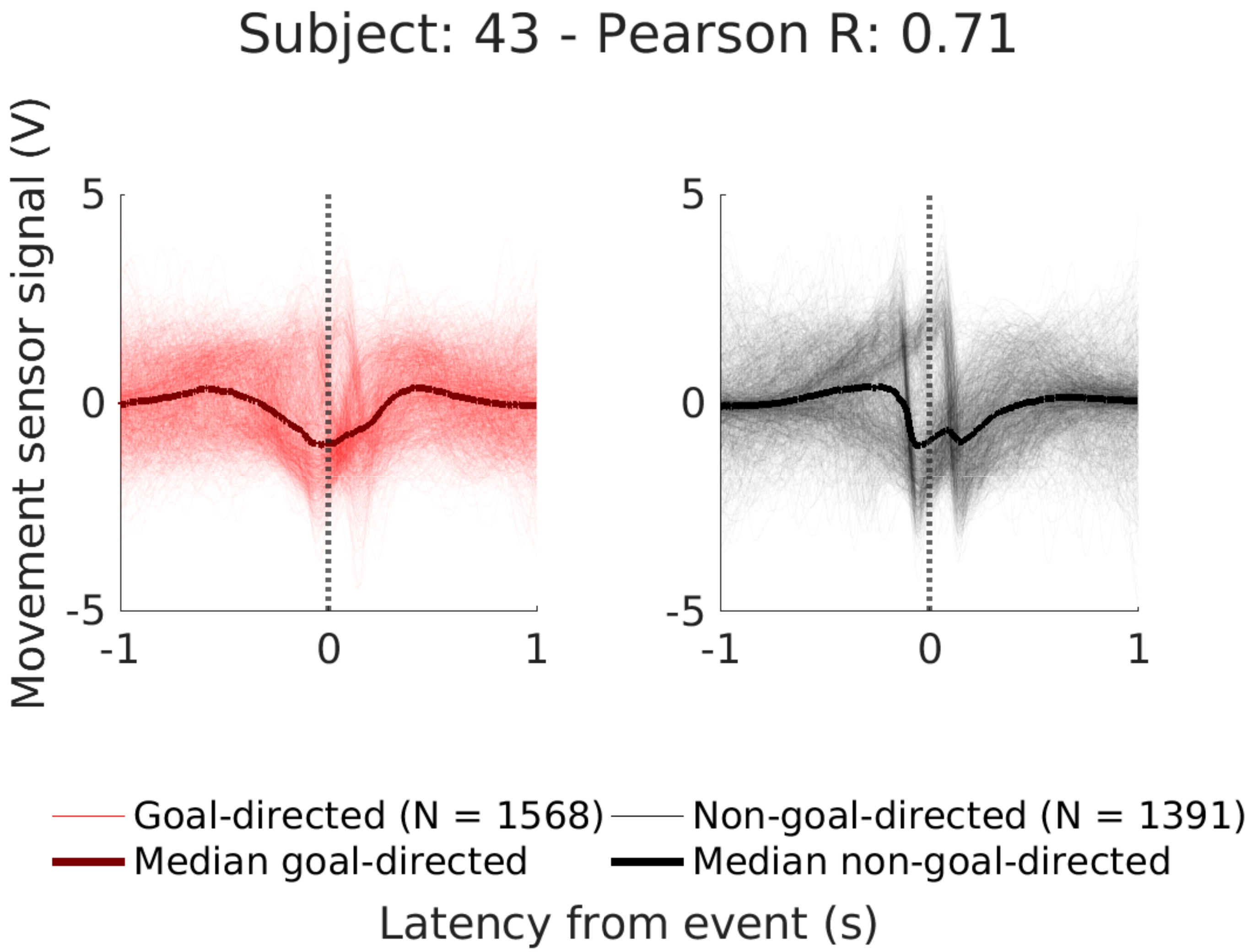
Subject: 43 - Pearson R: 0.71
Movement sensor signal (V)
5
0
-5
-1
0
1
5
0
-5
-1
0
1
Goal-directed (N = 1568)
Non-goal-directed (N = 1391)
Median goal-directed
Median non-goal-directed
Latency from event (s)

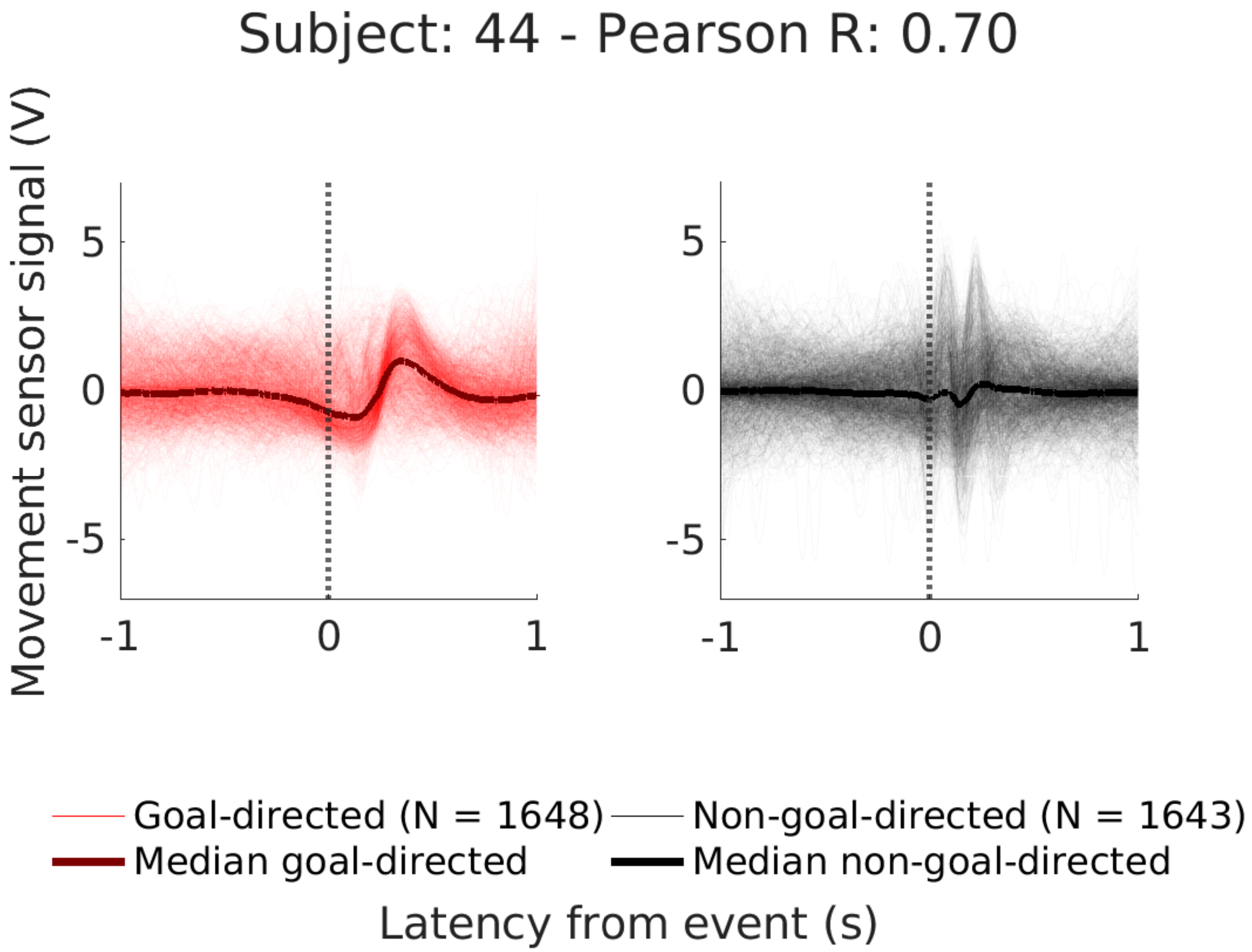
Subject: 44 - Pearson R: 0.70
Movement sensor signal (V)
5
0
-5
-1
0
1
Goal-directed (N = 1648)
Non-goal-directed (N = 1643)
Median goal-directed
Median non-goal-directed
Latency from event (s)

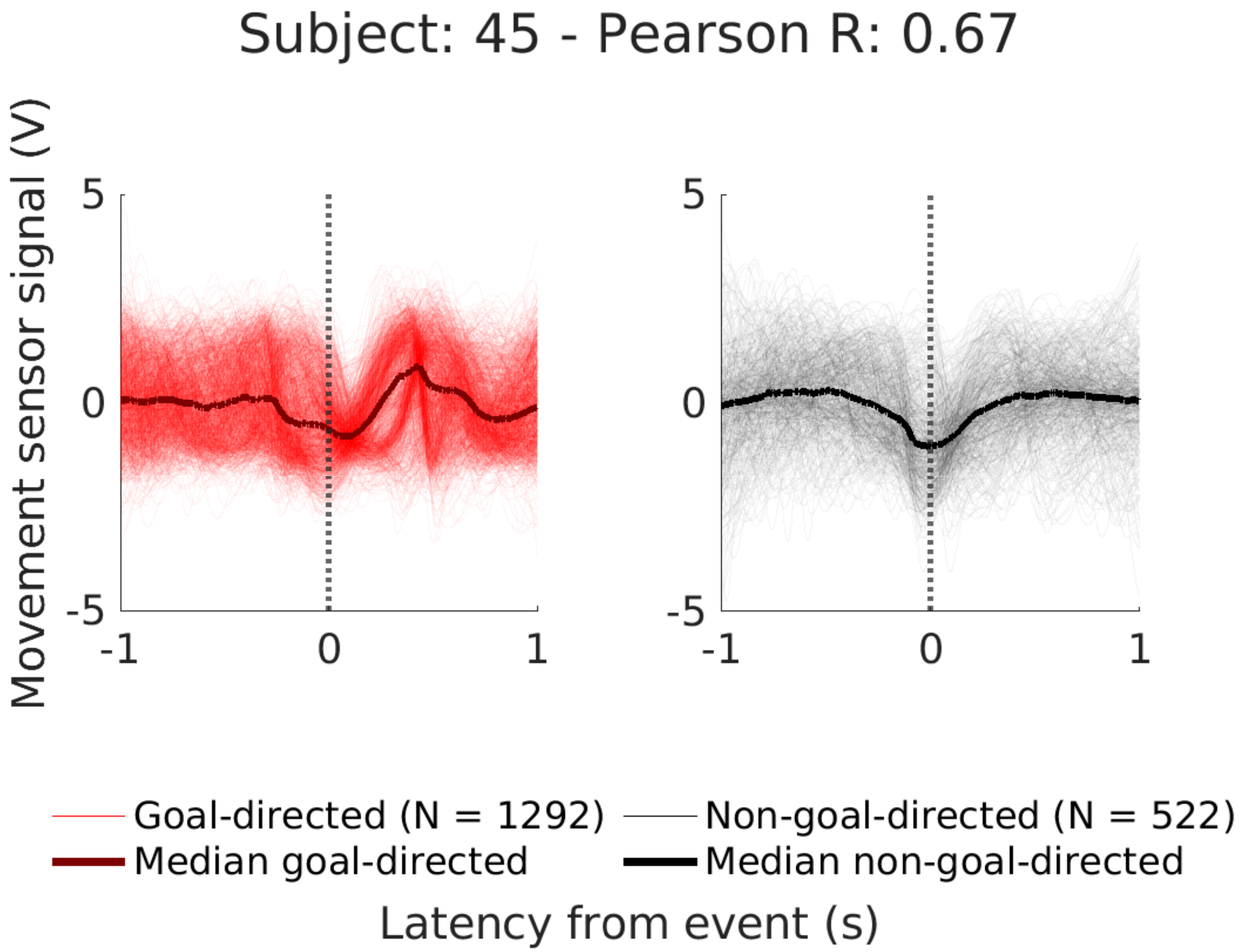

Subject: 45 - Pearson R: 0.67
Movement sensor signal (V)
5
0
-5
-1
0
1
5
0
-5
-1
0
1
Goal-directed (N = 1292)
Median goal-directed
Non-goal-directed (N = 522)
Median non-goal-directed
Latency from event (s)

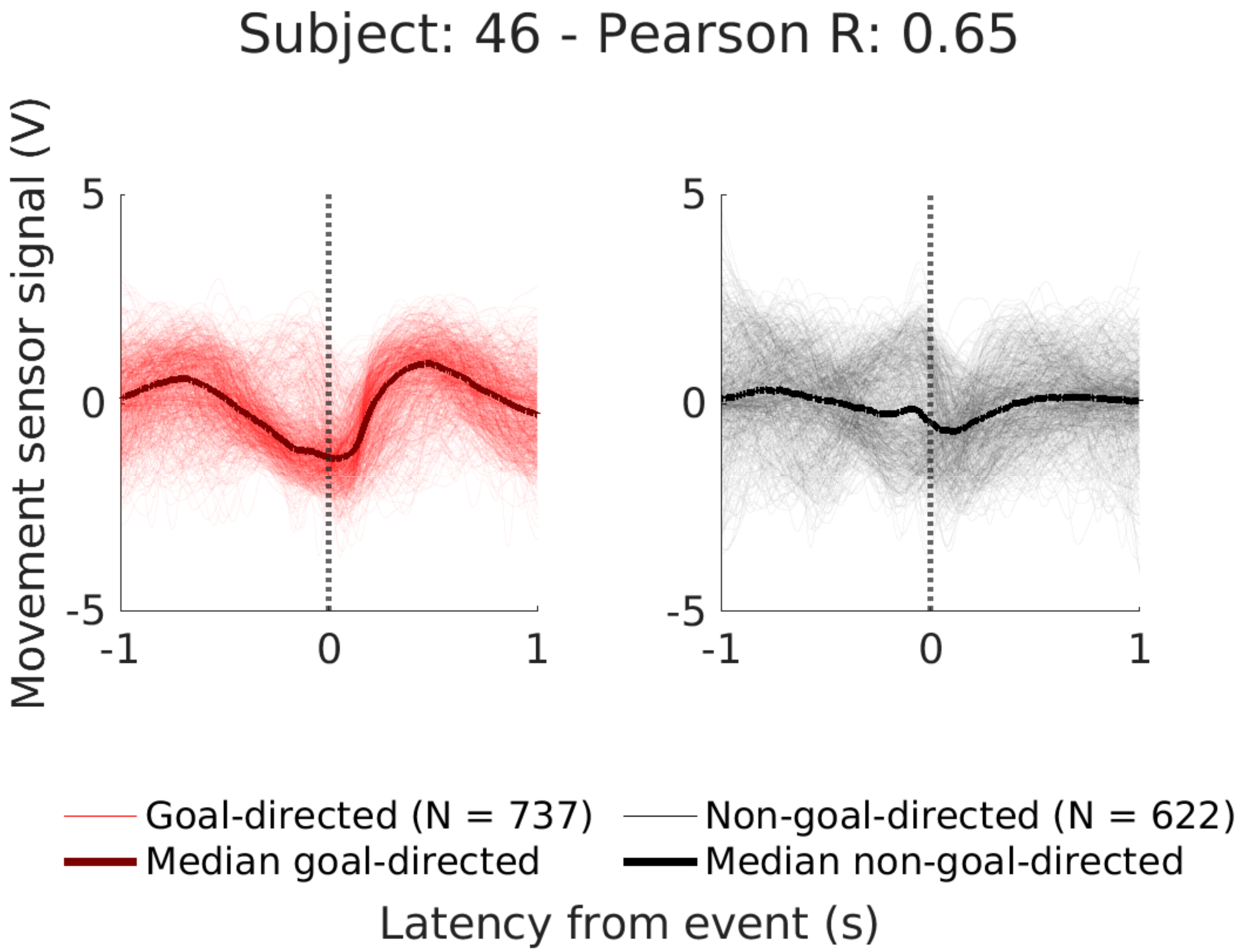

Subject: 46 - Pearson R: 0.65
Movement sensor signal (V)
5
0
-5
-1
0
1
5
0
-5
-1
0
1
Goal-directed (N = 737)
Median goal-directed
Non-goal-directed (N = 622)
Median non-goal-directed
Latency from event (s)

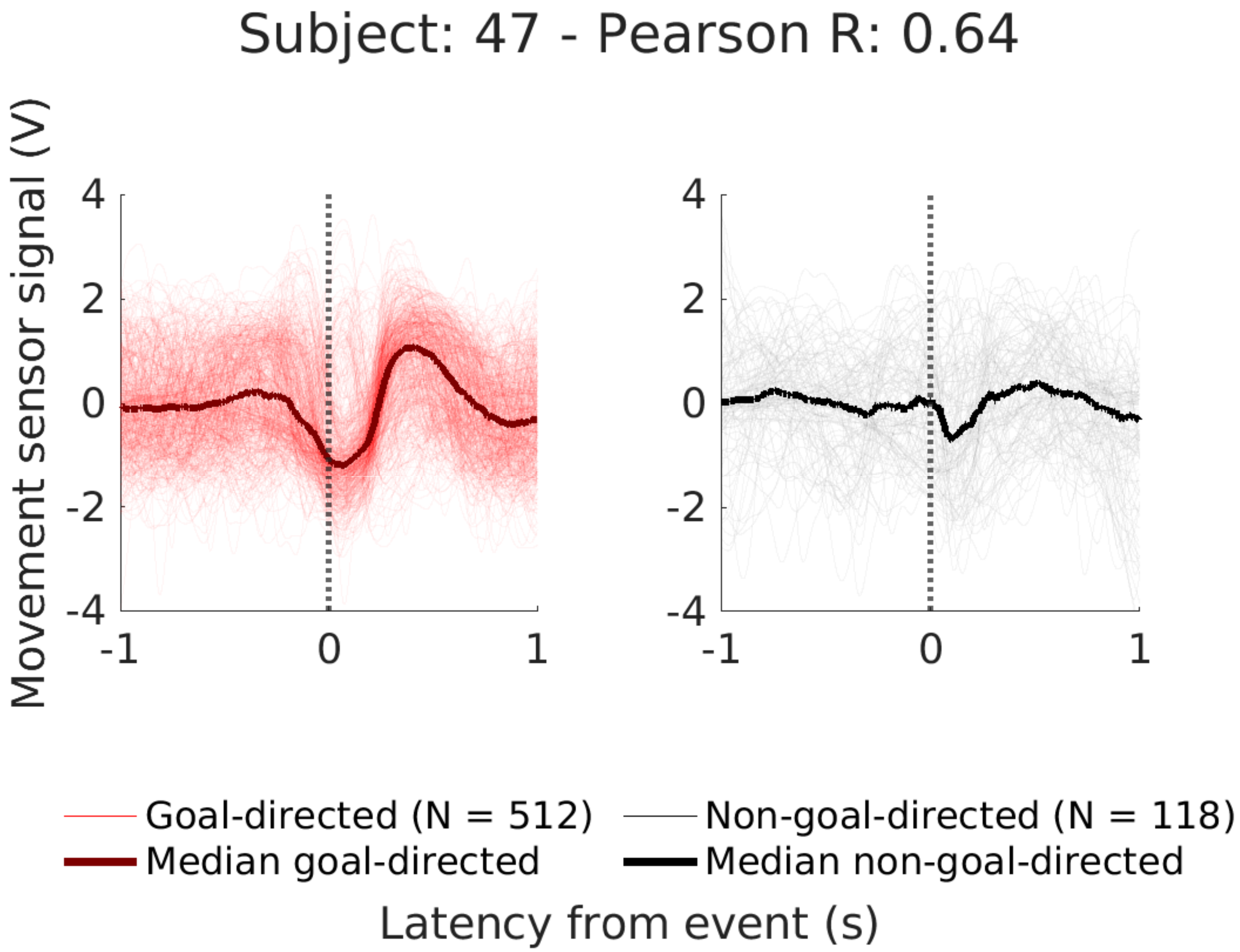

Subject: 47 - Pearson R: 0.64
Movement sensor signal (V)
4
2
0
-2
-4
-1
0
1
Goal-directed (N = 512)
Median goal-directed
Non-goal-directed (N = 118)
Median non-goal-directed
Latency from event (s)

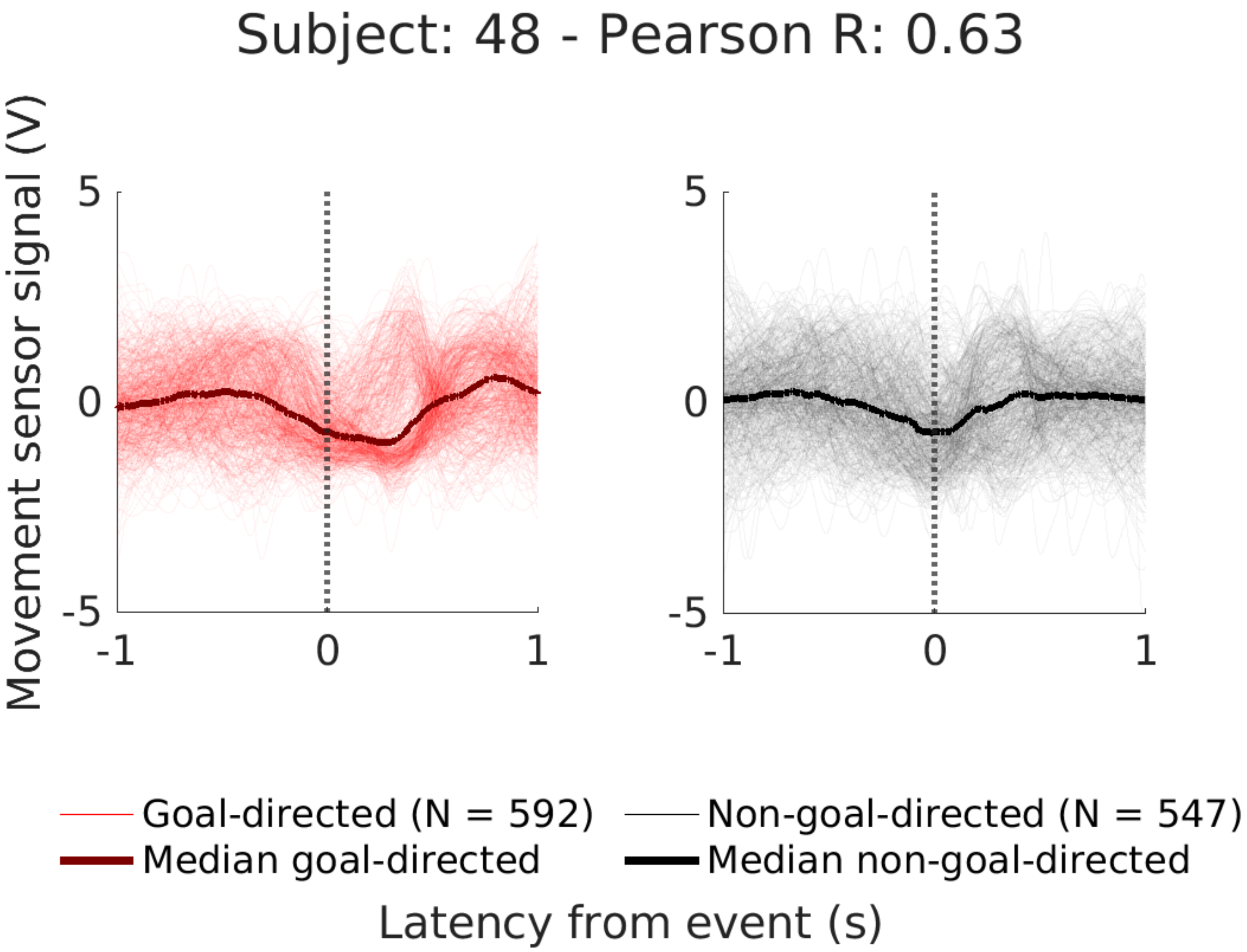
Subject: 48 - Pearson R: 0.63
Movement sensor signal (V)
5
0
-5
-1
0
1
5
0
-5
-1
0
1
Goal-directed (N = 592)
Non-goal-directed (N = 547)
Median goal-directed
Median non-goal-directed
Latency from event (s)

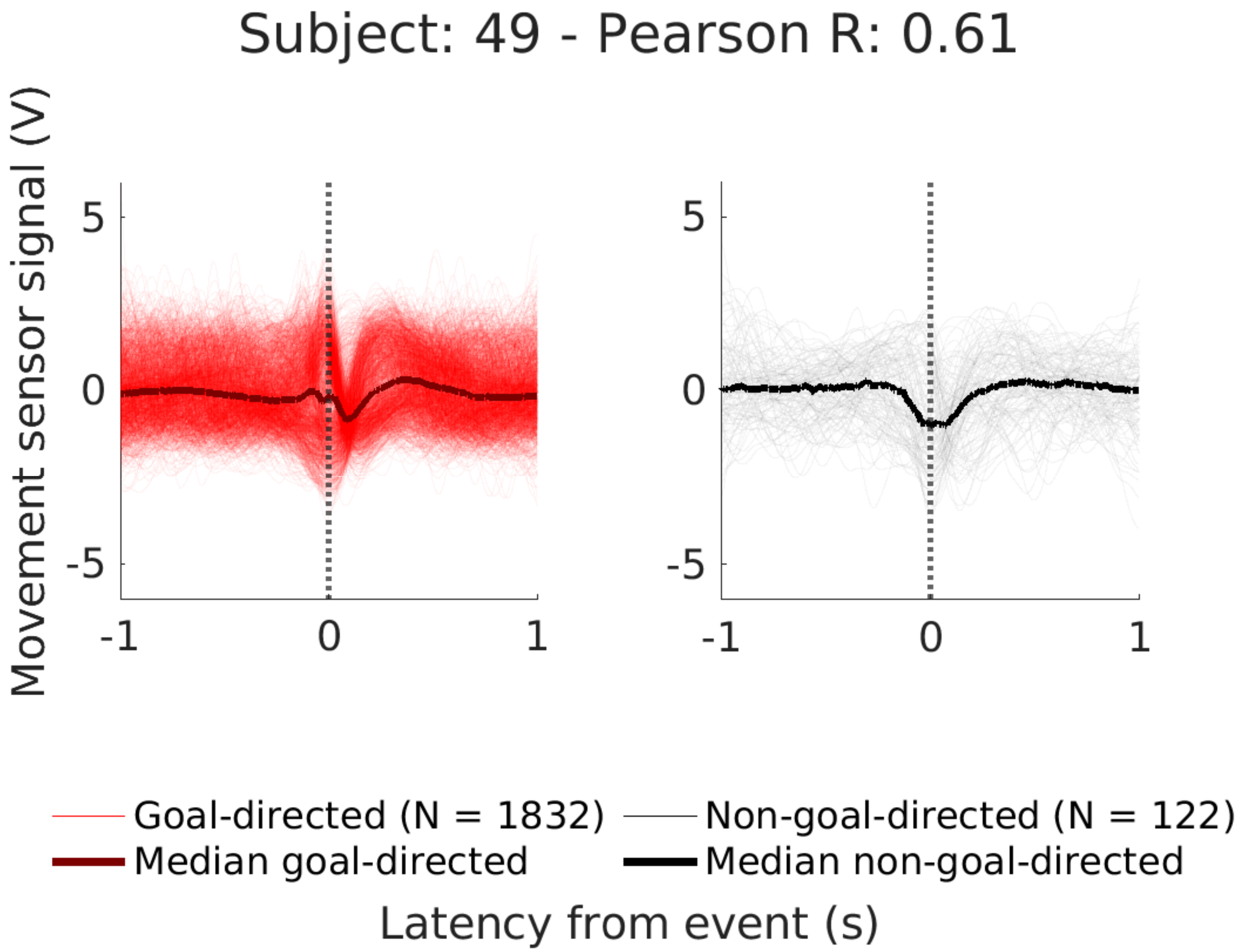
Subject: 49 - Pearson R: 0.61
Movement sensor signal (V)
5
0
-5
-1
0
1
5
0
-5
-1
0
1
Goal-directed (N = 1832)
Median goal-directed
Non-goal-directed (N = 122)
Median non-goal-directed
Latency from event (s)

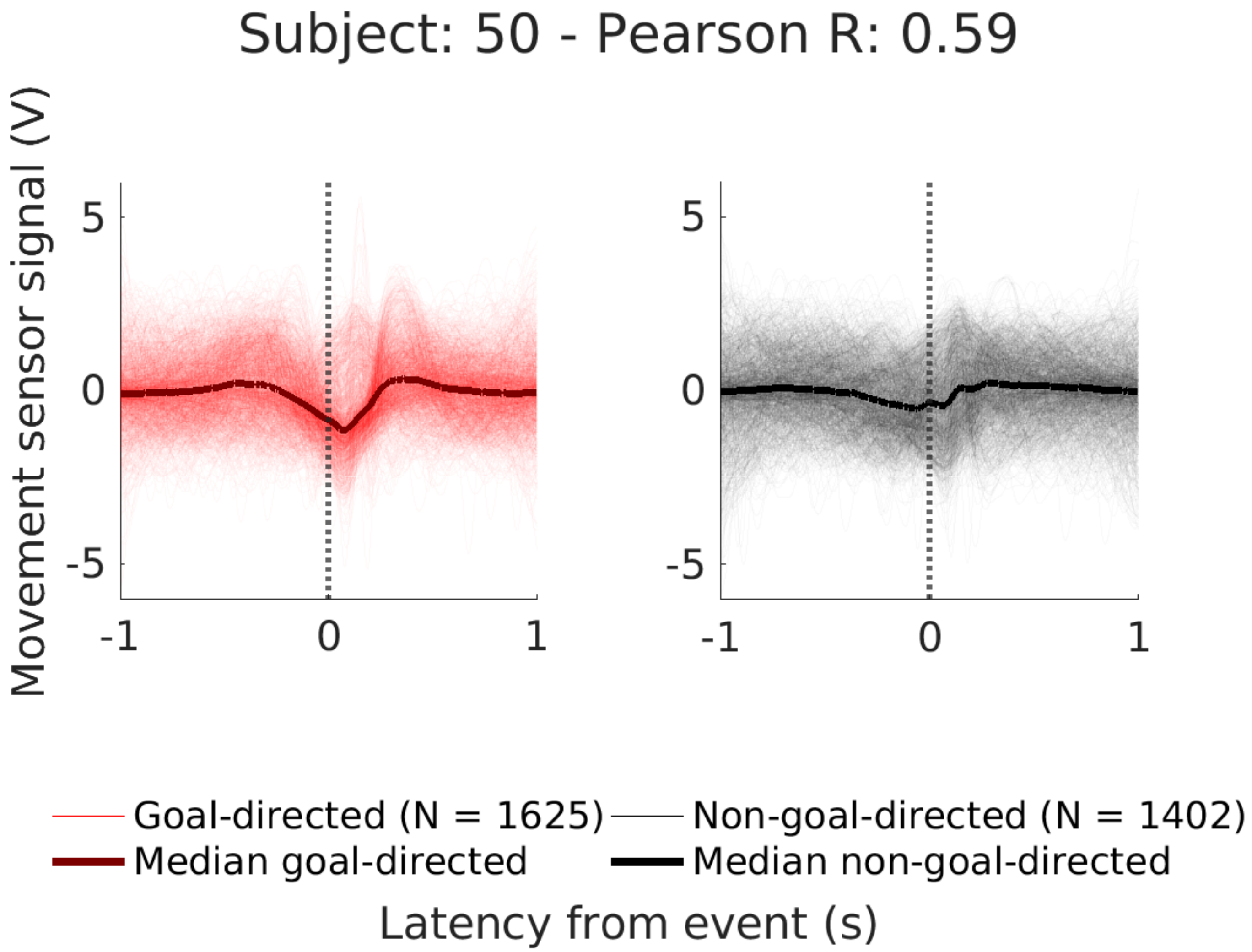

Subject: 50 - Pearson R: 0.59
Movement sensor signal (V)
5
0
-5
-1
0
1
5
0
-5
-1
0
1
Goal-directed (N = 1625)
Non-goal-directed (N = 1402)
Median goal-directed
Median non-goal-directed
Latency from event (s)

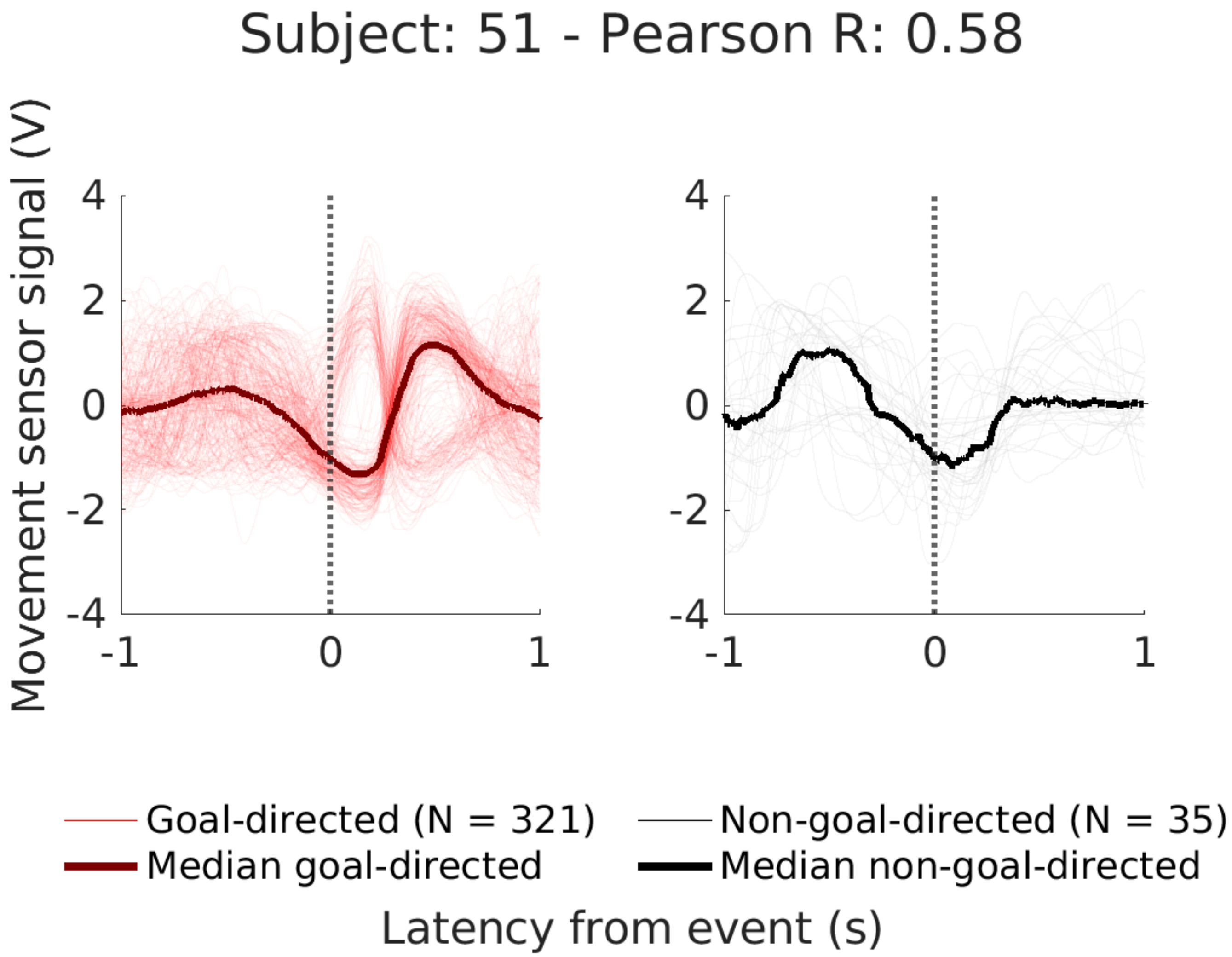

Subject: 51 - Pearson R: 0.58
Movement sensor signal (V)
4
2
0
-2
-4
-1
0
1
Goal-directed (N = 321)
Median goal-directed
Non-goal-directed (N = 35)
Median non-goal-directed
Latency from event (s)

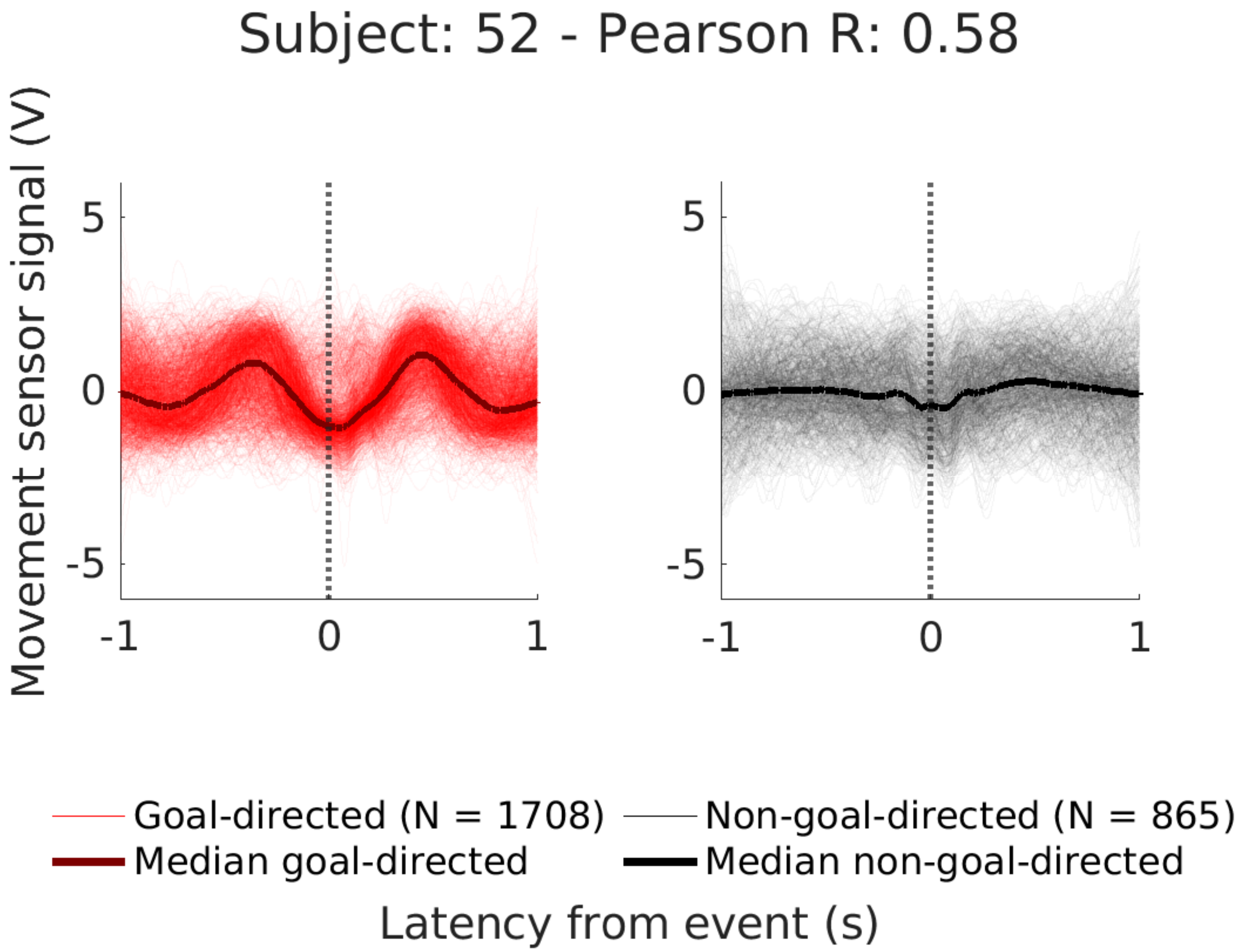

Subject: 52 - Pearson R: 0.58
Movement sensor signal (V)
5
0
-5
-1
0
1
5
0
-5
-1
0
1
Goal-directed (N = 1708)
Median goal-directed
Non-goal-directed (N = 865)
Median non-goal-directed
Latency from event (s)

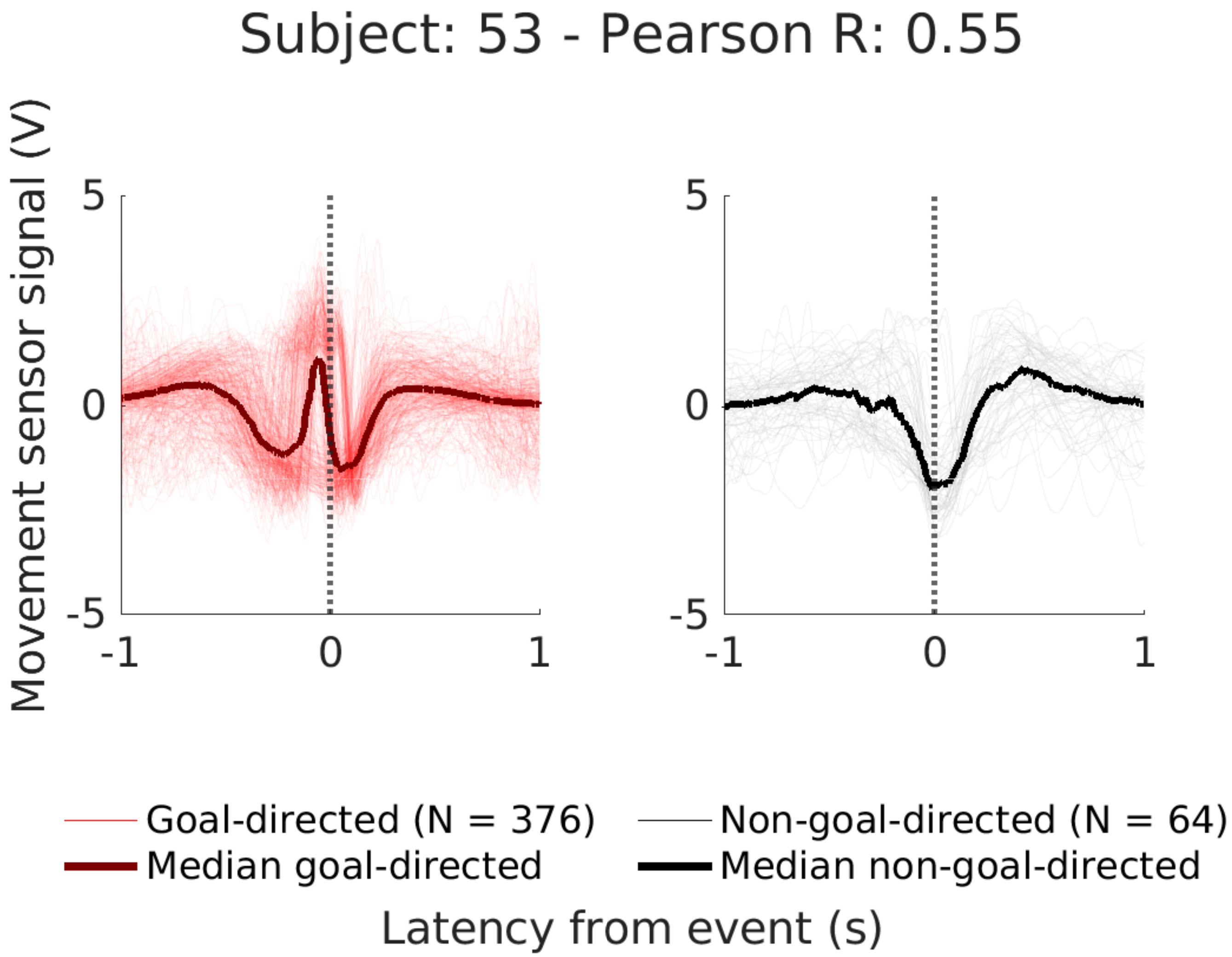

Subject: 53 - Pearson R: 0.55
Movement sensor signal (V)
5
0
-5
-1
0
1
5
0
-5
-1
0
1
Goal-directed (N = 376)
Median goal-directed
Non-goal-directed (N = 64)
Median non-goal-directed
Latency from event (s)

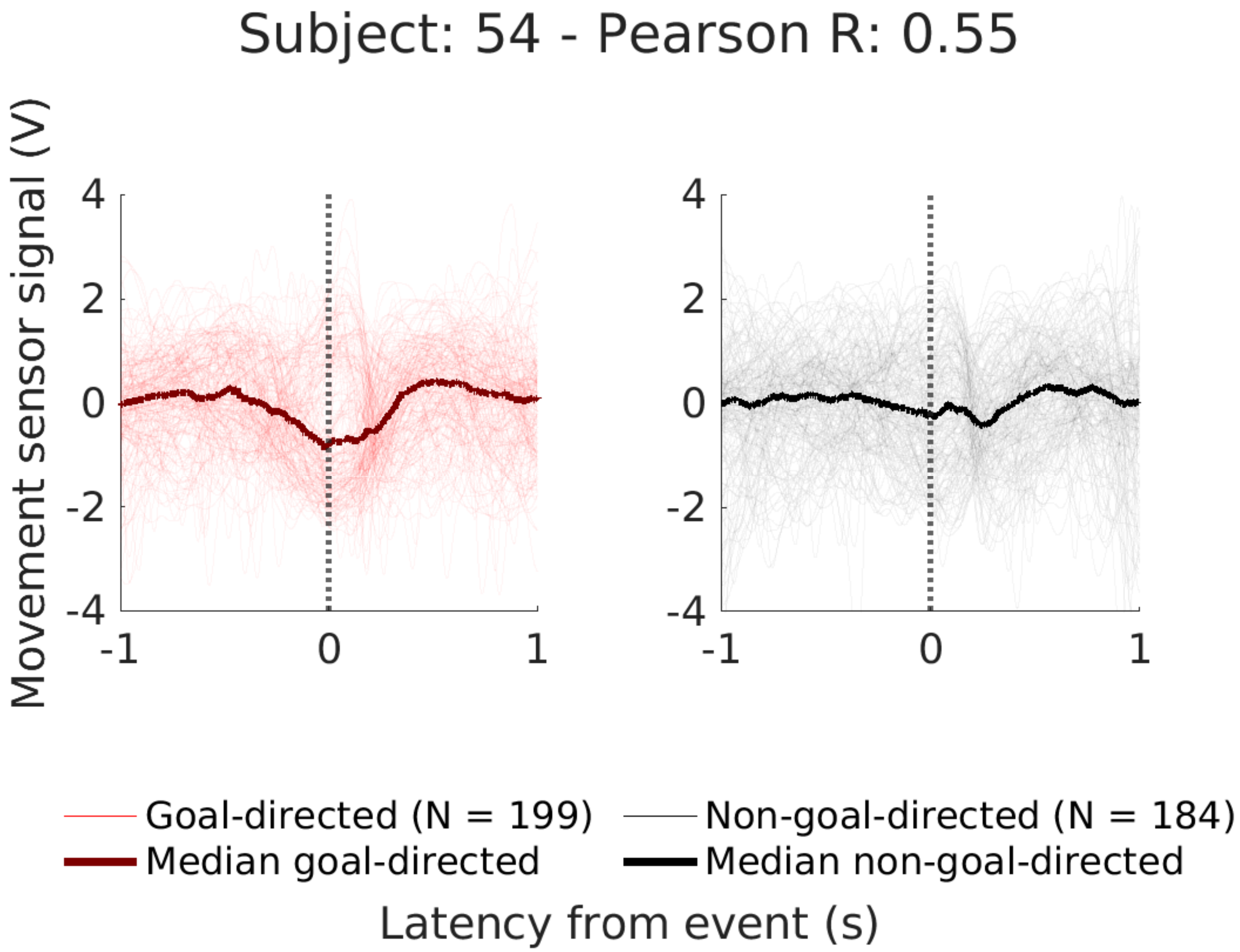

Subject: 54 - Pearson R: 0.55
Movement sensor signal (V)
4
2
0
-2
-4
-1
0
1
Goal-directed (N = 199)
Median goal-directed
Non-goal-directed (N = 184)
Median non-goal-directed
Latency from event (s)

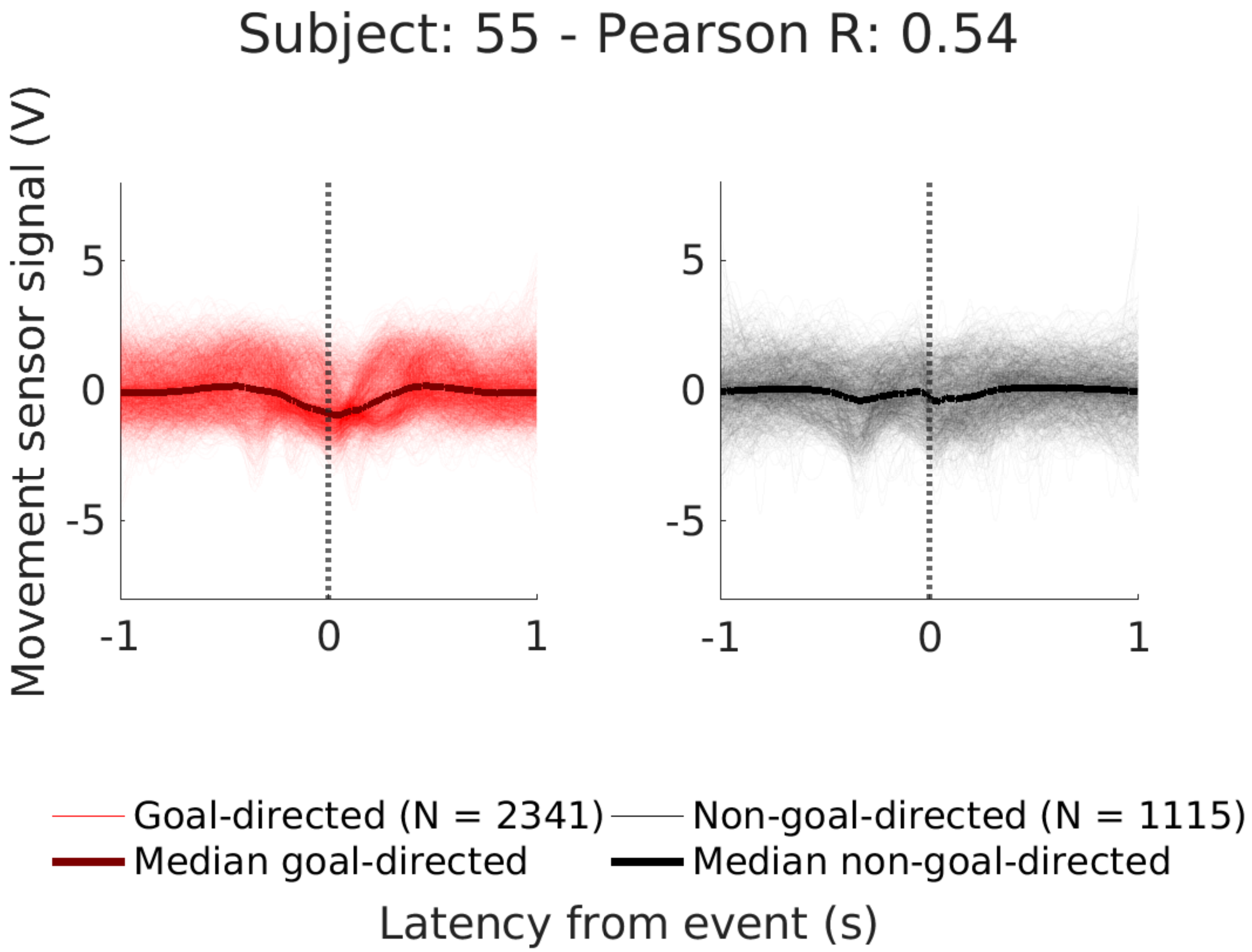

Subject: 55 - Pearson R: 0.54
Movement sensor signal (V)
5
0
-5
-1
0
1
Goal-directed (N = 2341)
Non-goal-directed (N = 1115)
Median goal-directed
Median non-goal-directed
Latency from event (s)

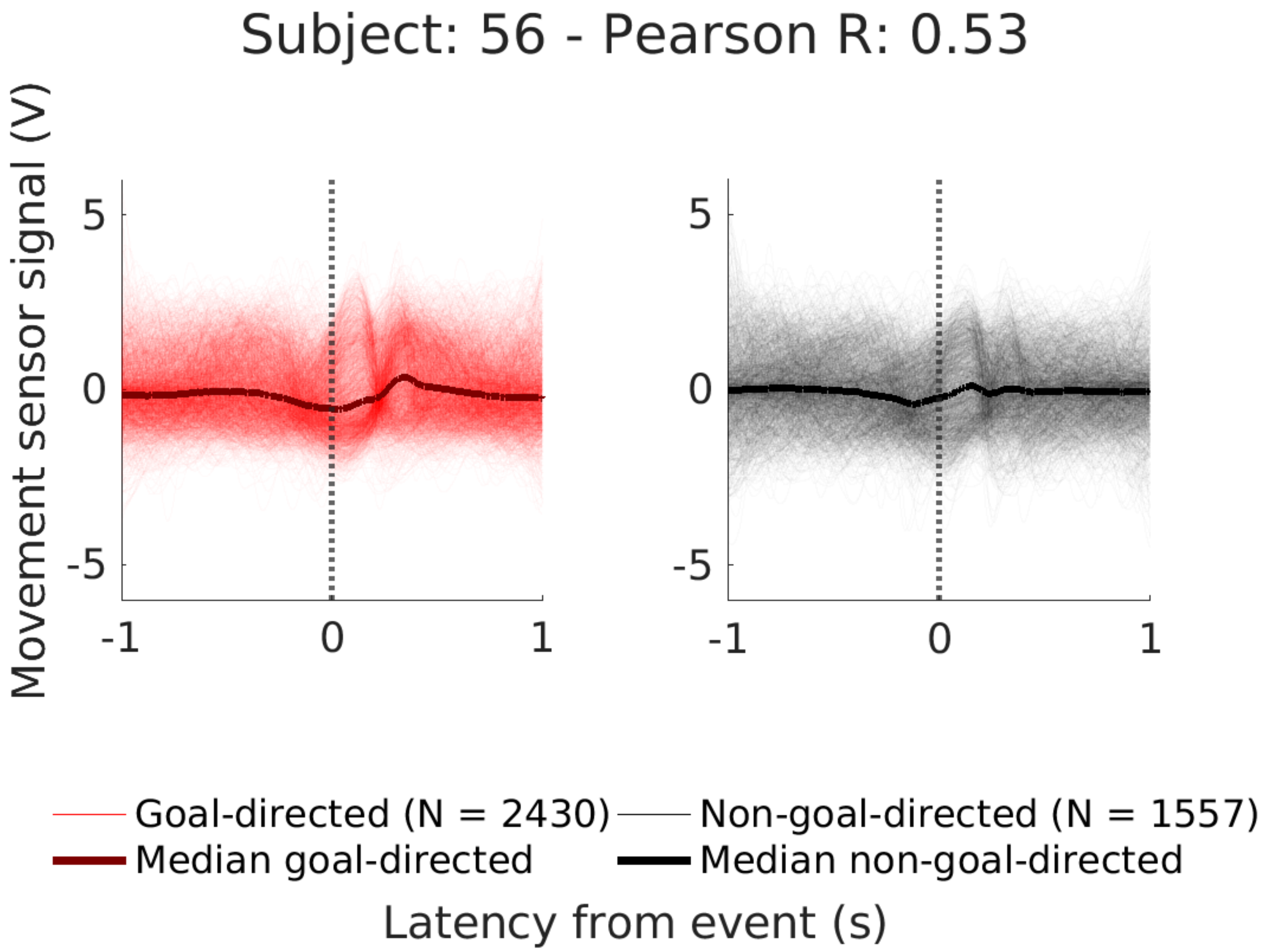

Subject: 56 - Pearson R: 0.53
Movement sensor signal (V)
5
0
-5
-1
0
1
Goal-directed (N = 2430)
Non-goal-directed (N = 1557)
Median goal-directed
Median non-goal-directed
Latency from event (s)

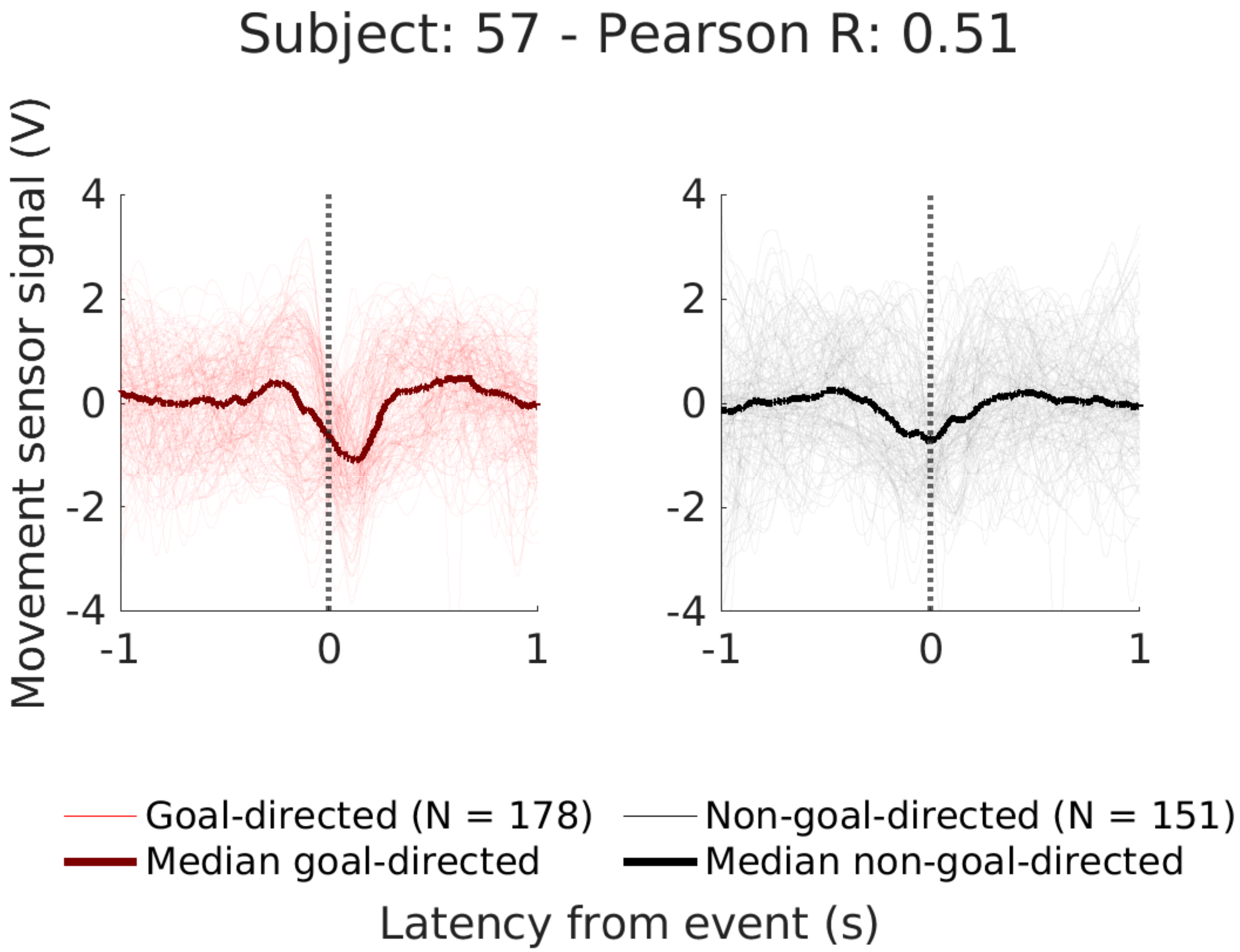
Subject: 57 - Pearson R: 0.51
Movement sensor signal (V)
4
2
0
-2
-4
-1
0
1
4
2
0
-2
-4
-1
0
1
Goal-directed (N = 178)
Median goal-directed
Non-goal-directed (N = 151)
Median non-goal-directed
Latency from event (s)

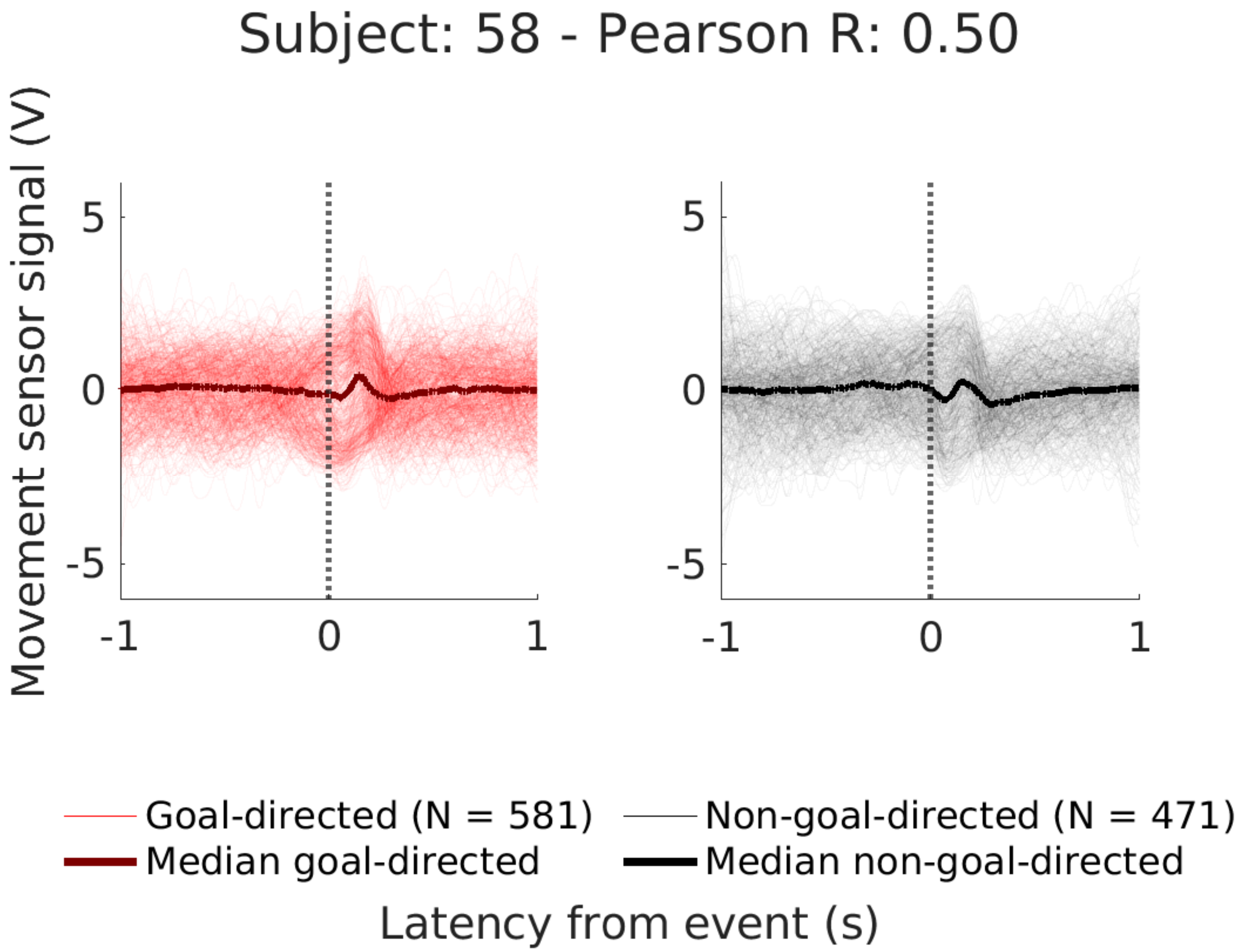

Subject: 58 - Pearson R: 0.50
Movement sensor signal (V)
5
0
-5
-1
0
1
5
0
-5
-1
0
1
Goal-directed (N = 581)
Median goal-directed
Non-goal-directed (N = 471)
Median non-goal-directed
Latency from event (s)

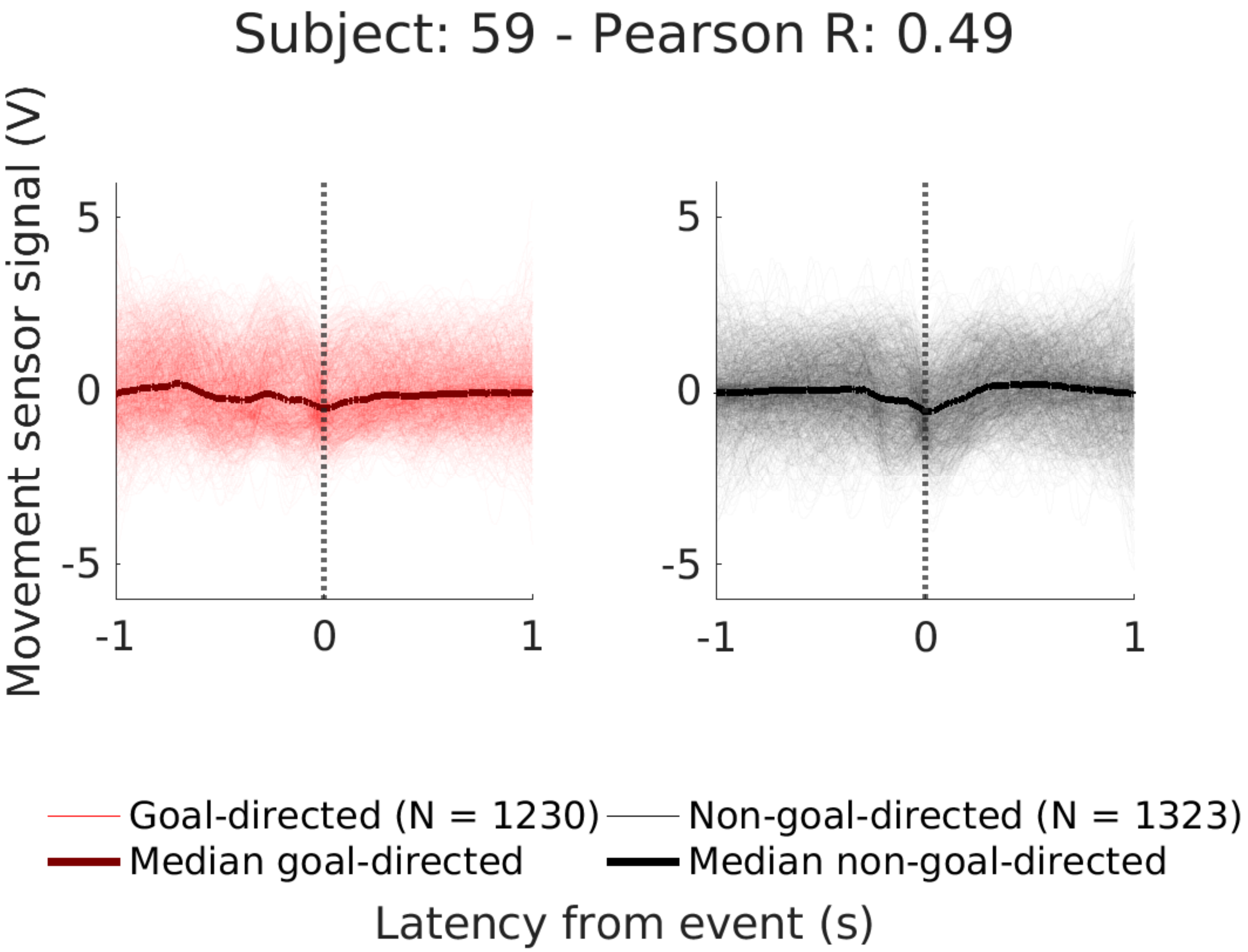

Subject: 59 - Pearson R: 0.49
Movement sensor signal (V)
5
0
-5
-1
0
1
5
0
-5
-1
0
1
Goal-directed (N = 1230)
Non-goal-directed (N = 1323)
Median goal-directed
Median non-goal-directed
Latency from event (s)

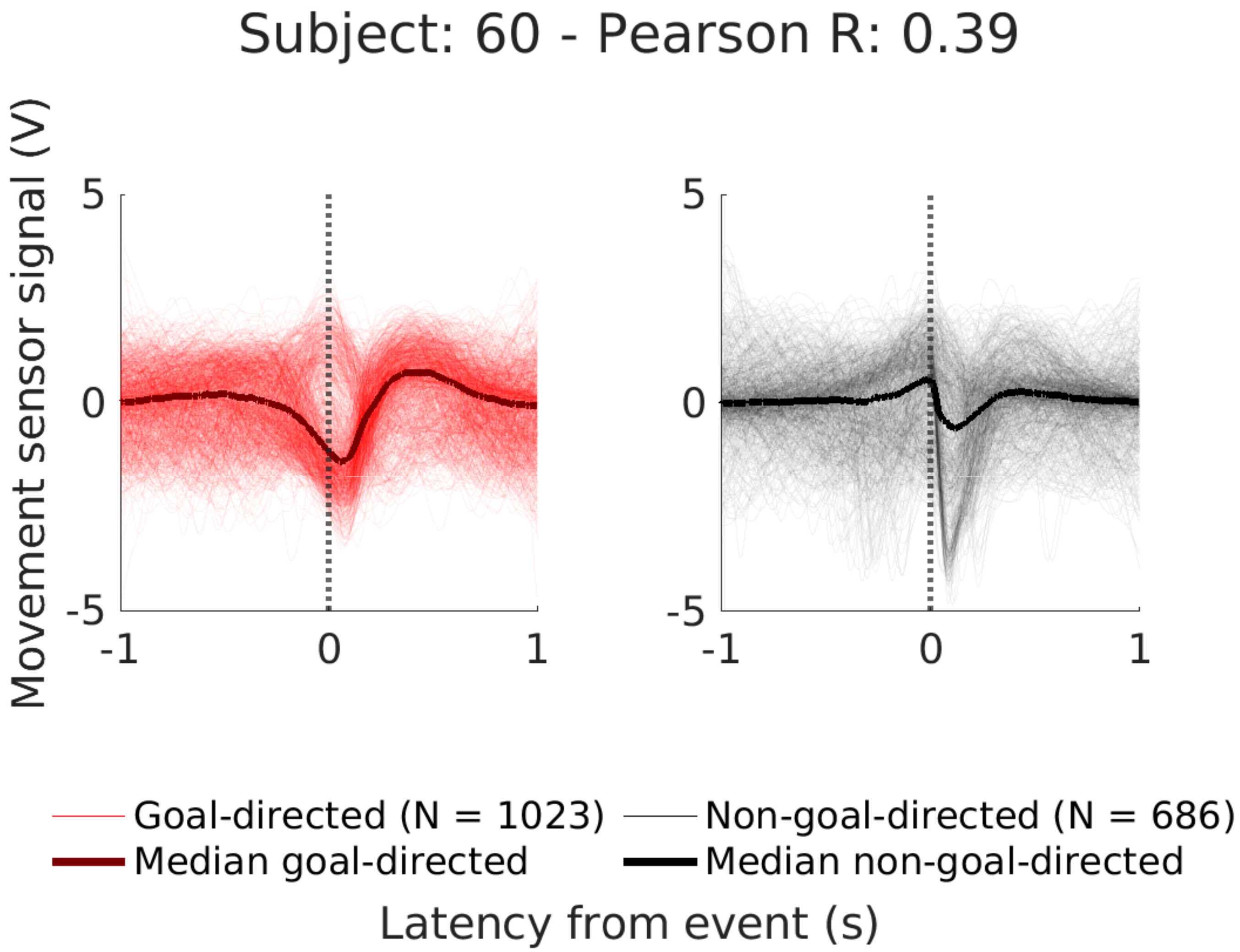
Subject: 60 - Pearson R: 0.39
Movement sensor signal (V)
5
0
-5
-1
0
1
5
0
-5
-1
0
1
Goal-directed (N = 1023)
Non-goal-directed (N = 686)
Median goal-directed
Median non-goal-directed
Latency from event (s)

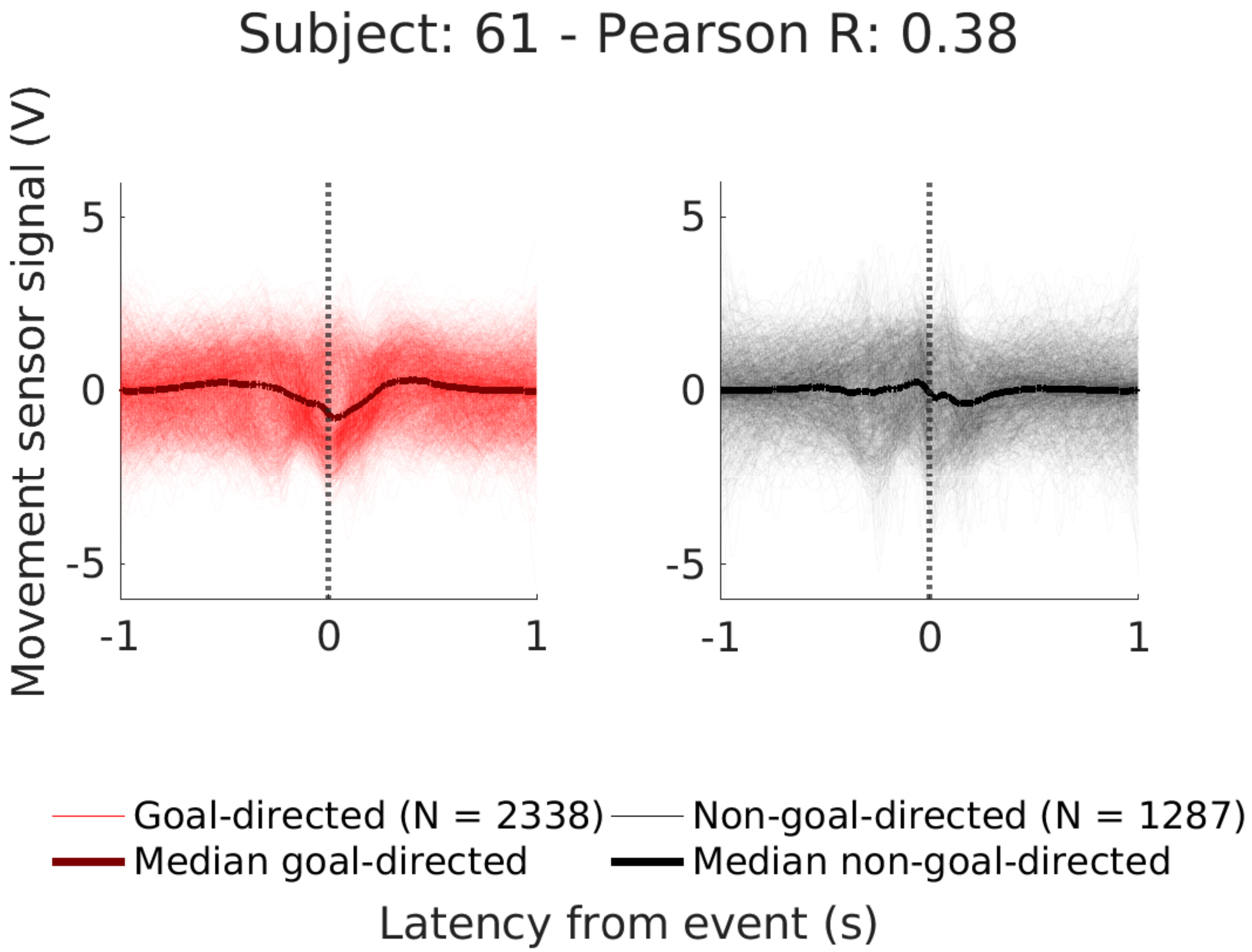
Subject: 61 - Pearson R: 0.38
Movement sensor signal (V)
5
0
-5
-1
0
1
Goal-directed (N = 2338)
Non-goal-directed (N = 1287)
Median goal-directed
Median non-goal-directed
Latency from event (s)

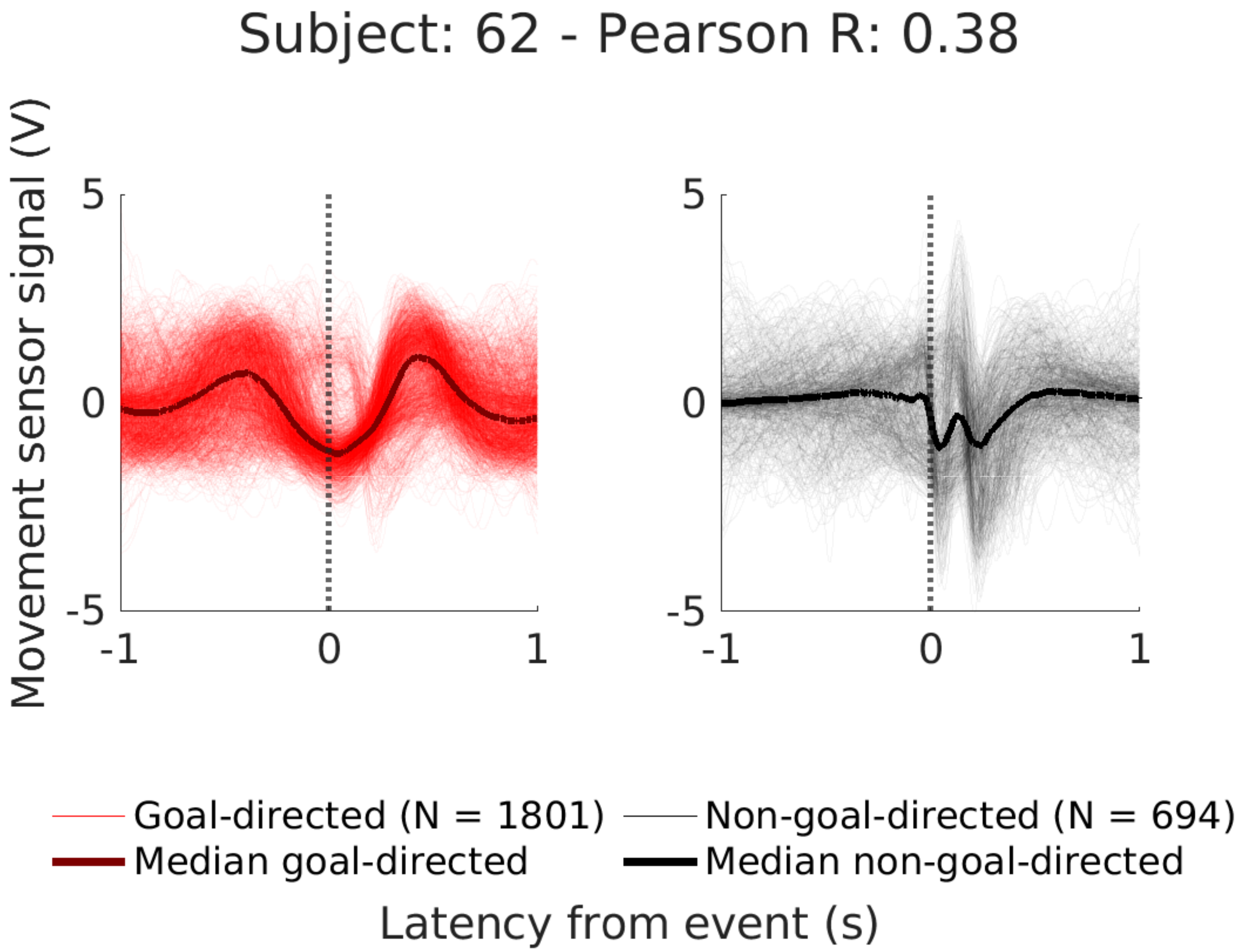

Subject: 62 - Pearson R: 0.38
Movement sensor signal (V)
5
0
-5
-1
0
1
5
0
-5
-1
0
1
Goal-directed (N = 1801)
Median goal-directed
Non-goal-directed (N = 694)
Median non-goal-directed
Latency from event (s)

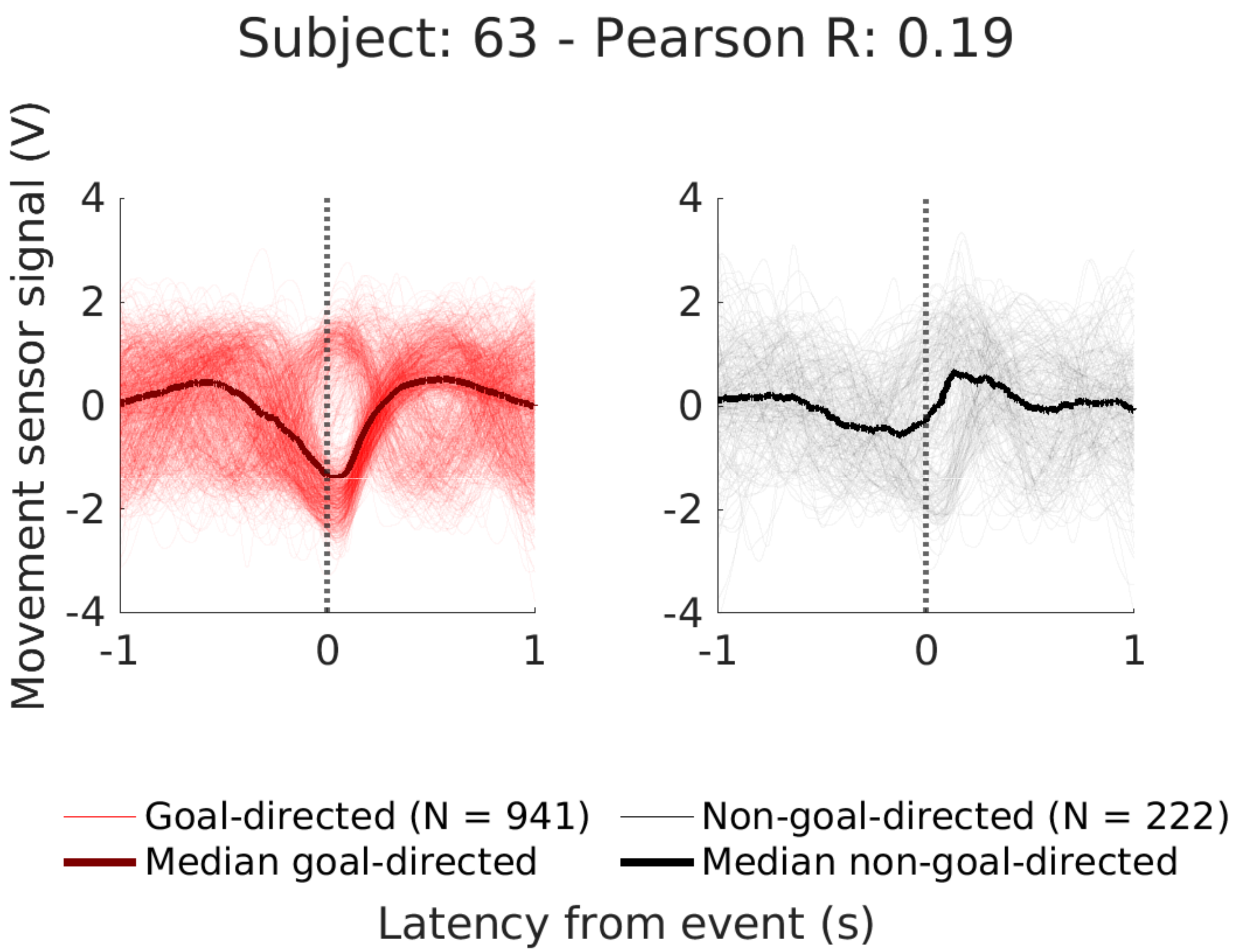

Subject: 63 - Pearson R: 0.19
Movement sensor signal (V)
4
2
0
-2
-4
-1
0
1
Goal-directed (N = 941)
Non-goal-directed (N = 222)
Median goal-directed
Median non-goal-directed
Latency from event (s)

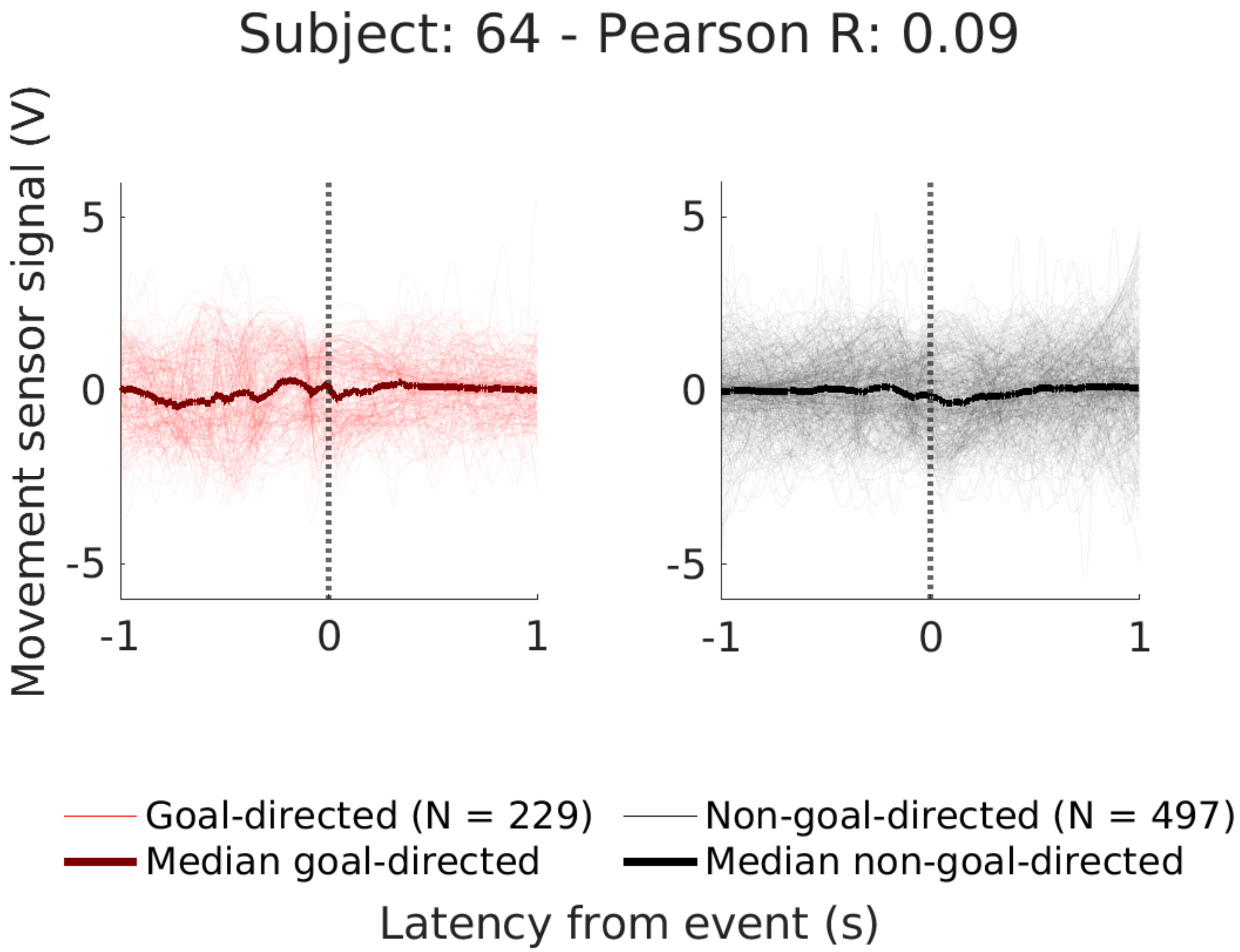
Subject: 64 - Pearson R: 0.09
Movement sensor signal (V)
5
0
-5
-1
0
1
5
0
-5
-1
0
1
Goal-directed (N = 229)
Median goal-directed
Non-goal-directed (N = 497)
Median non-goal-directed
Latency from event (s)

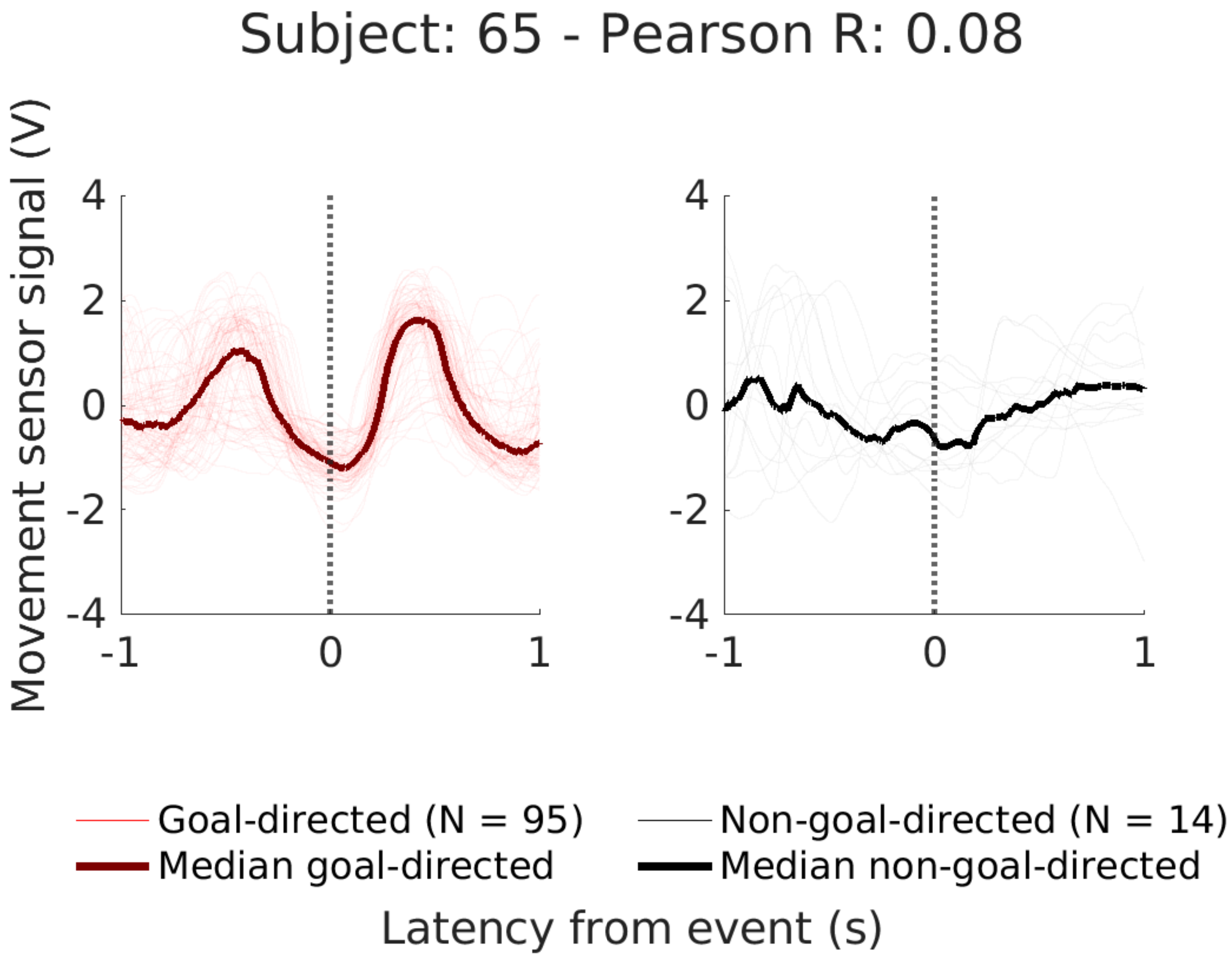

Subject: 65 - Pearson R: 0.08
Movement sensor signal (V)
4
2
0
-2
-4
-1
0
1
Goal-directed (N = 95)
Median goal-directed
Non-goal-directed (N = 14)
Median non-goal-directed
Latency from event (s)

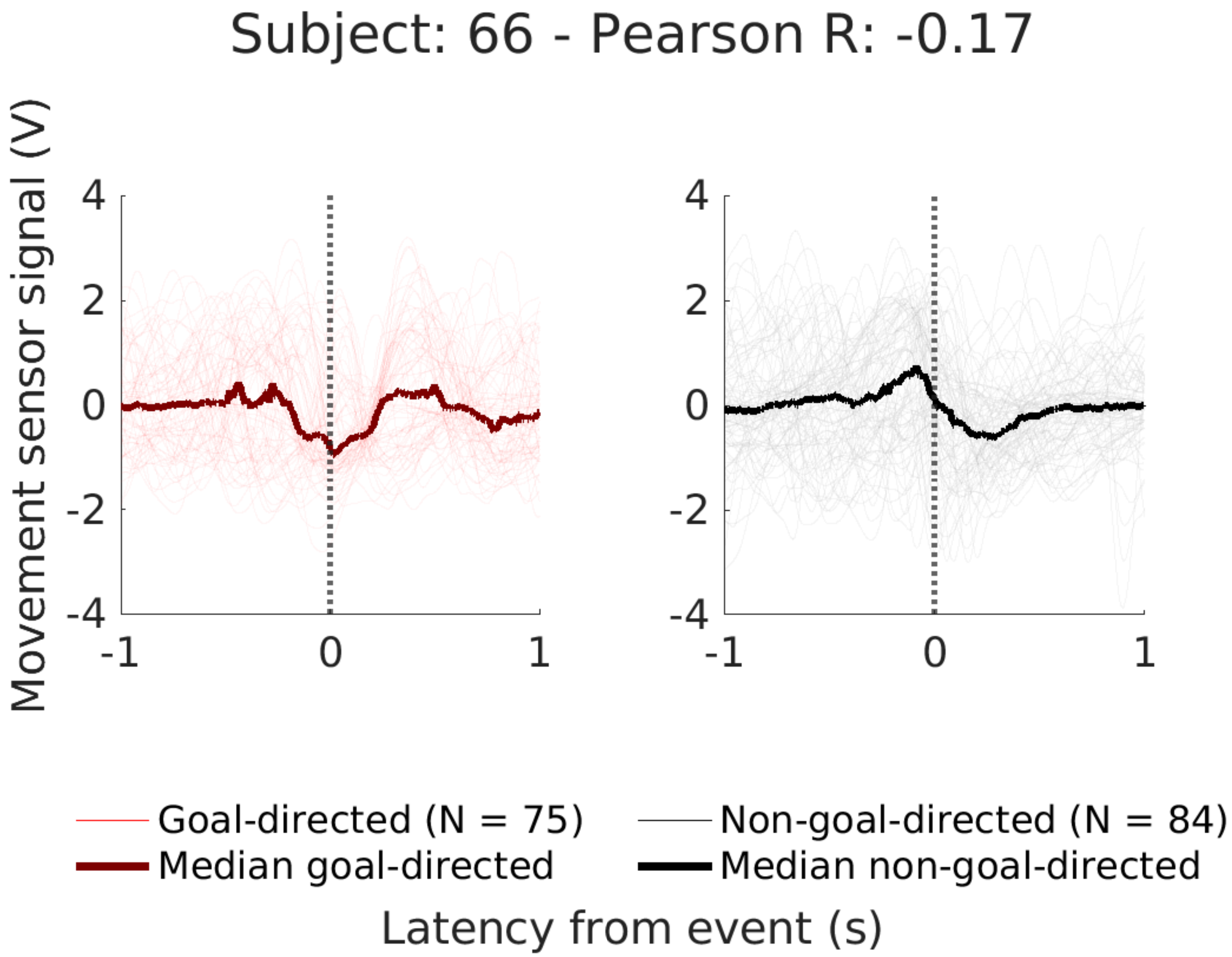
Subject: 66 - Pearson R: -0.17
Movement sensor signal (V)
4
2
0
-2
-4
-1
0
1
4
2
0
-2
-4
-1
0
1
Goal-directed (N = 75)
Median goal-directed
Non-goal-directed (N = 84)
Median non-goal-directed
Latency from event (s)

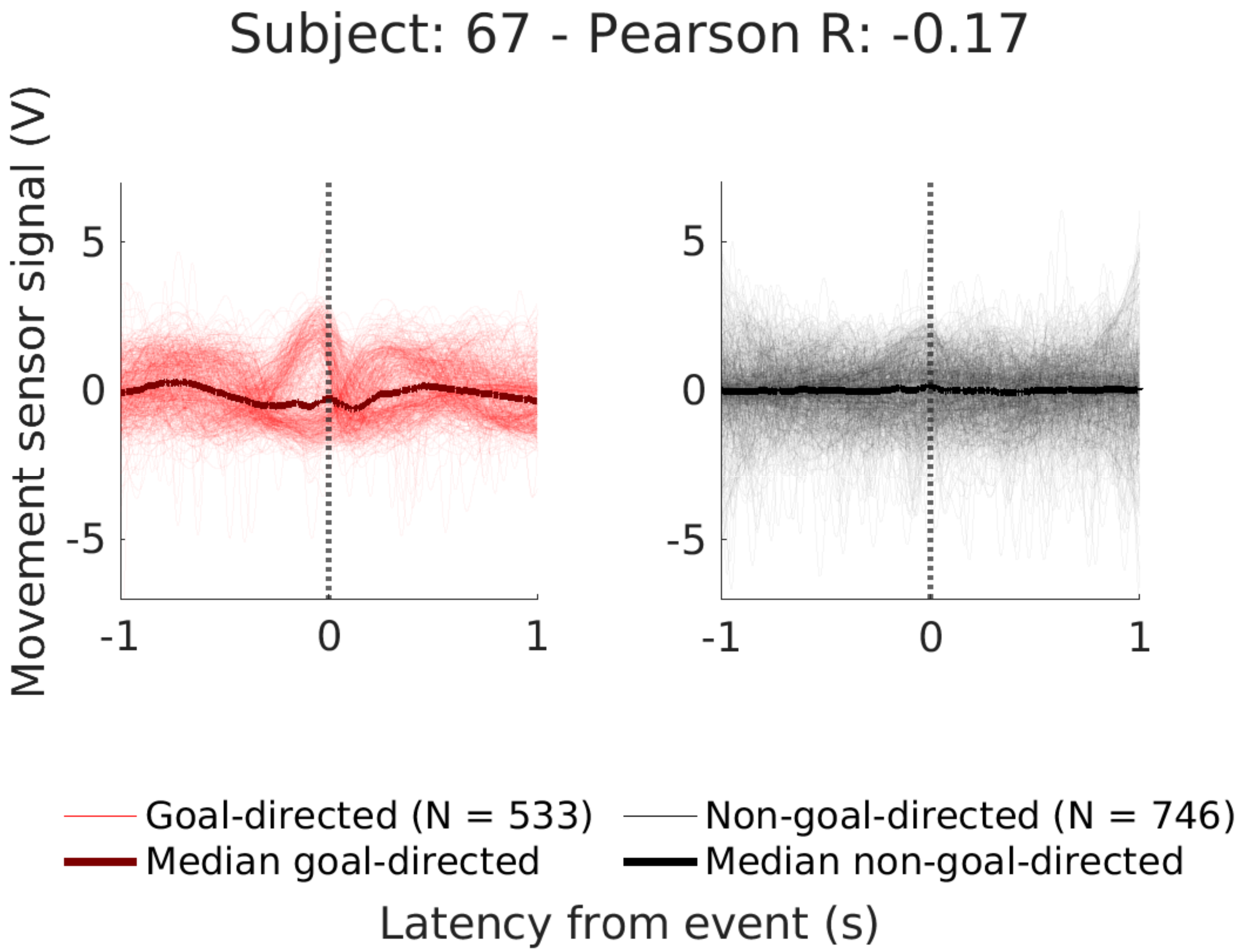

Subject: 67 - Pearson R: -0.17
Movement sensor signal (V)
5
0
-5
-1
0
1
5
0
-5
-1
0
1
Goal-directed (N = 533)
Non-goal-directed (N = 746)
Median goal-directed
Median non-goal-directed
Latency from event (s)

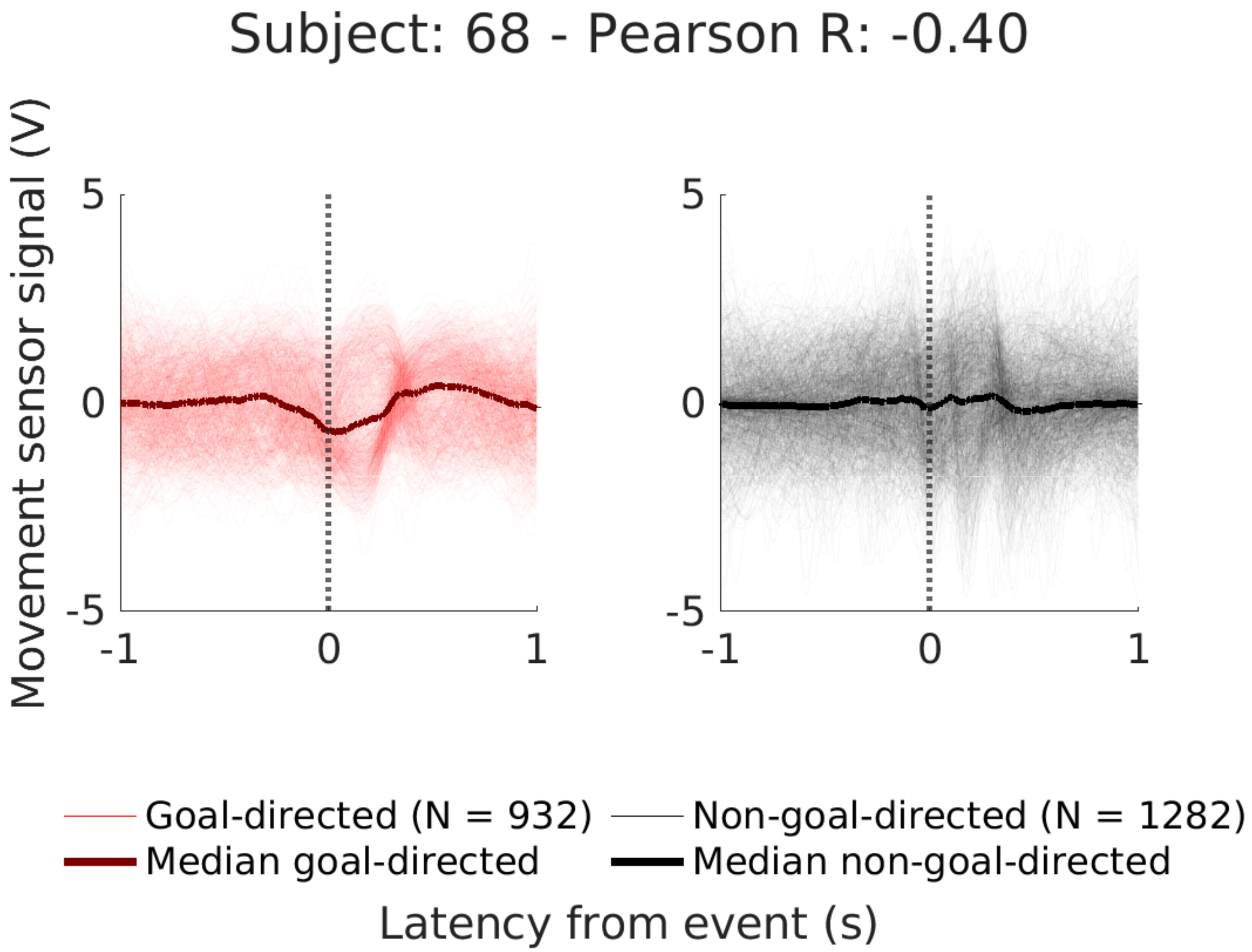
Subject: 68 - Pearson R: -0.40
Movement sensor signal (V)
5
0
-5
-1
0
1
5
0
-5
-1
0
1
Goal-directed (N = 932)
Non-goal-directed (N = 1282)
Median goal-directed
Median non-goal-directed
Latency from event (s)
