## Supplementary Methods for "Neural processing of goal and non-goal-directed movements on the smartphone"

**Supplementary Methods Figure 1.** Example movement sensor signals.............................................. 1

**Supplementary Methods Figure 2.** Participant selection diagram ...................................................... 2

**Supplementary Methods Figure 3.** Alignment model methods and results ....................................... 3

**Supplementary Methods Table 1.** Alignment model hyperparameters.............................................. 4

**Supplementary Methods Figure 4.** Alignment model correction........................................................ 5

**Supplementary Methods Table 2.** Goal and non-goal-directed movement identification model hyperparameters ......................................................................................................................... 6

**Supplementary Methods Figure 5.** Participant selection based on kinematic profile ......................... 7

**Supplementary Methods Figure 6.** Goal and non-goal-directed movements during smartphone apps ......................................................................................................................................... 8

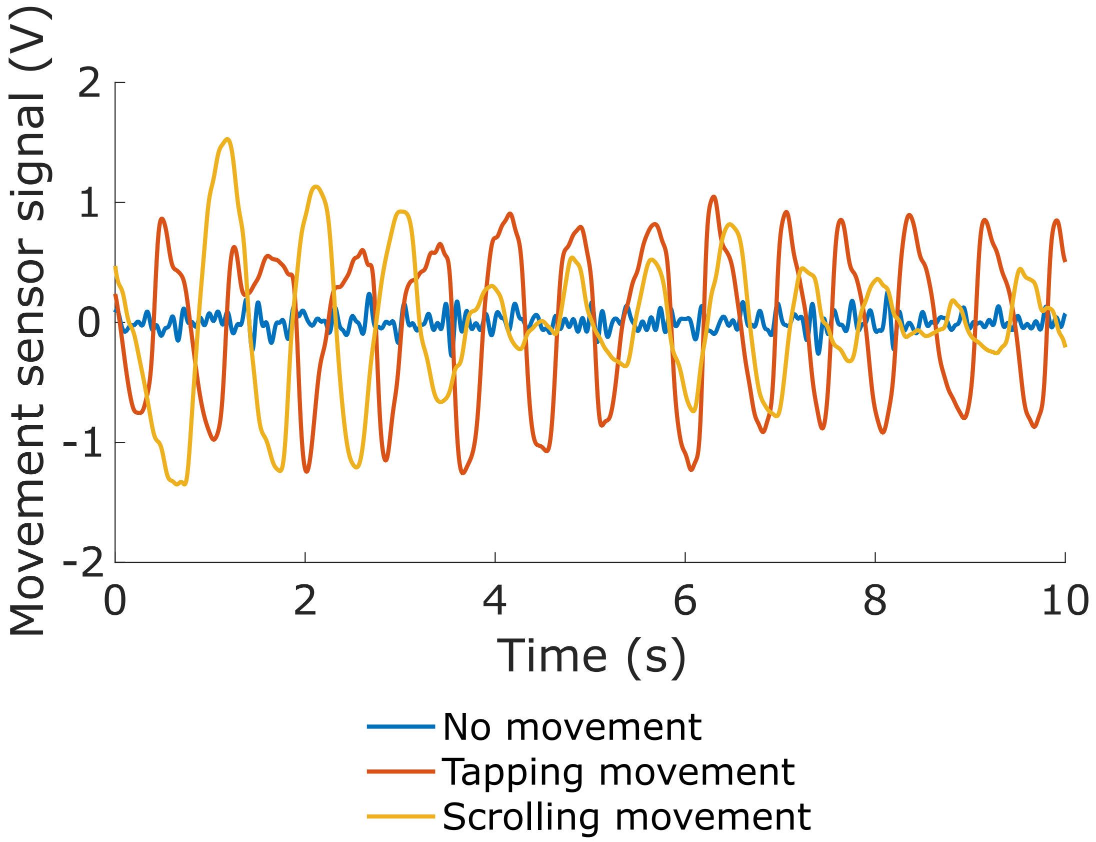


**Supplementary Methods Figure 1.** Common movements generated during smartphone use compared to no movements as measurement by the movement sensor.

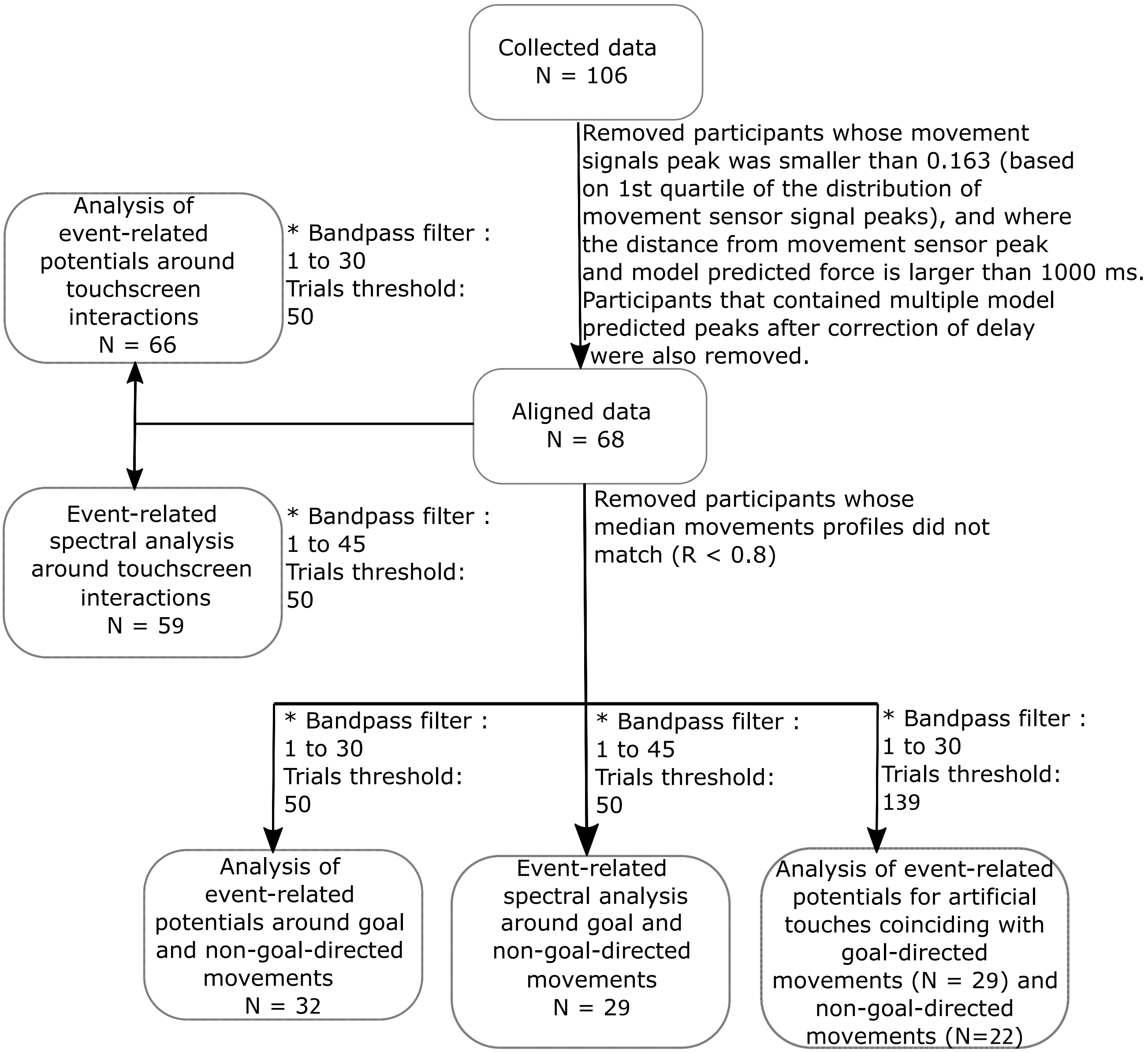


**Supplementary Methods Figure 2.** Participant selection at multiple stages of analysis.

* Participants were removed through the combination of a bandpass filter, artifact rejection (+/- 80 µV) and not enough trials for each event remaining.

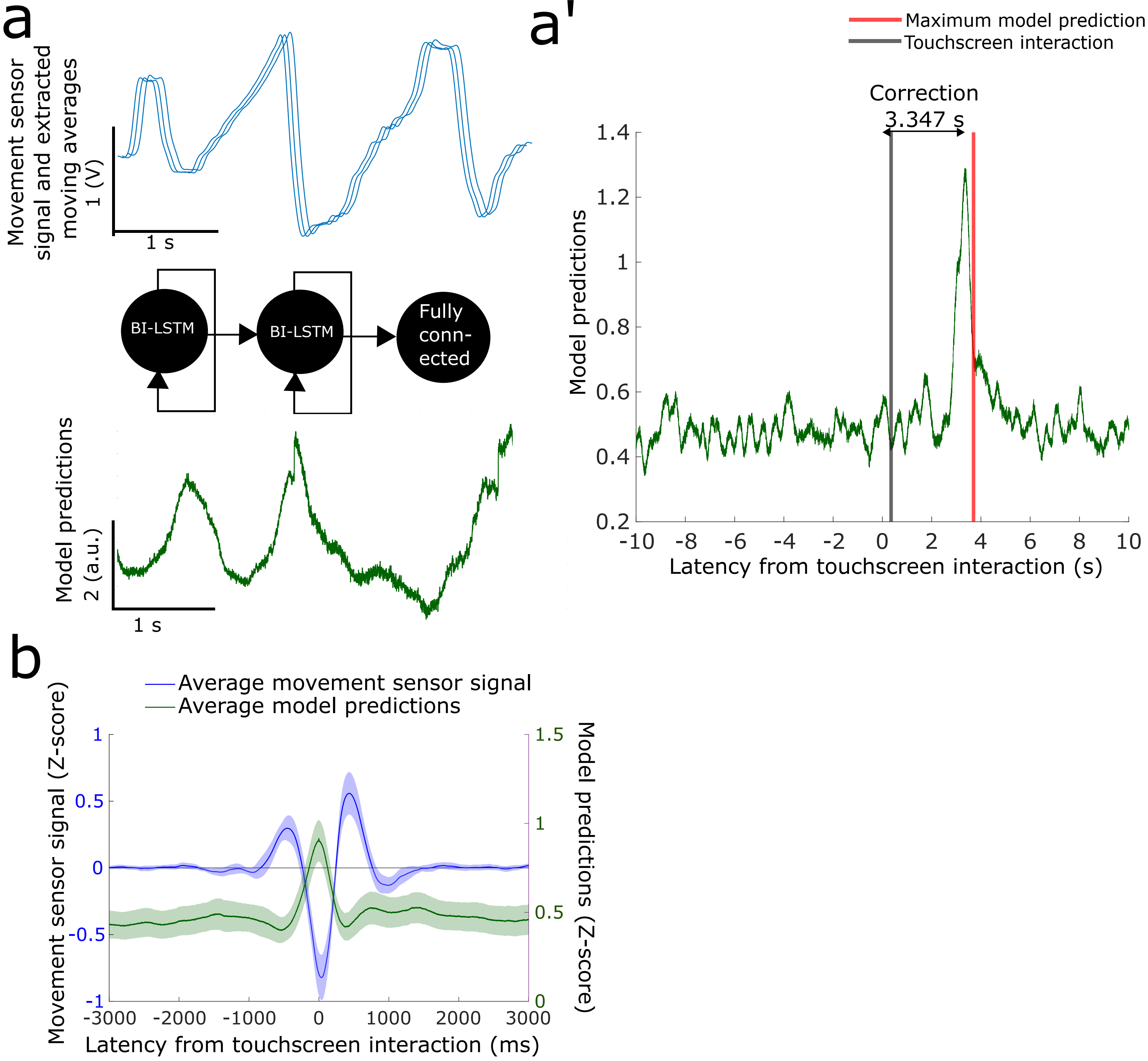


**Supplementary Methods Figure 3.** Alignment of smartphone data to EEG and movement sensor signals. **(a)** A BI-LSTM model was trained with movement sensor signals and 100 extracted moving averages to predict force sensor values. **(a')** The alignment was performed by correcting for the delay between the average predicted signal and the recorded touchscreen interaction. **(b)** Grand average of the movement sensor signals after alignment indicate that the lowest point of the bend of the thumb is near the touchscreen interaction (Z-score normalized for visualization). Grand average model predictions after alignment show that the model identified the location of the touchscreen interactions (Z-score normalized for visualization). Overall indicating successful alignment of the data.

**Supplementary Methods Table 1**

*Hyperparameters for alignment model*

| Parameter | Value |
|---|---|
| Train/Validation/Test split | 80/10/10 |
| Total number of trainable parameters | 425.480 |
| Optimizer | adam |
| Learning rate | 0.001 |
| Loss | Mean squared error |
| Max epoch | 5 |

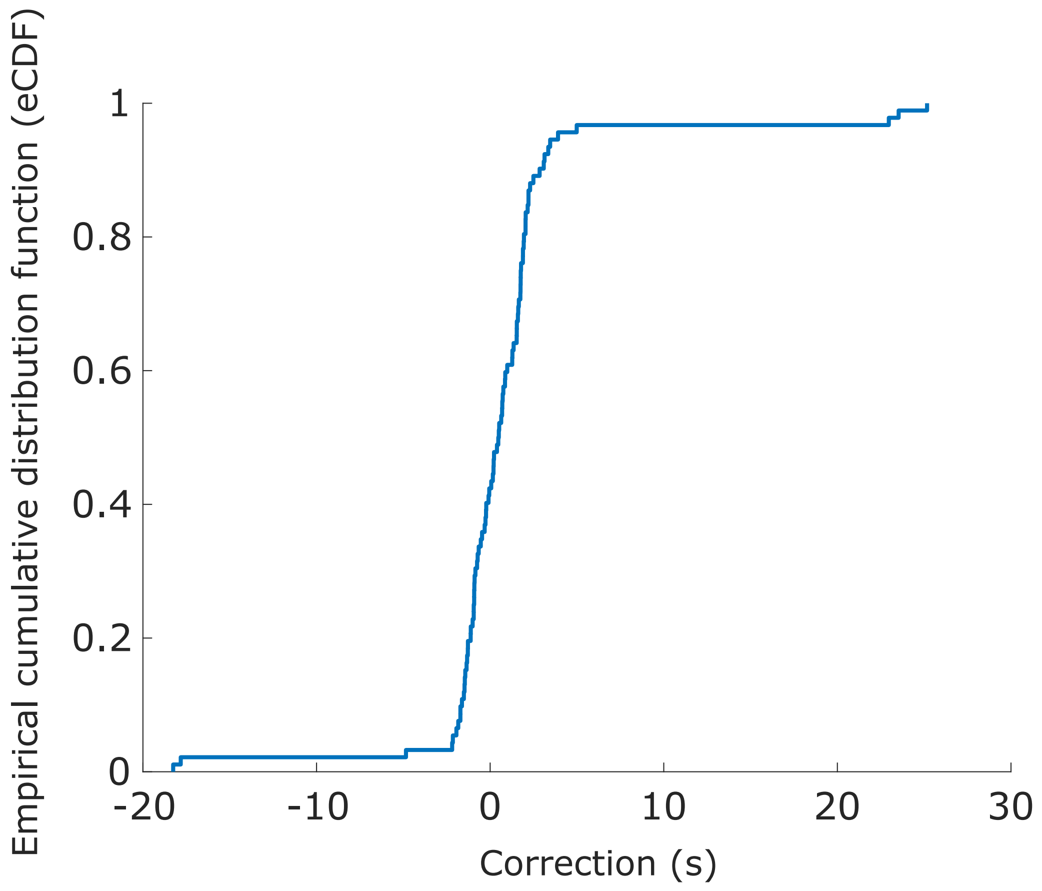


**Supplementary Methods Figure 4.** Alignment model correction. For each participant, the smartphone data and the movement data were aligned by correcting for the delay between the model predicted force and the touchscreen interaction.

**Supplementary Methods Table 2**

*Hyperparameters for artificial neural networks trained to identify goal and non-goal-directed movements*

| Parameter | Value |
| --- | --- |
| Train/Validation/Test split | 80/10/10 |
| Total number of trainable parameters | 66.689 |
| Optimizer | RMSprop |
| Starting learning rate | 0.0001 |
| Loss | Binary Cross Entropy |
| Max epoch | 100 |

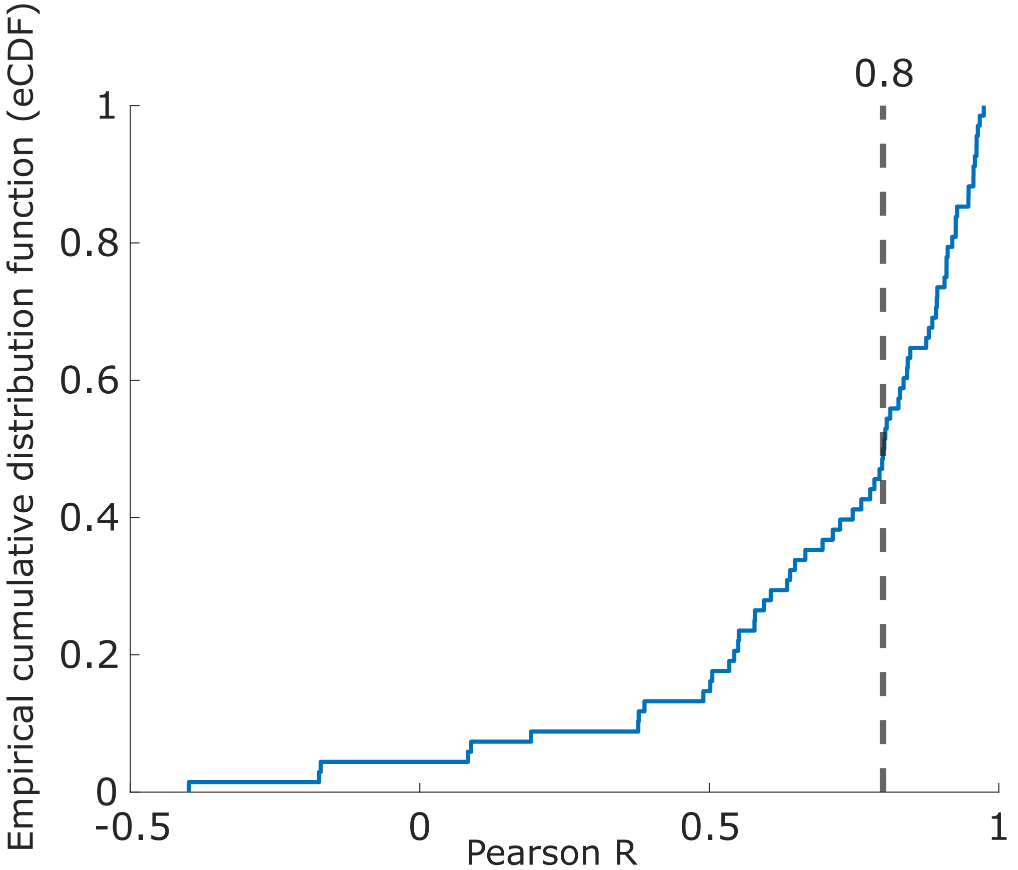


**Supplementary Methods Figure 5.** Participants with similar kinematic profiles were selected for EEG analysis based on a Pearson R larger than 0.8 (calculated over median z-normalized goal and non-goal-directed movements).

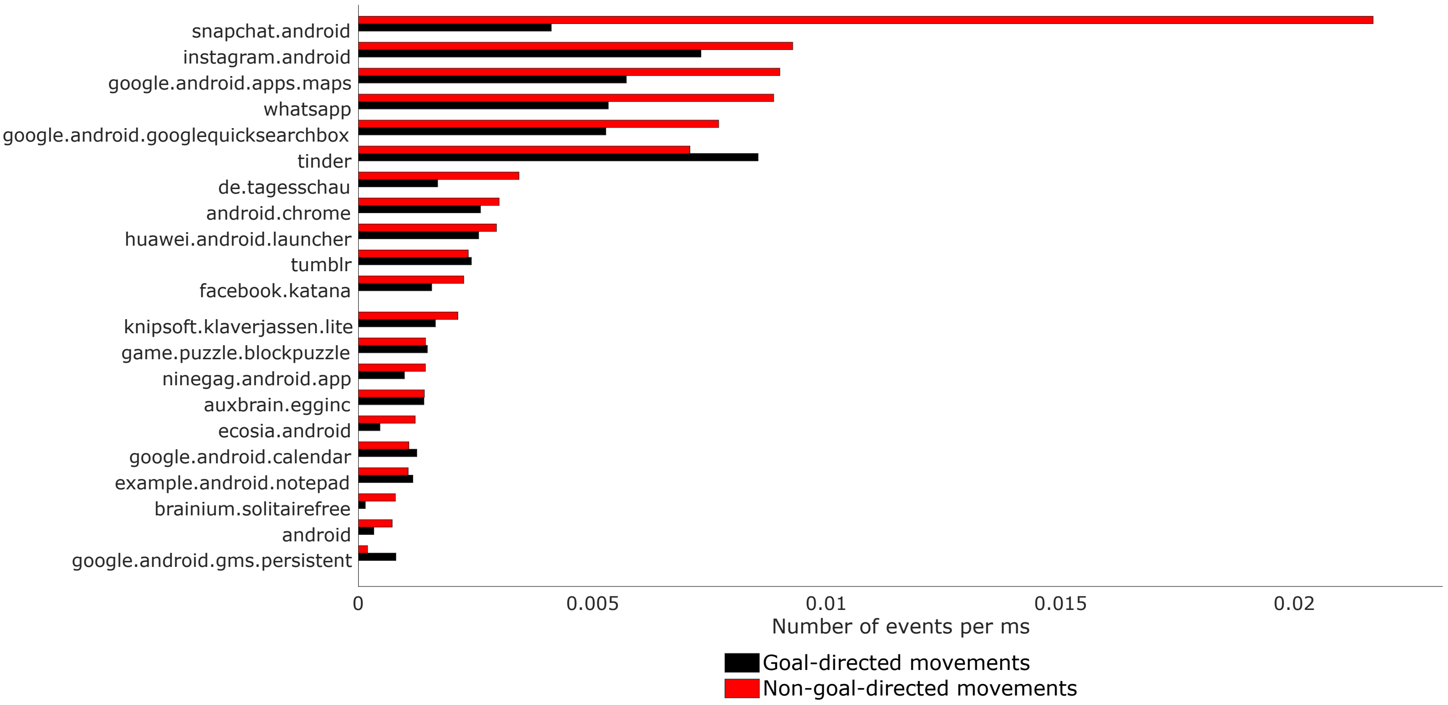


**Supplementary Methods Figure 6.** Frequency of goal-directed and non-goal-directed movements during use of smartphone apps
