## Supplementary Table 1 for "Neural processing of goal and non-goal-directed movements on the smartphone"

**Supplementary Table 1.** Descriptive statistics for artificial neural networks trained to identify the goal and non-goal-directed movements

| | **Median F2 score** | **Median Precision** | **Median Recall** |
|---|---|---|---|
| Selected participants (N = 32) | 0.3403 | 0.1419 | 0.5475 |
| All participants (N = 68) | 0.3453 | 0.1490 | 0.5484 |
